## Supplemental File for "Contemporary HIV-1 envelope pseudovirus panels for detecting and assessing B cell lineages with broadly neutralizing antibody potential"

#### Supplementary Figures

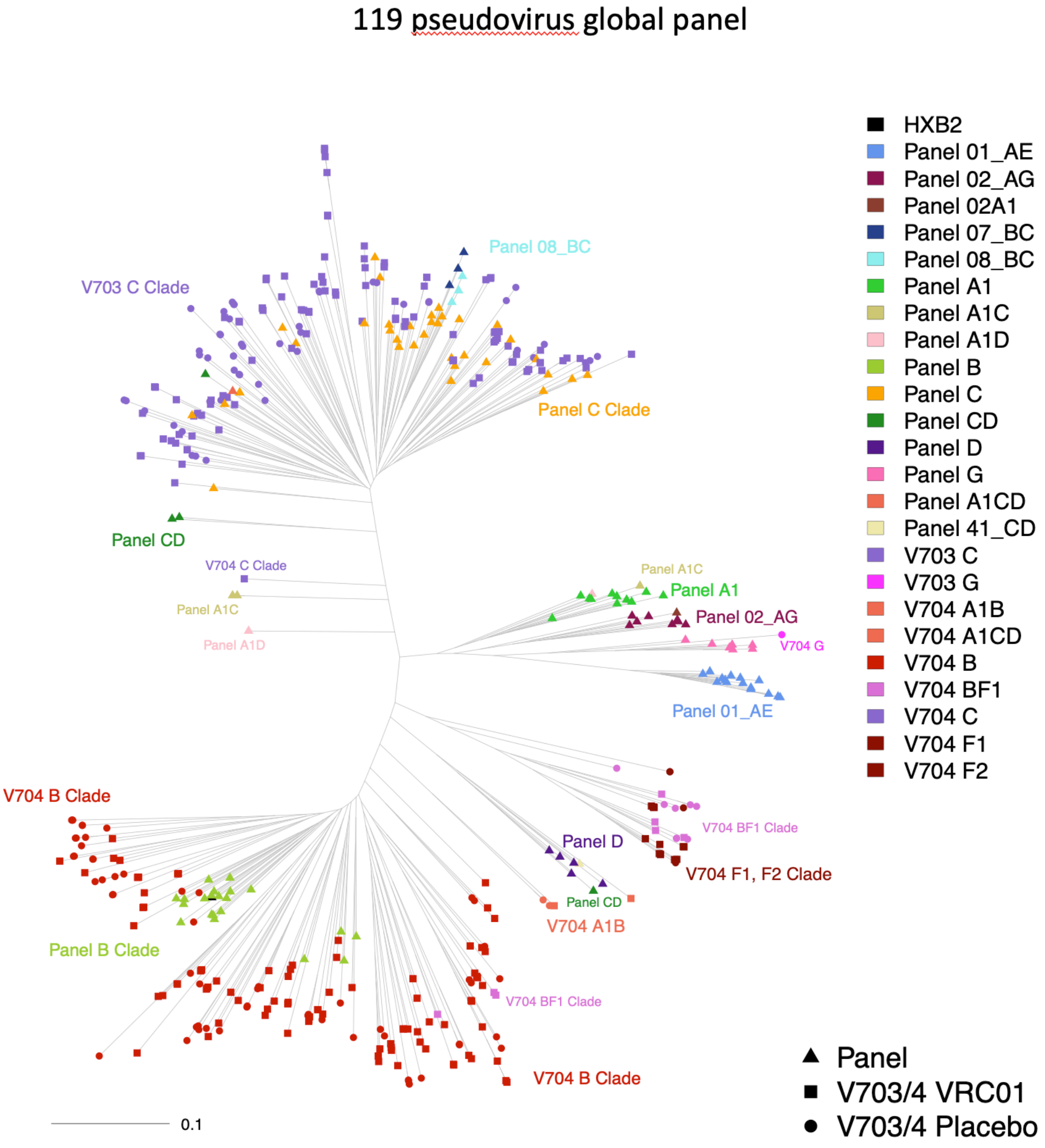

**Fig. S1. Midpoint rooted phylogenetic tree comparing a global panel of 119 viruses used to assess bnAb breadth and potency (1-4) to 154 AMP V703 and 191 V704 trial sequences (5, 6).** Phylogenetic analysis shows that within B and C clades the 119 viral panel sequences are not representative of the diversity sampled in the more contemporary AMP trial. Note that the exact composition of the panel can vary by a few viruses between studies. The phylogenetic tree was based on data from a version of the 119 virus panel that was downloaded using the CATNAP tool from the Los Alamos Database accessed on July 1, 2025 (7).

200 pseudovirus C clade panel

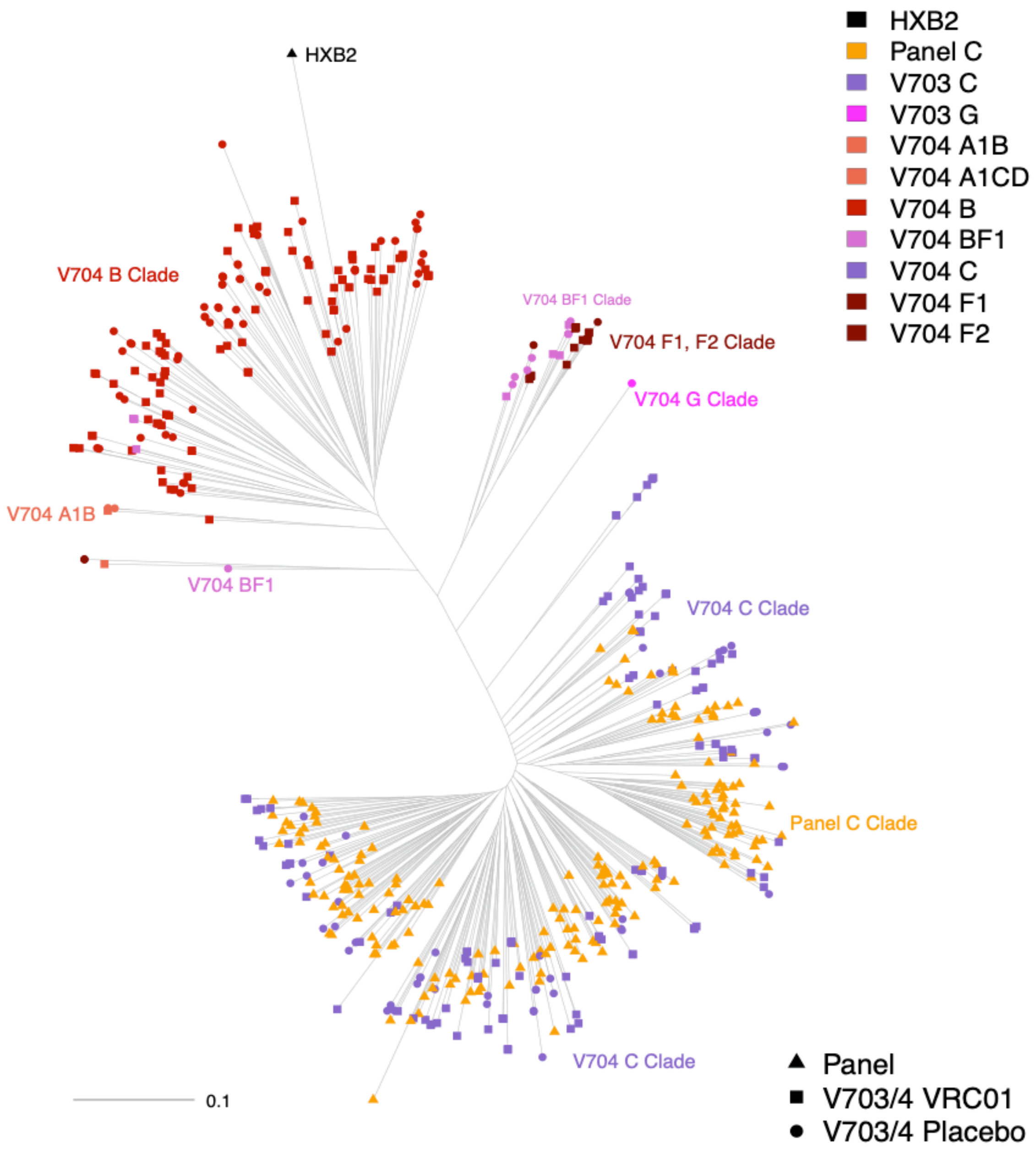

**Fig. S2. Midpoint rooted phylogenetic tree comparing a global panel of 200 clade C viruses used to assess bnAb breadth and potency (1-4) to 154 AMP V703 and 191 V704 trial sequences (5, 6)**. Of all commonly used panels described in this paper, this exclusively clade C panel, while lacking representation for all other clades, was the closest to contemporary C clade viruses sampled from the AMP V703 participants. Notably though, within the clade C subtree, AMP sequences tend to have longer branch lengths than the clade C panel pseudoviruses and are more spread out in the tree. The phylogenetic tree was based on data from a version of the 200 virus panel that was downloaded using the CATNAP tool from the Los Alamos Database accessed on July 1, 2025 (7).

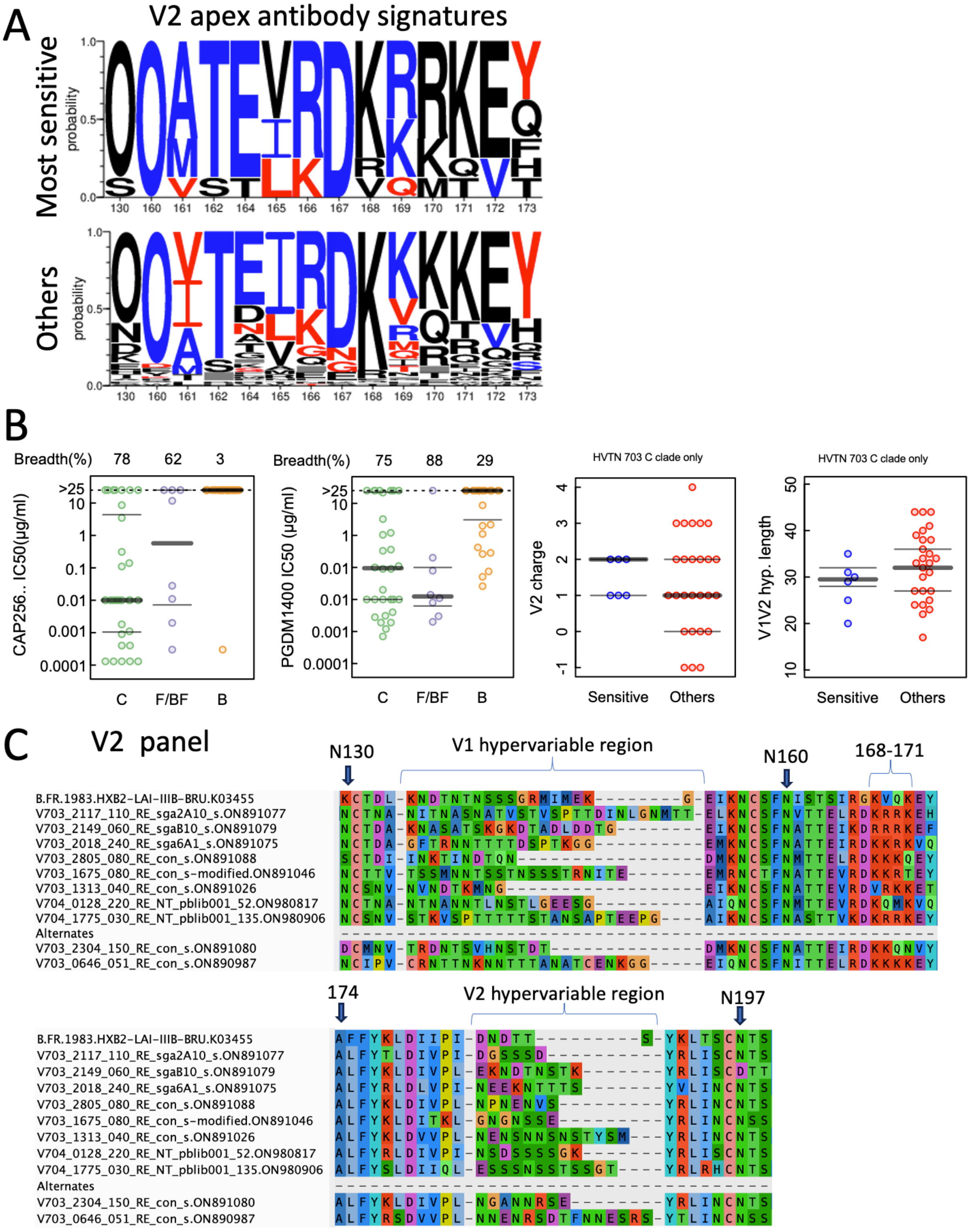

**Fig. S3. Details regarding V2 apex bNab signatures within the V2 apex bnAb sensitive panel for detection of developing responses. A.** The frequency of V2 apex bnAb sensitivity/resistance signatures as defined in Bricault et al. (8) in the class specific panel versus other AMP viruses. The height of the letter is indicative of the frequency of the amino acid in a given position in each group. An O indicates an N-linked glycosylation site. Blue are sensitivity signatures, red are resistance signatures, and black amino acids were not significantly associated with either one. **B.** On the left we illustrate (also see Table S5) the enhanced sensitivity of C clade and F/BF viruses relative to B clade viruses among the AMP collection of viruses to neutralization by the V2 apex bNab CAP256 VRC26.25 LS (far left) and PGDM1400 (second to the left). On the right we show the distribution of hypervariable region characteristics that were associated with V2 apex bNab neutralization (8). As B clade viruses tend to be resistant regardless of loop characteristics, we restricted these comparisons to AMP C clade viruses. Positive V2 charge and shorter combined V1V2 hypervariable region lengths are signatures of sensitivity to V2 Apex antibodies (8), and V2 hypervariable loop positive charge and shorter hypervariable regions are slightly enriched in the sensitive panel. **C.** The sequence alignment for the V2 apex bNab sensitive panel across the epitope region.

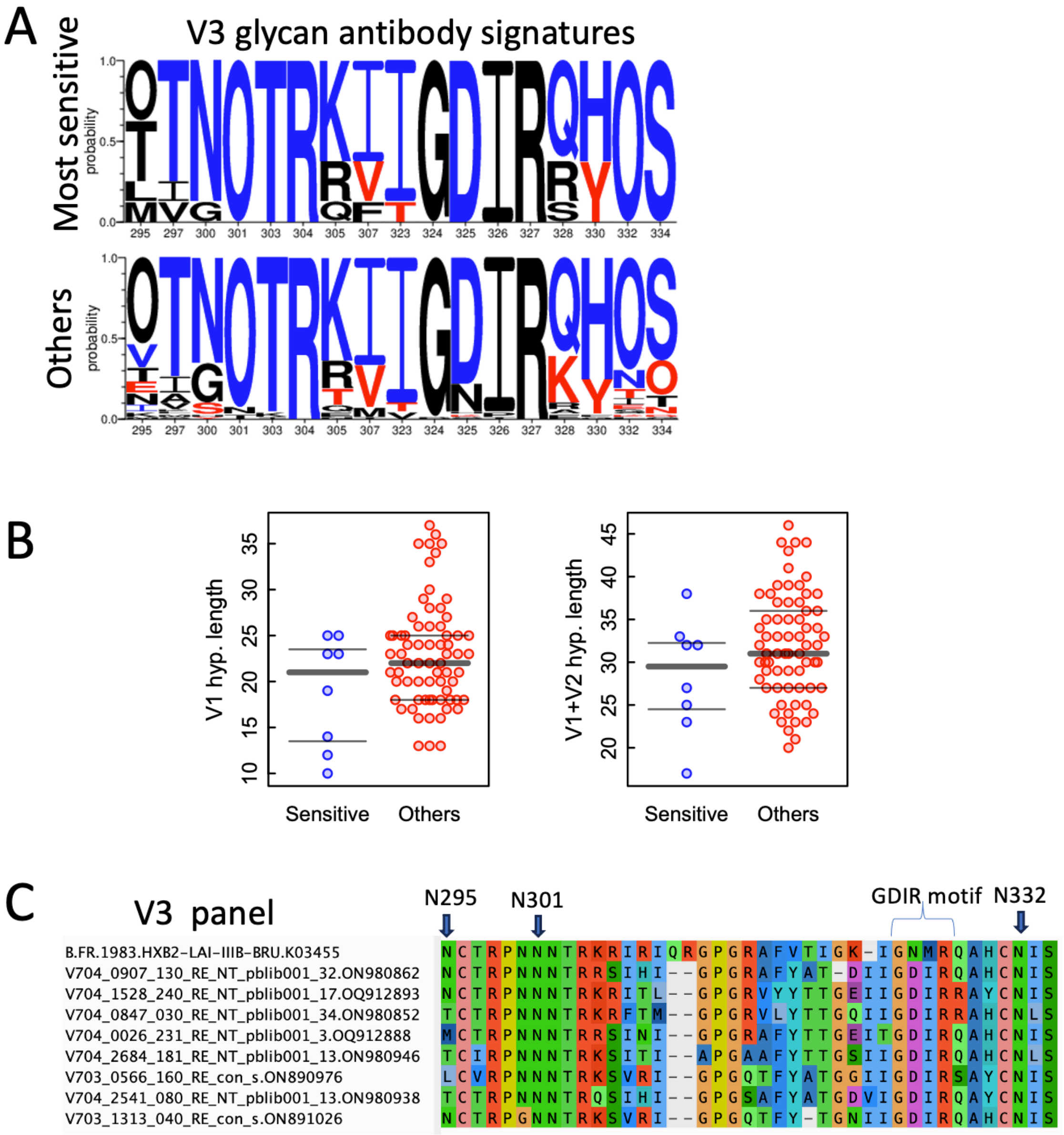

**Fig. S4. Details regarding V3 glycan bNab signatures within the V3 glycan bNab sensitive panel for detection of developing responses. A.** The frequency of V3 glycan bNab sensitivity/resistance signatures as defined in Bricault et al. (8) in the class specific panel versus other AMP viruses. The height of the letter is indicative of the frequency of the amino acid in a given position in each group. An O indicates an N-linked glycosylation site. Blue are sensitivity signatures, red are resistance signatures, and black amino acids were not significantly associated with either one. **B.** Shorter V1 hypervariable length and shorter combined V1V2 hypervariable region lengths are associated with enhanced sensitivity to V3 glycan bNabs (8), and there is an enrichment for shorter V1 and V1+V2 hypervariable regions in the V3 glycan sensitivity panel. **C.** The sequence alignment for the V3 glycan bNab sensitive panel across the epitope region.

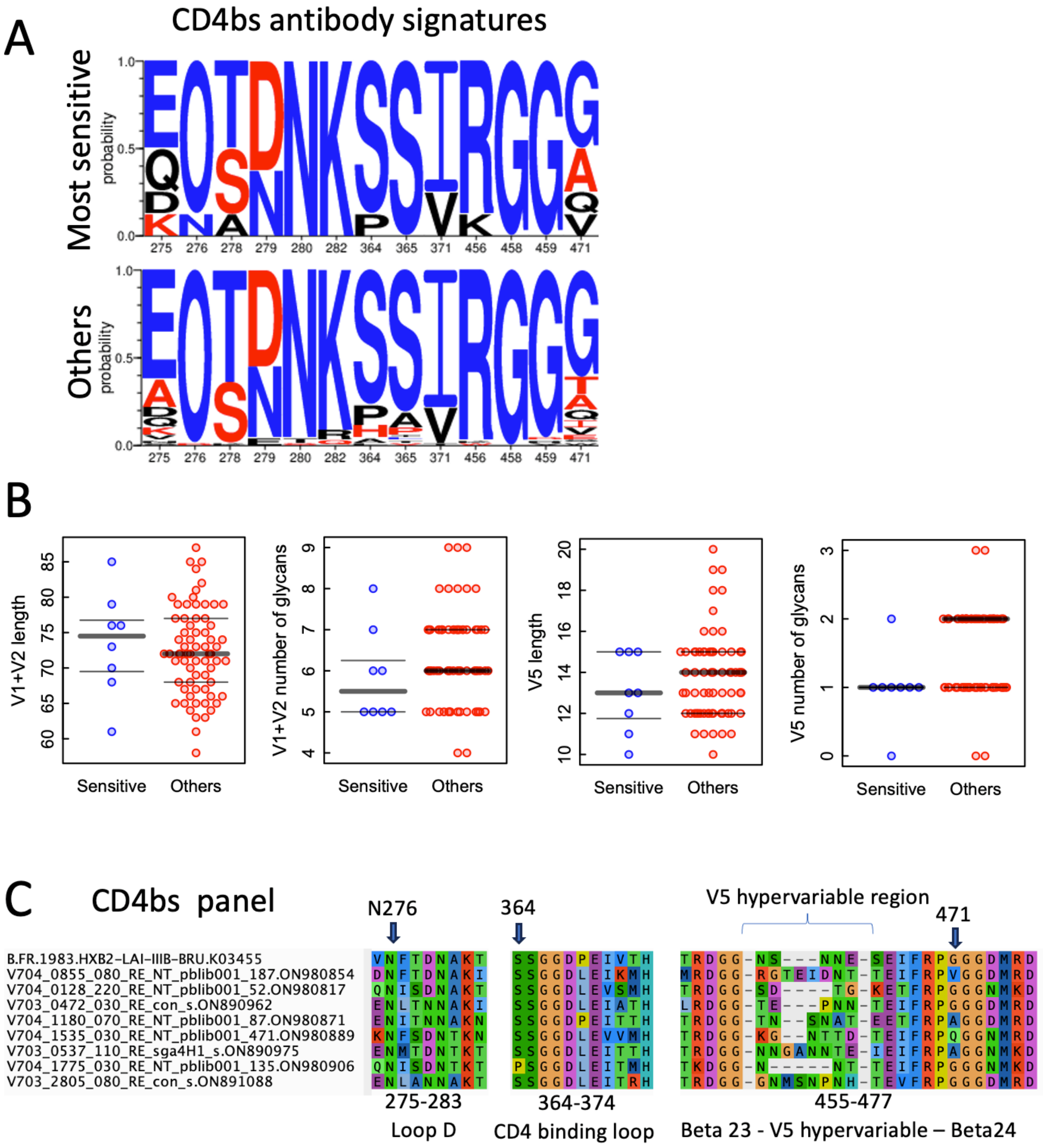

**Fig. S5. Details regarding CD4bs bNab signatures within the CD4 bNab sensitive panel for detection of developing responses. A.** The frequency of CD4bs bNab sensitivity/resistance signatures as defined in Bricault et al. (8) in the class specific panel versus other AMP viruses. The height of the letter is indicative of the frequency of the amino acid in a given position in each group. An O indicates an N-linked glycosylation site. Blue are sensitivity signatures, red are resistance signatures, and black amino acids were not significantly associated with either one. **B.** Shorter V1+V2 region lengths with fewer glycans were associated with CD4bs bNAb sensitivity (8), and while shorter combined V1V2 lengths were not enriched in the sensitive panel, fewer glycans in the V1V2 loop regions were. Shorter V5 loops with fewer glycans were also previously associated with enhanced CD4bs bNab sensitivity (8), and both of these characteristics were enriched in the sensitive panel. **C.** The sequence alignment for the CD4bs bNab sensitive panel across the epitope region.

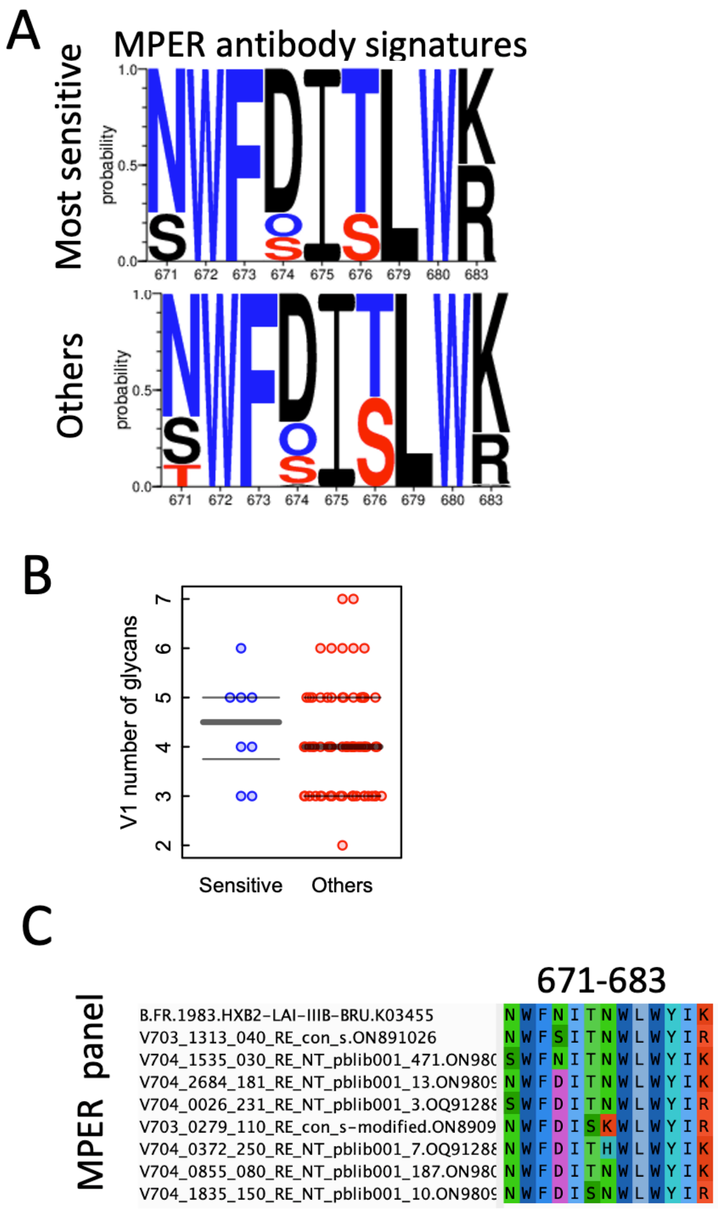

**Fig. S6. Details regarding MPER bNab signatures within the MPER sensitive panel for detection of developing responses. A.** The frequency of MPER sensitivity/resistance signatures as defined in Bricault et al. (8) in the class specific panel versus other AMP viruses. The height of the letter is indicative of the frequency of the amino acid in a given position in each group. An O indicates an N-linked glycosylation site. Blue are sensitivity signatures, red are resistance signatures, and black amino acids were not significantly associated with either one. **B.** Shorter V1 region lengths with fewer glycans were associated with MPER sensitivity (8), and these were not enriched in the MPER sensitive panel. **C.** The sequence alignment for the CD4bs bNab sensitive panel across the epitope region. This a highly conserved epitope.

### Supplementary Tables

### Table S1 A. AMP V703 IC50

| Sequence Name HVTN 703 | 1-18 | VRC07-523 LS | VRC01-23 LS | 3BNC117-LS | VRC01 | CH235.12 | PGT121-414 LS | 10-1074 LS | PGDM1400 | CAP256 VRC26.25 LS | PGT151 | VRC34.01 | ACS202 | 10E8v4 | CAP248-2B |
|---|---|---|---|---|---|---|---|---|---|---|---|---|---|---|---|
| V703_0013_090_RE_con_s | 0.52 | 0.4200 | 0.49 | >25 | 3.4725 | >10 | 7.9700 | 18.4100 | 0.0100 | 0.0100 | >10 | >10 | >10 | 1.6100 | >10 |
| V703_0203_081_RE_pblib002_s | 0.38 | 0.4500 | 1.23 | >25 | 3.8721 | 7.6 | 0.0400 | 0.0300 | >25 | >25 | >10 | >10 | >10 | 0.6700 | >10 |
| V703_0203_081_RE_sga8A5_s | 0.64 | 0.4100 | 0.91 | >25 | 4.1508 | >10 | 0.0300 | 0.0200 | >25 | >25 | >10 | >10 | >10 | 0.1300 | >10 |
| V703_0217_050_RE_pblib002_s | 0.0018 | 0.0700 | 0.06 | >25 | >25 | 0.15 | >25 | >25 | 0.0028 | 0.0011 | 0.0029 | 0.02 | 0.06 | 0.2400 | 0.01 |
| V703_0217_050_RE_sga2A3_s | 0.008 | 0.0300 | 0.04 | >25 | >25 | 0.29 | >25 | >25 | 0.0040 | 0.0020 | 0.01 | 0.0>253 | 2.21 | 0.1800 | 0.0247 |
| V703_0279_110_RE_con_s-modified | 0.08 | 0.2500 | 0.17 | 0.4400 | 1.3910 | 5.34 | 0.7700 | 0.4900 | 21.6100 | >25 | >10 | >10 | >10 | 0.0800 | >10 |
| V703_0279_110_RE_pblib002_s | 0.17 | 0.1800 | 0.14 | 0.3300 | 1.5089 | >10 | 0.9400 | 1.1900 | >25 | >25 | >10 | >10 | >10 | 0.3400 | >10 |
| V703_0472_030_RE_con_s | 0.0054 | 0.0100 | 0.005 | >25 | 0.0210 | 0.05 | >25 | 7.9400 | 0.1000 | 0.0100 | 0.028 | 5.67 | >10 | 0.4200 | >10 |
| V703_0537_110_RE_sga4H1_s | 0.01 | 0.0100 | 0.005 | 0.0200 | 0.0899 | 1.14 | 0.0100 | 0.0800 | 0.0900 | 0.0100 | >10 | >10 | >10 | 0.6800 | >10 |
| V703_0537_110_RE_sga2B5_s | 0.007 | 0.0300 | 0.005 | 0.0800 | 690443319 | 0.04 | 0.0200 | 0.1000 | 0.2200 | 0.0100 | >10 | >10 | >10 | 0.6600 | >10 |
| V703_0537_110_RE_pblib003_s | 0.0138 | 0.0200 | 0.005 | 0.0300 | 78>258823 | 0.19 | 0.0500 | 0.2100 | 1.0000 | 0.0100 | >10 | >10 | >10 | 0.2600 | >10 |
| V703_0566_160_RE_con_s | 0.04 | 0.0900 | 0.11 | 0.2000 | 1.3324 | 0.48 | 0.0100 | 0.0100 | >25 | >25 | 0.007 | 0.08 | >10 | 0.9200 | >10 |
| V703_0566_160_RE_pblib002_s | 0.06 | 0.1800 | 0.17 | 0.6600 | 3.5028 | 1.28 | 0.0300 | 0.0600 | >25 | >25 | 0.2 | 0.08 | >10 | 4.2100 | >10 |
| V703_0629_150_RE_pblib002_s | 0.08 | 0.0800 | 0.07 | 0.1100 | 1.4440 | 0.34 | 3.9000 | 3.1100 | 0.3600 | 0.1100 | 0.01 | 0.1 | >10 | 0.7400 | >10 |
| V703_0629_150_RE_sga5A2_s | 0.09 | 0.2200 | 0.0256 | 0.3900 | 0.7194 | 0.2 | 4.7200 | 4.4000 | 0.4200 | 0.0500 | 0.0032 | 0.16 | >10 | 4.2700 | >10 |
| V703_0629_150_RE_pblib003_s | 0.11 | 0.3300 | 0.06 | 0.7200 | 1.0759 | 1.5 | 9.8700 | 5.6100 | 1.5100 | >25 | 0.06 | >10 | >10 | 6.4400 | >10 |
| V703_0629_150_RE_pblib004_s | 0.12 | 0.2400 | 0.08 | 0.8000 | 1.0282 | 0.41 | 6.4100 | 4.1200 | 0.1100 | 0.2200 | 0.01 | 1.1 | >10 | 2.4600 | >10 |
| V703_0646_051_RE_con_s | 0.18 | 0.4500 | 0.63 | 1.5400 | 4.8972 | >10 | >25 | >25 | 0.0100 | 0.0100 | 0.07 | 0.07 | >10 | 1.5600 | >10 |
| V703_0739_110_RE_con_s-modified | 0.025 | 0.0500 | 0.023 | 0.0700 | 0.1431 | 0.56 | 0.6700 | 0.1600 | 3.2300 | 0.0009 | >10 | 4.86 | 0.09 | 1.9400 | >10 |
| V703_0842_200_RE_con_s | >10 | 1.4400 | 1.82 | 2.1300 | 20.8948 | 6.36 | 7.2300 | 1.5600 | 0.1100 | 0.0100 | 0.21 | >10 | >10 | 0.7300 | >10 |
| V703_0865_070_RE_con_s | 0.09 | 0.0900 | ND | ND | ND | 12 | ND | ND | ND | ND | >10 | >10 | >10 | ND | ND |
| V703_0926_070_RE_sga2H2_s | 2.02 | 0.8100 | 0.76 | 2.0400 | 7.8027 | 2.34 | 0.0400 | 0.1000 | 1.0400 | >25 | >10 | >10 | >10 | 0.5600 | >10 |
| V703_1060_080_RE_con_s | 0.09 | 0.1300 | 0.28 | 0.8000 | 1.7123 | 1.17 | >25 | 3.6300 | >25 | 0.3100 | 0.05 | >10 | >10 | 0.0800 | >10 |
| V703_1104_100_RE_pblib001_s | 0.18 | 1.8100 | 0.92 | 2.5700 | 4.0481 | >10 | >25 | 2.9800 | 0.3300 | 0.1400 | >10 | >10 | 0.19 | 0.9200 | >10 |
| V703_1104_100_RE_sga10A5_s | 0.33 | 1.3200 | 1.69 | 1.8000 | 8.0808 | >10 | >25 | 3.5800 | 0.9200 | 0.1200 | >10 | >10 | 0.15 | 0.2800 | >10 |
| V703_1104_100_RE_sga10G5_s | 0.28 | 1.6100 | 1.41 | 2.9900 | 6.4095 | >10 | >25 | 10.4800 | 0.4700 | 0.1200 | >10 | >10 | 4.23 | 0.8600 | >10 |
| V703_1298_080_RE_pblib002_s | 0.15 | 1.2700 | 0.13 | >25 | >25 | 0.16 | >25 | >25 | 0.0900 | 0.0100 | 0.1 | >10 | >10 | 0.8400 | >10 |
| V703_1313_040_RE_con_s | 0.046 | 0.0400 | 0.07 | 0.1100 | 0.6860 | >10 | 0.0100 | 0.0600 | 0.0040 | 0.0018 | >10 | >10 | >10 | 0.0100 | >10 |
| V703_1471_190_RE_con_s | 0.05 | 0.0800 | 0.0235 | 0.0600 | 782805>25 | 0.6 | 0.0100 | 0.1000 | 0.0200 | 0.0100 | 0.01 | >10 | >10 | 4.9200 | 0.011 |
| V703_1675_080_RE_con_s-modified | 0.02 | 0.0800 | 0.0414 | 0.0600 | 0.3446 | 0.15 | 0.9800 | 1.4200 | 0.0100 | 0.0001 | 0.59 | >10 | 4.47 | 18.5300 | >10 |
| V703_1675_080_RE_con_s | 0.0279 | 0.0500 | 0.0528 | 0.0400 | 9183384>2 | 0.23 | 0.2600 | 0.7900 | 0.0100 | 0.0100 | >10 | >10 | 1.49 | 14.6600 | >10 |
| V703_1750_140_RE_con_s | 0.11 | 0.1800 | 0.13 | 0.2300 | 0.8813 | 1.35 | 0.0500 | 0.0800 | >25 | 8.5900 | 0.05 | 0.34 | >10 | 0.5000 | >10 |
| V703_1764_250_RE_con_s | 0.03 | 0.1000 | 0.06 | 0.1000 | 0.2402 | 0.1 | 0.2000 | 0.1300 | >25 | >25 | 0.18 | >10 | >10 | 1.4100 | >10 |
| V703_1789_230_RE_sga3A3_s | 0.09 | 0.9900 | 0.55 | >25 | 17.8963 | >10 | 0.8300 | 0.3400 | >25 | >25 | >10 | >10 | >10 | 2.4300 | >10 |
| V703_1848_190_RE_pblib002_s | 0.0517 | 0.1500 | 0.07 | 0.0600 | 1.0367 | 0.92 | >25 | 6.7400 | 0.0100 | 0.0100 | 0.06 | >10 | >10 | 0.2800 | >10 |
| V703_1848_190_RE_sga6D1_s | 0.09 | 0.3100 | 0.2 | 0.2400 | 1.0544 | 0.24 | >25 | >25 | 11.4600 | >25 | 0.02 | >10 | >10 | 2.7600 | >10 |
| V703_1889_100_RE_con_s | 0.09 | 2.3000 | 0.19 | >25 | >25 | 5.78 | 0.0900 | 0.2300 | 0.0100 | 0.0001 | >10 | >10 | >10 | >25 | >10 |
| V703_1915_250_RE_sgaA3_s | 0.0841 | 0.3600 | 0.0406 | 1.6000 | 0.5291 | 5.6 | 0.3200 | 0.8400 | >25 | 3.4800 | 0.04 | >10 | >10 | 3.8300 | >10 |
| V703_2018_240_RE_sga6A1_s | 0.16 | 0.3000 | 0.09 | 1.1300 | 1.2489 | 6.38 | 0.0500 | 0.0700 | 0.0031 | 0.0001 | >10 | >10 | >10 | 2.8500 | >10 |
| V703_2117_110_RE_sga2A10_s | 0.17 | 0.0400 | 0.06 | 0.1100 | 0.2285 | 0.52 | >25 | >25 | 0.0007 | 0.0001 | >10 | 0.81 | >10 | 0.8800 | >10 |
| V703_2149_060_RE_sgaB10_s | 0.0324 | 0.0700 | 0.0208 | >25 | 0.2050 | >10 | 0.1100 | 0.0700 | 0.0012 | 0.0001 | 9.17 | >10 | >10 | 0.9800 | >10 |
| V703_2304_150_RE_con_s | 0.06 | 0.0200 | 0.0106 | 0.0600 | 0.1135 | 0.08 | 0.0700 | 0.0600 | 0.0100 | 0.0100 | >10 | 0.03 | >10 | 0.3600 | >10 |
| V703_2539_070_RE_sga6F6_s | 0.74 | 0.3900 | 0.27 | >25 | 1.7655 | >10 | 0.0100 | 0.0400 | >25 | >25 | 4.57 | >10 | >10 | 0.2300 | >10 |
| V703_2631_150_RE_sga2F8_s | 0.02 | 0.0800 | 0.09 | 0.0700 | 0.4631 | 1.02 | >25 | >25 | 0.0019 | 0.0004 | >10 | 0.97 | >10 | 1.2300 | >10 |
| V703_2788_030_RE_con_s | 1.56 | 0.2400 | 4.11 | 0.0900 | 0.6812 | 4.9 | 0.3000 | 0.0900 | 1.2000 | 0.0100 | 1.84 | >10 | >10 | 5.3800 | >10 |
| V703_2805_080_RE_con_s | 0.027 | 0.0100 | 0.0287 | 0.0200 | 0.2668 | 0.24 | >25 | >25 | 0.0019 | 0.0004 | 0.06 | 0.23 | >10 | 1.2200 | >10 |
| V703_2805_080_RE_pblib002_s | >10 | 1.8700 | >10 | 5.1700 | >25 | >10 | >25 | >25 | >25 | 5.9000 | >10 | 0.84 | >10 | 2.3800 | >10 |

Sequences from the same person are grouped.
Gray: Most sensitive TFL virus
Yellow: Most resistant TFL virus

| IC50/IC80 |
|---|
| HVTN 703: >10 |
| HVTN 704: >25 |
| HVTN 704: >10 - 25 |
| >1 - 10 |
| >0.1 - 1 |
| >0.01 - 0.1 |
| >0.001 - 0.01 |
| <= 0.001 |

### Table S1 B. AMP V704 IC50

| Sequence Name HVTN 704 | IC50 | | | | | | | | | | | | | | |
|---|---|---|---|---|---|---|---|---|---|---|---|---|---|---|---|
| | CD4bs | | | | | | V3 | | V2 | | FP | | | MPER | |
| | 1-18 | VRC07-523 LS | VRC01-23 LS | 3BNC117-LS | VRC01 | CH235.12 | PGT121-414 LS | 10-1074 LS | PGDM1400 | CAP256 VRC26.25 LS | PGT151 | VRC34.01 | ACS202 | 10E8v4 | CAP248-2B |
| V704_0026_231_RE_NT_pblib001_3 | 0.097 | 0.081 | 0.088 | 0.103 | 0.46 | >25 | 0.006 | 0.017 | 0.075 | >25 | >25 | >25 | >25 | 0.07 | >25 |
| V704_0128_220_RE_NT_pblib001_52 | 0.023 | 0.006 | 0.011 | 0.026 | 0.04 | 0.021 | 0.018 | 0.059 | 0.008 | 0.002 | >25 | 0.194 | 22.918 | 1.221 | ND |
| V704_0128_220_RE_NT_pblib002_40 | 0.044 | 0.026 | 0.034 | 0.025 | 0.16 | 0.089 | 0.047 | 0.171 | 0.031 | 0.0003 | >25 | 2.044 | >25 | 1.33 | >25 |
| V704_0445_180_RE_NT_pblib001_22 | 0.012 | 0.063 | 0.021 | >25 | 82.985 | 0.205 | 0.291 | 0.086 | 0.285 | >25 | 0.083 | >25 | 0.08 | 1.49 | >25 |
| V704_0496_040_RE_NT_pblib001_37 | 0.037 | 0.242 | 0.601 | >25 | 4.940 | 1.236 | 0.142 | 0.078 | 0.011 | 0.011 | >25 | >25 | >25 | 0.289 | >25 |
| V704_0513_150_RE_NT_pblib007_2 | 0.413 | 0.263 | 0.354 | 0.29 | 1.08 | 17.817 | >25 | >25 | >25 | >25 | 0.047 | 1.468 | >25 | 0.98 | >25 |
| V704_0513_150_RE_NT_pblib002_10 | 0.465 | 0.401 | 0.387 | 0.251 | 1.445 | 5.793 | >25 | >25 | >25 | >25 | 0.044 | >25 | >25 | 1.049 | >25 |
| V704_0513_150_RE_NT_pblib001_34 | 0.612 | 0.38 | 0.329 | 0.276 | 1.49 | 6.07 | >25 | >25 | >25 | >25 | 0.49 | >25 | >25 | 1.13 | >25 |
| V704_0575_060_RE_NT_pblib002_16 | 0.244 | 0.989 | 0.615 | 1.294 | 1.934 | 2.286 | >25 | >25 | 0.051 | >25 | >25 | 0.239 | 0.039 | 0.57 | >25 |
| V704_0575_060_RE_NT_pblib001_60 | 0.412 | 1.02 | 0.652 | 1.241 | 2.195 | 2.275 | >25 | >25 | 0.047 | >25 | >25 | 0.657 | 0.083 | 0.417 | >25 |
| V704_0575_060_RE_NT_con02_s | 0.467 | 0.756 | 0.771 | 0.965 | 2.118 | 2.354 | >25 | >25 | 0.042 | >25 | >25 | >25 | 0.157 | 0.494 | >25 |
| V704_0644_060_RE_NT_pblib001_216 | 0.084 | 0.155 | 0.279 | 0.29 | 1.02 | 3.275 | 6.348 | 0.454 | >25 | >25 | >25 | 5.728 | >25 | 0.297 | >25 |
| V704_0726_080_RE_NT_pblib001_26 | 0.199 | 1.020 | 0.466 | 0.537 | 5.636 | 3.445 | 0.36 | 0.734 | 0.421 | >25 | >25 | >25 | >25 | 2.473 | >25 |
| V704_0746_760_RE_NT_pblib001_37 | 0.19 | 0.397 | 0.15 | 0.255 | 0.87 | 5.201 | 0.015 | 0.014 | >25 | >25 | >25 | >25 | >25 | 3.08 | >25 |
| V704_0746_760_RE_NT_pblib003_34 | 0.187 | 0.415 | 0.122 | 0.26 | 1.07 | 7.051 | 0.018 | 0.013 | >25 | >25 | >25 | >25 | >25 | 2.56 | >25 |
| V704_0746_760_RE_NT_pblib002_36 | 0.224 | 0.447 | 0.189 | 0.282 | 1.25 | 6.796 | 0.02 | 0.015 | >25 | >25 | >25 | >25 | >25 | 1.96 | >25 |
| V704_0847_030_RE_NT_pblib001_34 | 6.634 | 0.19 | 0.053 | >25 | 2.048 | >25 | 0.01 | 0.003 | >25 | >25 | >25 | >25 | >25 | 0.221 | 0.041 |
| V704_0847_030_RE_NT_pblib002_23 | 9.694 | 0.204 | 0.161 | >25 | 3.118 | >25 | 0.011 | 0.006 | >25 | >25 | >25 | >25 | >25 | 0.27 | 1.448 |
| V704_0855_080_RE_NT_pblib001_187 | 0.012 | 0.003 | 0.002 | 0.009 | 0.013 | >25 | 0.05 | 0.061 | >25 | >25 | 0.007 | >25 | >25 | 0.134 | >25 |
| V704_0856_240_RE_NT_pblib001_9 | 0.095 | 0.063 | 0.055 | 0.06 | 0.524 | 0.503 | 0.011 | 0.016 | >25 | >25 | 1.04 | 7.845 | >25 | 0.54 | >25 |
| V704_0886_250_RE_NT_pblib001_10 | 0.173 | 0.177 | 0.099 | 0.516 | 1.745 | >25 | 0.338 | 0.185 | >25 | >25 | >25 | >25 | 0.498 | 0.171 | >25 |
| V704_0907_130_RE_NT_pblib001_32 | 0.016 | 0.193 | 0.019 | 0.199 | 0.364 | 15.857 | 0.002 | 0.006 | >25 | >25 | 0.017 | 0.25 | 0.236 | 0.655 | >25 |
| V704_0911_150_RE_NT_pblib001_58 | 0.437 | 0.538 | 1.525 | 0.622 | 1.391 | 10.111 | 0.013 | 0.018 | >25 | >25 | 3.99 | >25 | >25 | 0.869 | >25 |
| V704_0944_180_RE_NT_pblib001_76 | 0.023 | 0.02 | 0.007 | 0.061 | 0.074 | >25 | >25 | >25 | >25 | >25 | 0.367 | >25 | >25 | >25 | >25 |
| V704_1109_140_RE_NT_pblib001_191 | >25 | 0.283 | 2.455 | >25 | 3.36 | >25 | >25 | >25 | 0.201 | >25 | >25 | >25 | >25 | 1.25 | >25 |
| V704_1180_070_RE_NT_pblib001_87 | 0.005 | 0.045 | 0.013 | 0.016 | 0.17 | 0.145 | >25 | >25 | 0.188 | >25 | 0.009 | 0.283 | 0.273 | 0.32 | 0.114 |
| V704_1180_070_RE_NT_pblib027_1 | 0.005 | 0.074 | 0.009 | 0.018 | 0.137 | 0.419 | >25 | >25 | 0.405 | >25 | 0.009 | 0.683 | 0.388 | 0.16 | >25 |
| V704_1183_220_RE_NT_pblib001_70 | 0.041 | 0.347 | 0.038 | 0.065 | 0.39 | 1.185 | 0.032 | 0.051 | >25 | >25 | 3.666 | >25 | 0.228 | 0.495 | >25 |
| V704_1429_090_RE_NT_pblib002_75 | 0.111 | 0.097 | 0.078 | 1.216 | 0.235 | >25 | >25 | >25 | 0.079 | 11.679 | 4.01 | >25 | >25 | 0.537 | >25 |
| V704_1429_090_RE_NT_pblib001_111 | 0.119 | 0.112 | 0.122 | 1.806 | 0.307 | >25 | >25 | >25 | 0.141 | >25 | 4.52 | >25 | >25 | 0.45 | >25 |
| V704_1528_240_RE_NT_pblib001_17 | 0.06 | 0.439 | 0.114 | 0.092 | 2.39 | 7.107 | 0.003 | 0.004 | >25 | >25 | 1.418 | 0.04 | 0.041 | 0.243 | 1.15 |
| V704_1535_030_RE_NT_pblib001_471 | 0.052 | 0.068 | 0.018 | 0.019 | 0.088 | >25 | 0.011 | 0.071 | 1.109 | >25 | 0.197 | 0.391 | 0.512 | 0.015 | 0.334 |
| V704_1706_040_RE_NT_pblib001_26 | 0.2 | 0.058 | 0.058 | 0.164 | 0.489 | 0.186 | 0.457 | 0.301 | 1.986 | >25 | >25 | >25 | >25 | 0.488 | >25 |
| V704_1775_030_RE_NT_pblib001_135 | 0.014 | 0.029 | 0.031 | 0.109 | 0.09 | 0.05 | >25 | >25 | 0.002 | 0.028 | 0.112 | >25 | >25 | 0.291 | >25 |
| V704_1783_150_RE_NT_pblib001_144 | 0.006 | 0.064 | 0.009 | 0.036 | 0.135 | >25 | 0.248 | 0.485 | >25 | >25 | 0.06 | >25 | 0.353 | 0.388 | >25 |
| V704_1835_150_RE_NT_pblib001_10 | 0.027 | 21.596 | 0.041 | >25 | 99 | >25 | 3.127 | 0.095 | >25 | >25 | 0.112 | 0.412 | 0.113 | 0.147 | >25 |
| V704_1835_150_RE_NT_pblib002_9 | 0.026 | 24.845 | 0.078 | >25 | >100 | >25 | 3.791 | 0.081 | >25 | >25 | 0.24 | 1.729 | 0.397 | 0.099 | >25 |
| V704_1930_170_RE_NT_pblib002_114 | 0.032 | 0.025 | 0.013 | 0.028 | 0.11 | 0.087 | >25 | >25 | 0.014 | >25 | >25 | >25 | >25 | 1 | >25 |
| V704_1930_170_RE_NT_pblib001_234 | 0.041 | 0.035 | 0.023 | 0.043 | 0.134 | 0.118 | >25 | >25 | 0.018 | >25 | >25 | >25 | >25 | 1.25 | >25 |
| V704_1991_230_RE_NT_pblib001_38 | 0.093 | 0.284 | 0.171 | 0.242 | 1.51 | 0.303 | >25 | >25 | 0.026 | 0.0003 | 0.373 | >25 | 0.128 | 0.5 | >25 |
| V704_1991_230_RE_NT_pblib002_19 | 0.067 | 0.215 | 0.14 | 0.165 | 1.488 | 0.309 | >25 | >25 | 0.052 | 0.0003 | >25 | >25 | 0.12 | 0.317 | >25 |
| V704_2065_060_RE_NT_pblib001_47 | 0.069 | 0.146 | 0.028 | 0.047 | 0.22 | 0.052 | >25 | >25 | >25 | >25 | >25 | >25 | >25 | 0.1 | >25 |
| V704_2065_060_RE_NT_pblib002_42 | 0.062 | 0.082 | 0.037 | 0.048 | 0.22 | 0.036 | >25 | >25 | >25 | >25 | >25 | >25 | >25 | 0.161 | >25 |
| V704_2095_130_RE_NT_pblib001_136 | 0.079 | 0.096 | 0.031 | 0.053 | 0.312 | 0.283 | 2.167 | 0.085 | >25 | >25 | 0.147 | >25 | >25 | 0.51 | >25 |
| V704_2448_240_RE_NT_pblib001_288 | 0.133 | 0.108 | 0.01 | 0.058 | 0.14 | 0.146 | >25 | >25 | >25 | >25 | >25 | >25 | 7.416 | >25 | >25 |
| V704_2541_080_RE_NT_pblib001_13 | 0.037 | 0.056 | 0.033 | 0.032 | 0.305 | 0.288 | 0.026 | 0.008 | 2.163 | >25 | 0.033 | 0.755 | >25 | 0.25 | >25 |
| V704_2541_080_RE_NT_pblib042_1 | 0.026 | 0.036 | 0.018 | 0.021 | 0.21 | 0.112 | 0.022 | 0.027 | 0.218 | >25 | 0.168 | 2.872 | >25 | 0.08 | >25 |
| V704_2541_080_RE_NT_pblib003_5 | 0.053 | 0.069 | 0.025 | 0.051 | 0.38 | 0.409 | 0.044 | 0.011 | 3.382 | >25 | 0.048 | 0.477 | >25 | 0.184 | >25 |
| V704_2544_140_RE_NT_pblib002_10 | 0.431 | 0.122 | 0.136 | 0.108 | 0.922 | 4.104 | 0.493 | 0.611 | >25 | >25 | >25 | >25 | >25 | 0.4 | >25 |
| V704_2544_140_RE_NT_pblib001_11 | 0.99 | 0.431 | 0.494 | 0.82 | 5.76 | 8.648 | 16.44 | 0.421 | 13.864 | >25 | 0.059 | >25 | >25 | 1.127 | 17.994 |
| V704_2544_150_RE_NT_pblib001_33 | 0.262 | 0.2 | 0.18 | 0.35 | 1.84 | 3.257 | 0.03 | 0.063 | 0.101 | >25 | >25 | >25 | >25 | 0.29 | >25 |
| V704_2555_240_RE_NT_pblib001_231 | 0.092 | 0.052 | 0.033 | 0.075 | 0.424 | >25 | 0.012 | 0.03 | >25 | >25 | >25 | 2.808 | 0.151 | >25 | >25 |
| V704_2684_181_RE_NT_pblib001_13 | 0.01 | 0.044 | 0.014 | 0.051 | 0.09 | >25 | 0.006 | 0.019 | >25 | >25 | >25 | >25 | 0.913 | 0.03 | >25 |
| V704_2684_181_RE_NT_pblib006_3 | 0.007 | 0.057 | 0.012 | 0.43 | 0.092 | >25 | 0.007 | 0.019 | >25 | >25 | >25 | >25 | 0.97 | 0.031 | >25 |
| V704_2767_070_RE_NT_pblib001_63 | 0.217 | 0.126 | 0.068 | 0.796 | 2.06 | 1.275 | 0.066 | 0.241 | >25 | >25 | 0.038 | >25 | >25 | 2.03 | >25 |
| V704_2788_060_RE_NT_pblib001_96 | 0.075 | 0.173 | 0.277 | 0.167 | 1.36 | 0.694 | 0.171 | 0.201 | 0.003 | 0.0003 | 2.78 | >25 | >25 | 0.43 | >25 |
| V704_2788_060_RE_NT_pblib002_20 | 0.192 | 0.260 | 0.284 | 0.223 | 1.409 | 1.144 | 0.134 | 0.144 | 0.011 | 0.011 | 16.77 | >25 | >25 | 0.082 | >25 |
| V704_2834_210_RE_NT_pblib002_26 | 0.056 | 0.199 | 0.088 | 0.059 | 2.039 | >25 | 7.78 | 3.385 | >25 | >25 | >25 | 1.179 | >25 | 0.397 | >25 |
| V704_2834_210_RE_NT_pblib001_44 | 0.107 | 0.312 | 0.073 | 0.086 | 2.19 | >25 | 19.219 | 4.297 | >25 | >25 | >25 | 0.922 | >25 | 0.12 | >25 |
| V704_2839_140_RE_NT_pblib001_208 | 0.116 | 0.183 | 0.346 | 1.426 | 1.58 | >25 | >25 | >25 | >25 | >25 | >25 | 1.771 | 1.486 | 0.63 | >25 |
| V704_2981_150_RE_NT_pblib004_2 | 0.392 | 0.414 | 0.102 | 0.129 | 2.43 | 0.279 | 1.271 | 0.702 | 0.262 | >25 | >25 | 9.095 | >25 | 0.23 | >25 |
| V704_2981_150_RE_NT_pblib001_33 | 5.259 | 1.404 | 0.942 | 1.038 | 6.07 | 2.665 | 0.609 | 0.421 | 0.022 | >25 | >25 | 24.65 | >25 | 2.89 | >25 |
| V704_2981_150_RE_NT_pblib002_11 | 0.809 | 0.443 | 0.304 | 0.248 | 1.89 | 0.734 | >25 | >25 | >25 | >25 | >25 | 16.439 | >25 | 1.09 | >25 |
| V704_2981_150_RE_NT_pblib003_4 | 1.058 | 0.39 | 0.178 | 0.155 | 1.84 | 0.333 | >25 | >25 | >25 | >25 | >25 | >25 | >25 | 0.28 | >25 |
| V704_3000_240_RE_NT_pblib001_8 | 0.037 | 0.11 | 0.392 | 0.156 | 0.523 | >25 | >25 | >25 | 8.698 | >25 | >25 | 0.281 | >25 | 0.111 | >25 |
| V704_3000_240_RE_NT_pblib019_1 | 0.062 | 0.201 | 0.562 | 0.271 | 1.92 | >25 | >25 | >25 | 16.221 | >25 | >25 | 0.145 | >25 | 0.14 | >25 |
| V704_3008_040_RE_NT_pblib001_113 | 0.067 | 0.431 | 0.025 | 0.163 | 3.02 | >25 | 0.017 | 0.178 | >25 | >25 | >25 | >25 | >25 | 0.184 | >25 |
| V704_0372_250_RE_NT_pblib001_7 | 0.03 | 0.167 | 0.062 | 0.199 | 0.415 | 0.191 | >25 | >25 | >25 | >25 | >25 | >25 | >25 | 0.087 | >25 |
| V704_0372_250_RE_NT_pblib002_6 | 0.023 | 0.217 | ND | 0.151 | 0.326 | 2.426 | >25 | >25 | >25 | 2.114 | ND | ND | ND | 0.09 | ND |

### Table S1 C. AMP V703 IC80

| Sequence Name HVTN 703 | 1-18 | VRC07-523 LS | VRC01-23 LS | 3BNC117-LS | VRC01 | CH235.12 | PGT121-414 LS | 10-1074 LS | PGDM1400 | CAP256 VRC26.25 LS | PGT151 | VRC34.01 | ACS202 | 10E8v4 | CAP248-2B |
|---|---|---|---|---|---|---|---|---|---|---|---|---|---|---|---|
| V703_0013_090_RE_con_s | 1.93 | 1.1200 | 1.07 | >25 | 9.2118 | >10 | >25 | >25 | 0.0200 | 0.0300 | >10 | >10 | >10 | 16.9200 | >10 |
| V703_0203_081_RE_pblib002_s | 1.91 | 1.6100 | 5.69 | >25 | 15.2818 | >10 | 0.1100 | 0.1000 | >25 | >25 | >10 | >10 | >10 | 2.4500 | >10 |
| V703_0203_081_RE_sga8A5_s | 2.26 | 1.5400 | 3.27 | >25 | >25 | >10 | 0.0800 | 0.0600 | >25 | >25 | >10 | >10 | >10 | 0.6200 | >10 |
| V703_0217_050_RE_pblib002_s | 0.01 | 0.1700 | 0.24 | >25 | >25 | 0.79 | >25 | >25 | 0.0100 | 0.0100 | 0.0162 | 0.06 | >10 | 0.8100 | 0.04 |
| V703_0217_050_RE_sga2A3_s | 0.08 | 0.1100 | 0.22 | >25 | >25 | 1.16 | >25 | >25 | 0.0100 | 0.0020 | 0.04 | 0.09 | >10 | 1.0000 | >10 |
| V703_0279_110_RE_con_s-modified | 0.36 | 1.0900 | 0.51 | 1.3500 | 3.9365 | >10 | 5.6000 | 3.5300 | >25 | >25 | >10 | >10 | >10 | 0.6100 | >10 |
| V703_0279_110_RE_pblib002_s | 0.64 | 0.5800 | 0.35 | 1.5100 | 4.6208 | >10 | 3.4600 | 5.9200 | >25 | >25 | >10 | >10 | >10 | 1.3000 | >10 |
| V703_0472_030_RE_con_s | 0.0115 | 0.0100 | <0,005 | >25 | 0.0735 | 0.18 | >25 | >25 | 2.9900 | 0.0400 | 0.13 | >10 | >10 | 2.7200 | >10 |
| V703_0537_110_RE_sga4H1_s | 0.06 | 0.0400 | 0.01 | 0.0500 | >25 | 2.63 | 0.0500 | >25 | 0.4500 | 0.1900 | >10 | >10 | >10 | 4.3200 | >10 |
| V703_0537_110_RE_sga2B5_s | 0.02 | 0.0900 | 0.0072 | 0.2800 | 0.0695 | 0.13 | 0.1200 | 0.4400 | 0.8100 | 0.6000 | >10 | >10 | >10 | 3.8200 | >10 |
| V703_0537_110_RE_pblib003_s | 0.0411 | 0.0600 | 0.01 | 0.0800 | 0.1947 | 0.56 | 0.1500 | 0.5600 | 5.7800 | 11.0400 | >10 | >10 | >10 | 1.7700 | >10 |
| V703_0566_160_RE_con_s | 0.2 | 0.3100 | 0.65 | 0.9400 | 5.2121 | 1.33 | 0.0300 | 0.0300 | >25 | >25 | >10 | 0.3 | >10 | 5.3200 | >10 |
| V703_0566_160_RE_pblib002_s | 0.21 | 0.4900 | 0.64 | 4.5100 | 10.6147 | 3.6 | 0.0600 | 0.1300 | >25 | >25 | >10 | 0.27 | >10 | 21.2100 | >10 |
| V703_0629_150_RE_pblib002_s | 0.31 | 0.2300 | 0.28 | 0.5000 | 4.3929 | 2.3 | >25 | 10.4000 | 4.2400 | >25 | 0.04 | 0.87 | >10 | 3.6200 | >10 |
| V703_0629_150_RE_sga5A2_s | 0.23 | 0.6800 | 0.0775 | 1.2300 | 2.5153 | 0.73 | >25 | 20.1100 | 13.4300 | 5.2800 | 0.02 | >10 | >10 | 19.5800 | >10 |
| V703_0629_150_RE_pblib003_s | 0.29 | 1.1600 | 0.19 | 2.2300 | 3.7899 | 4.66 | >25 | 22.4500 | >25 | >25 | 0.19 | >10 | >10 | 20.8200 | >10 |
| V703_0629_150_RE_pblib004_s | 0.31 | 0.5800 | 0.24 | 4.4000 | 3.3096 | 1.51 | >25 | 11.6400 | 0.6400 | >25 | 0.04 | >10 | >10 | 7.9700 | >10 |
| V703_0646_051_RE_con_s | 1.85 | 1.2900 | 3 | 4.2900 | 15.7791 | >10 | >25 | >25 | 0.0500 | 0.0300 | >10 | 0.32 | >10 | 8.7100 | >10 |
| V703_0739_110_RE_con_s-modified | 0.1 | 0.1400 | 0.07 | 0.1900 | 0.4577 | 1.11 | 2.3000 | 0.5800 | 3.2300 | 0.0100 | >10 | >10 | >10 | 7.5400 | >10 |
| V703_0842_200_RE_con_s | >10 | 5.3600 | 6.75 | 8.9200 | >25 | >10 | >25 | 14.4900 | 0.3000 | 0.0700 | >10 | >10 | >10 | 5.1500 | >10 |
| V703_0865_070_RE_con_s | 0.39 | 4.8300 | ND | ND | ND | 4.15 | ND | ND | ND | ND | >10 | >10 | >10 | ND | ND |
| V703_0926_070_RE_sga2H2_s | 7.85 | 2.4000 | 2.36 | >25 | >25 | 9.8 | 0.1300 | 0.3200 | 4.0700 | >25 | >10 | >10 | >10 | 2.7100 | >10 |
| V703_1060_080_RE_con_s | 0.76 | 0.5600 | 1 | 3.5300 | 6.2655 | 4.39 | >25 | 19.0600 | >25 | 1.6700 | >10 | >10 | >10 | 0.3900 | >10 |
| V703_1104_100_RE_pblib001_s | 0.63 | 9.7000 | 3.69 | 14.4000 | >25 | >10 | >25 | >25 | 0.9500 | 3.5400 | >10 | >10 | >10 | 5.2000 | >10 |
| V703_1104_100_RE_sga10A5_s | 0.93 | 5.8200 | 4.4 | 8.7200 | >25 | >10 | >25 | 12.3000 | 3.0000 | 4.2800 | >10 | >10 | >10 | 1.5200 | >10 |
| V703_1104_100_RE_sga10G5_s | 0.85 | 7.2800 | 4.31 | 15.4100 | 22.0665 | >10 | >25 | >25 | 2.4300 | >25 | >10 | >10 | >10 | 2.8500 | >10 |
| V703_1298_080_RE_pblib002_s | 0.77 | 25.0000 | 0.38 | >25 | >25 | 0.59 | >25 | >25 | 0.8400 | 0.0700 | 9.49 | >10 | >10 | 4.3500 | >10 |
| V703_1313_040_RE_con_s | 0.26 | 0.1200 | 0.24 | 0.4200 | 2.4732 | >10 | 0.0500 | 0.2300 | 0.0157 | 0.0200 | >10 | >10 | >10 | 0.4300 | >10 |
| V703_1471_190_RE_con_s | 0.15 | 0.6100 | 0.07 | 0.2200 | 0.5218 | 1.89 | 0.0200 | 0.2600 | 0.1400 | 0.0200 | 0.02 | >10 | >10 | 25.0000 | >10 |
| V703_1675_080_RE_con_s-modified | 0.09 | 0.1600 | 0.19 | 0.1600 | 1.3422 | 0.55 | 5.5000 | 10.6700 | 0.0400 | 0.0100 | 4.81 | >10 | >10 | >25 | >10 |
| V703_1675_080_RE_con_s | 0.11 | 0.1500 | 0.17 | 0.1200 | 1.3686 | 0.7 | 1.1800 | 3.8500 | 0.0200 | 0.0100 | >10 | >10 | >10 | >25 | >10 |
| V703_1750_140_RE_con_s | 0.69 | 0.2100 | 0.52 | 0.7200 | 2.8549 | 3.32 | 0.2000 | 0.2700 | >25 | >25 | 0.3 | 8.3 | >10 | 2.9100 | >10 |
| V703_1764_250_RE_con_s | 0.09 | 0.2300 | 0.2 | 0.2600 | 0.8038 | 0.57 | 0.5700 | 0.2900 | >25 | >25 | >10 | >10 | >10 | 4.0300 | >10 |
| V703_1789_230_RE_sga3A3_s | 0.43 | 4.2100 | 1.75 | >25 | >25 | >10 | 2.8300 | 1.1300 | >25 | >25 | >10 | >10 | >10 | 14.8400 | >10 |
| V703_1848_190_RE_pblib002_s | **0.29** | **0.5100** | **0.24** | **0.1900** | **4.2223** | **3.7** | >25 | >25 | **0.1100** | **2.3400** | **>10** | **>10** | **>10** | **1.4100** | **>10** |
| V703_1848_190_RE_sga6D1_s | **0.36** | **0.9000** | **0.65** | **0.6900** | **4.1500** | **0.92** | >25 | >25 | >25 | >25 | **0.14** | **>10** | **>10** | **8.3000** | **>10** |
| V703_1889_100_RE_con_s | 0.33 | 20.0400 | 0.9 | >25 | >25 | 9.48 | 0.3500 | 0.6800 | 0.0200 | 0.0047 | >10 | >10 | >10 | 25.0000 | >10 |
| V703_1915_250_RE_sgaA3_s | 0.28 | 0.6300 | 0.18 | 2.7700 | >25 | >10 | 1.1000 | 2.3500 | >25 | 15.7800 | 5.62 | >10 | >10 | 12.4700 | >10 |
| V703_2018_240_RE_sga6A1_s | 0.68 | 0.8800 | 0.31 | 12.6600 | 2.8441 | >10 | 0.1100 | 0.1600 | 0.0031 | 0.0003 | >10 | >10 | >10 | 11.3200 | >10 |
| V703_2117_110_RE_sga2A10_s | 0.37 | 0.0800 | 0.11 | 0.3500 | 0.6427 | 1.33 | >25 | >25 | 0.0028 | 0.0016 | >10 | >10 | >10 | 3.6900 | >10 |
| V703_2149_060_RE_sgaB10_s | 0.1 | 0.1700 | 0.07 | >25 | 0.6489 | >10 | 0.2700 | 0.1400 | 0.0040 | 0.0050 | >10 | >10 | >10 | 4.0000 | >10 |
| V703_2304_150_RE_con_s | 0.24 | 0.0400 | 0.0413 | 0.8600 | 0.5357 | 0.22 | 0.2900 | 0.1700 | 0.1200 | 0.0100 | >10 | 0.11 | >10 | 3.8700 | >10 |
| V703_2539_070_RE_sga6F6_s | >10 | 1.0600 | >10 | >25 | >25 | >10 | 0.0400 | 0.1200 | >25 | >25 | >10 | >10 | >10 | 0.9400 | >10 |
| V703_2631_150_RE_sga2F8_s | 0.12 | 0.2600 | 0.26 | 0.1900 | 1.8281 | 2.8 | >25 | >25 | 0.0100 | 0.0021 | >10 | 3.9 | >10 | 6.7200 | >10 |
| V703_2788_030_RE_con_s | 2.55 | 0.5900 | 7.44 | 0.2300 | 2.7234 | 7.8 | 0.8400 | 0.1600 | >25 | 0.6900 | >10 | >10 | >10 | 19.7300 | >10 |
| V703_2805_080_RE_con_s | 0.1 | 0.0300 | 0.0985 | 0.0900 | 1.2914 | 0.85 | >25 | >25 | >25 | 0.0009 | 0.5 | 0.81 | >10 | 6.1900 | >10 |
| V703_2805_080_RE_pblib002_s | >10 | 11.0700 | >10 | >25 | >25 | >10 | >25 | >25 | >25 | >25 | >10 | 9.05 | >10 | 16.2600 | >10 |

### Table S1 D. AMP V704 IC80

| | IC80 | | | | | | | | | | | | | | |
|---|---|---|---|---|---|---|---|---|---|---|---|---|---|---|---|
| | CD4bs | | | | | | V3 | | V2 | | FP | | | MPER | |
| Sequence Name HVTN 704 | 1-18 | VRC07-523 LS | VRC01-23 LS | 3BNC117-LS | VRC01 | CH235.12 | PGT121-414 LS | 10-1074 LS | PGDM1400 | CAP256 VRC26.25 LS | PGT151 | VRC34.01 | ACS202 | 10E8v4 | CAP248-2B |
| V704_0026_231_RE_NT_pblib001_3 | 0.34 | 0.221 | 0.321 | 0.348 | 1.3 | >25 | 0.018 | 0.057 | 1.186 | >25 | >25 | >25 | >25 | 0.66 | >25 |
| V704_0128_220_RE_NT_pblib001_52 | 0.076 | 0.017 | 0.032 | 0.202 | 2.35 | 0.068 | 0.061 | 0.161 | 0.024 | 0.184 | >25 | 10.094 | >25 | 5.112 | ND |
| V704_0128_220_RE_NT_pblib002_40 | 0.134 | 0.07 | 0.12 | 0.092 | 0.37 | 0.329 | 0.131 | 0.464 | 0.108 | 0.253 | >25 | >25 | >25 | 0.38 | >25 |
| V704_0445_180_RE_NT_pblib001_22 | 0.035 | 0.694 | 0.09 | >25 | >25 | 0.948 | 1.497 | 0.255 | 1.096 | >25 | 0.557 | >25 | 0.79 | 4.358 | >25 |
| V704_0496_040_RE_NT_pblib001_37 | 0.142 | 0.565 | 2.091 | >25 | 12.57 | 4.89 | 0.457 | 0.238 | 0.021 | 2.61 | >25 | >25 | >25 | 1.207 | >25 |
| V704_0513_150_RE_NT_pblib007_2 | 1.477 | 0.706 | 0.732 | 0.673 | 2.47 | >25 | >25 | >25 | >25 | >25 | 1.89 | >25 | >25 | 4.3 | >25 |
| V704_0513_150_RE_NT_pblib002_10 | 1.315 | 0.887 | 1.007 | 0.687 | 3.64 | 19.266 | >25 | >25 | >25 | >25 | 1.084 | >25 | >25 | 4.13 | >25 |
| V704_0513_150_RE_NT_pblib001_34 | 1.71 | 1.037 | 0.892 | 0.785 | 3.2 | 22.674 | >25 | >25 | >25 | >25 | >25 | >25 | >25 | 4.04 | >25 |
| V704_0575_060_RE_NT_pblib002_16 | 1.278 | 3.279 | 2.171 | 4.895 | 6.12 | 6.285 | >25 | >25 | 0.188 | >25 | >25 | 0.849 | 0.193 | 2.06 | >25 |
| V704_0575_060_RE_NT_pblib001_60 | 1.926 | 4.692 | 2.289 | 6.593 | 6.718 | 8.447 | >25 | >25 | 0.23 | >25 | >25 | 2.715 | 0.287 | 1.53 | >25 |
| V704_0575_060_RE_NT_con02_s | 2.119 | 2.68 | 2.44 | 5.349 | 6.46 | 8.191 | >25 | >25 | 0.164 | >25 | >25 | >25 | 0.557 | 1.74 | >25 |
| V704_0644_060_RE_NT_pblib001_216 | 0.358 | 0.513 | 0.816 | 1.025 | 2.49 | 11.908 | >25 | 1.551 | >25 | >25 | >25 | >25 | >25 | 1.427 | >25 |
| V704_0726_080_RE_NT_pblib001_26 | 0.571 | 3.3 | 1.177 | 1.568 | 18.649 | 14.337 | 1.31 | 1.913 | 1.668 | >25 | >25 | >25 | >25 | 11.4 | >25 |
| V704_0746_760_RE_NT_pblib001_37 | 0.629 | 1.056 | 0.502 | 0.716 | 2.27 | >25 | 0.034 | 0.032 | >25 | >25 | >25 | >25 | >25 | 8.99 | >25 |
| V704_0746_760_RE_NT_pblib003_34 | 0.625 | 1.095 | 0.442 | 0.909 | 2.87 | >25 | 0.049 | 0.032 | >25 | >25 | >25 | >25 | >25 | 7.78 | >25 |
| V704_0746_760_RE_NT_pblib002_36 | 0.775 | 1.184 | 0.868 | 1.025 | 3.08 | >25 | 0.053 | 0.035 | >25 | >25 | >25 | >25 | >25 | 5.37 | >25 |
| V704_0847_030_RE_NT_pblib001_34 | >25 | 0.663 | 0.269 | >25 | 6.537 | >25 | 0.028 | 0.008 | >25 | >25 | >25 | >25 | >25 | 1.129 | >25 |
| V704_0847_030_RE_NT_pblib002_23 | >25 | 1.022 | 0.397 | >25 | 8.502 | >25 | 0.026 | 0.02 | >25 | >25 | >25 | >25 | >25 | 1.602 | >25 |
| V704_0855_080_RE_NT_pblib001_187 | 0.028 | 0.01 | 0.01 | 0.025 | 0.044 | >25 | 0.259 | 0.235 | >25 | >25 | 0.028 | >25 | >25 | 0.836 | >25 |
| V704_0856_240_RE_NT_pblib001_9 | 0.379 | 0.184 | 0.164 | 0.24 | 1.32 | 1.98 | 0.037 | 0.042 | >25 | >25 | >25 | >25 | >25 | 2.44 | >25 |
| V704_0886_250_RE_NT_pblib001_10 | 0.742 | 0.745 | 0.354 | 2.225 | 4.601 | >25 | 3.693 | 0.789 | >25 | >25 | >25 | >25 | >25 | 1.04 | >25 |
| V704_0907_130_RE_NT_pblib001_32 | 0.049 | 0.522 | 0.053 | 2.65 | 0.816 | >25 | 0.009 | 0.011 | >25 | >25 | 0.05 | 0.99 | 3.53 | 2.397 | >25 |
| V704_0911_150_RE_NT_pblib001_58 | 1.545 | 1.77 | 3.313 | 1.814 | 3.503 | >25 | 0.032 | 0.04 | >25 | >25 | >25 | >25 | >25 | 4.326 | >25 |
| V704_0944_180_RE_NT_pblib001_76 | 0.077 | 0.068 | 0.021 | 0.221 | 0.24 | >25 | >25 | >25 | >25 | >25 | >25 | >25 | >25 | 20 | >25 |
| V704_1109_140_RE_NT_pblib001_191 | >25 | 0.774 | 6.788 | >25 | 8.7 | >25 | >25 | >25 | 0.449 | >25 | >25 | >25 | >25 | 4.7 | >25 |
| V704_1180_070_RE_NT_pblib001_87 | 0.017 | 0.155 | 0.031 | 0.062 | 0.47 | 0.487 | >25 | >25 | 1.031 | >25 | 0.027 | 2.325 | 6.402 | 2.06 | >25 |
| V704_1180_070_RE_NT_pblib027_1 | 0.021 | 0.193 | 0.026 | 0.062 | 0.419 | 1.701 | >25 | >25 | 2.394 | >25 | 0.027 | 3.647 | 6.066 | 0.88 | >25 |
| V704_1183_220_RE_NT_pblib001_70 | 0.115 | 0.73 | 0.086 | 0.23 | 0.91 | 3.05 | 0.09 | 0.134 | >25 | >25 | >25 | >25 | 5.842 | 1.77 | >25 |
| V704_1429_090_RE_NT_pblib002_75 | 0.421 | 0.25 | 0.212 | 6.731 | 0.618 | >25 | >25 | >25 | 0.339 | >25 | >25 | >25 | >25 | 2.683 | >25 |
| V704_1429_090_RE_NT_pblib001_111 | 0.322 | 0.287 | 0.318 | 9.705 | 0.727 | >25 | >25 | >25 | 0.892 | >25 | >25 | >25 | >25 | 1.737 | >25 |
| V704_1528_240_RE_NT_pblib001_17 | 0.238 | 1.509 | 0.114 | 0.464 | 7.13 | 24.662 | 0.012 | 0.012 | >25 | >25 | >25 | 0.385 | 0.181 | 0.949 | >25 |
| V704_1535_030_RE_NT_pblib001_471 | 0.129 | 0.183 | 0.054 | 0.061 | 0.281 | >25 | 0.041 | 0.241 | 4.061 | >25 | >25 | 1.36 | 5.17 | 0.143 | >25 |
| V704_1706_040_RE_NT_pblib001_26 | 1.103 | 0.203 | 0.161 | 0.924 | 1.25 | 0.641 | 1.784 | 1.126 | 13.413 | >25 | >25 | >25 | >25 | 1.95 | >25 |
| V704_1775_030_RE_NT_pblib001_135 | 0.014 | 0.077 | 0.073 | 6.421 | 0.487 | 0.166 | >25 | >25 | 0.007 | >25 | 1.767 | >25 | >25 | 1.811 | >25 |
| V704_1783_150_RE_NT_pblib001_144 | 0.02 | 0.15 | 0.027 | 0.09 | 0.28 | >25 | 1.474 | 1.766 | >25 | >25 | >25 | >25 | >25 | 1.49 | >25 |
| V704_1835_150_RE_NT_pblib001_10 | 0.079 | >25 | 0.187 | >25 | >25 | >25 | >25 | 0.312 | >25 | >25 | 1.176 | >25 | 1.291 | 0.982 | >25 |
| V704_1835_150_RE_NT_pblib002_9 | 0.107 | >25 | 0.179 | >25 | >25 | >25 | >25 | 0.346 | >25 | >25 | 5.51 | >25 | 6.834 | 0.88 | >25 |
| V704_1930_170_RE_NT_pblib002_114 | 0.115 | 0.086 | 0.04 | 0.1 | 0.297 | 0.394 | >25 | >25 | 0.063 | >25 | >25 | >25 | >25 | 6.298 | >25 |
| V704_1930_170_RE_NT_pblib001_234 | 0.148 | 0.121 | 0.079 | 0.121 | 0.376 | 0.424 | >25 | >25 | 0.063 | >25 | >25 | >25 | >25 | 20 | >25 |
| V704_1991_230_RE_NT_pblib001_38 | 0.329 | 0.951 | 0.606 | 0.839 | 4.22 | 1.374 | >25 | >25 | 0.109 | 0.0001 | >25 | >25 | 1.054 | 2.54 | >25 |
| V704_1991_230_RE_NT_pblib002_19 | 0.23 | 0.753 | 0.523 | 0.581 | 4.21 | 1.059 | >25 | >25 | 0.173 | 0.057 | >25 | >25 | 0.589 | 1.433 | >25 |
| V704_2065_060_RE_NT_pblib001_47 | 0.405 | 0.691 | 0.076 | 0.316 | 0.603 | 0.572 | >25 | >25 | >25 | >25 | >25 | >25 | >25 | 0.729 | >25 |
| V704_2065_060_RE_NT_pblib002_42 | 0.335 | 0.854 | 0.121 | 0.292 | 0.67 | 0.378 | >25 | >25 | >25 | >25 | >25 | >25 | >25 | 1.33 | >25 |
| V704_2095_130_RE_NT_pblib001_136 | 0.234 | 0.256 | 0.084 | 0.124 | 0.82 | 1.004 | >25 | 0.239 | >25 | >25 | >25 | >25 | >25 | 2.82 | >25 |
| V704_2448_240_RE_NT_pblib001_288 | 0.489 | 0.297 | 0.03 | 0.211 | 0.412 | 0.393 | >25 | >25 | >25 | >25 | >25 | >25 | >25 | 20 | >25 |
| V704_2541_080_RE_NT_pblib001_13 | 0.109 | 0.152 | 0.088 | 0.093 | 0.74 | 0.763 | 0.136 | 0.026 | 9.447 | >25 | 0.161 | 2.733 | >25 | 1.12 | >25 |
| V704_2541_080_RE_NT_pblib042_1 | 0.058 | 0.118 | 0.05 | 0.071 | 0.58 | 0.391 | 0.074 | 0.088 | 1.459 | >25 | >25 | >25 | >25 | 0.73 | >25 |
| V704_2541_080_RE_NT_pblib003_5 | 0.152 | 0.246 | 0.071 | 0.148 | 1.02 | 1.417 | 0.264 | 0.042 | 18.004 | >25 | 0.215 | 1.791 | >25 | 0.82 | >25 |
| V704_2544_140_RE_NT_pblib002_10 | 0.493 | 0.424 | 0.346 | 0.298 | 2.24 | >25 | 2.283 | 2.125 | >25 | >25 | >25 | >25 | >25 | 1.567 | >25 |
| V704_2544_140_RE_NT_pblib001_11 | 3.149 | 1.12 | 1.339 | 2.977 | 14.88 | >25 | >25 | 1.314 | >25 | >25 | 0.232 | >25 | >25 | 4.195 | >25 |
| V704_2544_150_RE_NT_pblib001_33 | 0.871 | 0.549 | 0.687 | 1.181 | 5.31 | 16.051 | 0.106 | 0.171 | 0.28 | >25 | >25 | >25 | >25 | 1.12 | >25 |
| V704_2555_240_RE_NT_pblib001_231 | 0.38 | 0.148 | 0.087 | 0.258 | 1.061 | >25 | 0.052 | 0.057 | >25 | >25 | >25 | >25 | >25 | 20 | >25 |
| V704_2684_181_RE_NT_pblib001_13 | 0.029 | 0.117 | 0.044 | 0.139 | 0.31 | >25 | 0.018 | 0.05 | >25 | >25 | >25 | >25 | >25 | 0.12 | >25 |
| V704_2684_181_RE_NT_pblib006_3 | 0.033 | 0.158 | 0.043 | 0.131 | 0.275 | >25 | 0.019 | 0.057 | >25 | >25 | >25 | >25 | >25 | 0.13 | >25 |
| V704_2767_070_RE_NT_pblib001_63 | 0.476 | 0.252 | 0.184 | 2.979 | 8.32 | 3.189 | 0.195 | 0.63 | >25 | >25 | 0.854 | >25 | >25 | 7.16 | >25 |
| V704_2788_060_RE_NT_pblib001_96 | 0.221 | 0.45 | 0.714 | 0.609 | 3.171 | 2.516 | 0.571 | 0.536 | 0.008 | 0.0001 | >25 | >25 | >25 | 2.06 | >25 |
| V704_2788_060_RE_NT_pblib002_20 | 0.502 | 0.596 | 0.735 | 0.708 | 3.5 | 3.104 | 0.356 | 0.381 | 0.01 | 0.01 | >25 | >25 | >25 | 0.434 | >25 |
| V704_2834_210_RE_NT_pblib002_26 | 0.229 | 0.727 | 0.3 | 0.23 | 5.79 | >25 | >25 | 18.87 | >25 | >25 | >25 | 5.353 | >25 | 1.84 | >25 |
| V704_2834_210_RE_NT_pblib001_44 | 0.328 | 1.11 | 0.211 | 0.253 | 5.62 | >25 | >25 | 15.601 | >25 | >25 | >25 | 3.189 | >25 | 0.415 | >25 |
| V704_2839_140_RE_NT_pblib001_208 | 0.318 | 0.5 | 0.731 | 5.705 | 3.96 | >25 | >25 | >25 | >25 | >25 | >25 | 8.989 | >25 | 2.355 | >25 |
| V704_2981_150_RE_NT_pblib004_2 | 1.826 | 1.134 | 0.288 | 0.351 | 6.36 | 0.929 | 8.173 | 2.389 | 0.704 | >25 | >25 | >25 | >25 | 0.922 | >25 |
| V704_2981_150_RE_NT_pblib001_33 | >25 | 3.909 | 3.159 | 3.012 | 16.5 | 7.835 | 3.613 | 1.139 | 0.06 | >25 | >25 | >25 | >25 | 9.84 | >25 |
| V704_2981_150_RE_NT_pblib002_11 | 4.473 | 1.502 | 0.649 | 0.917 | 4.94 | 2.453 | >25 | >25 | >25 | >25 | >25 | >25 | >25 | 4.03 | >25 |
| V704_2981_150_RE_NT_pblib003_4 | 4.22 | 1.355 | 0.497 | 0.53 | 5.58 | 0.886 | >25 | >25 | >25 | >25 | >25 | >25 | >25 | 1.17 | >25 |
| V704_3000_240_RE_NT_pblib001_8 | 0.175 | 0.287 | 1.724 | 0.527 | 1.79 | >25 | >25 | >25 | >25 | >25 | >25 | 3.358 | >25 | ND | >25 |
| V704_3000_240_RE_NT_pblib019_1 | 0.178 | 0.461 | 1.597 | 1.445 | 5.22 | >25 | >25 | >25 | >25 | >25 | >25 | 8.711 | >25 | 0.745 | >25 |
| V704_3008_040_RE_NT_pblib001_113 | 0.402 | 1.972 | 0.092 | 0.704 | 9.935 | >25 | 0.22 | 4.87 | >25 | >25 | >25 | >25 | >25 | 1.11 | >25 |
| V704_0372_250_RE_NT_pblib001_7 | 0.165 | 1.52 | 0.284 | 1.47 | 1.32 | 3.16 | >25 | >25 | >25 | >25 | >25 | >25 | >25 | 0.427 | >25 |
| V704_0372_250_RE_NT_pblib002_6 | 0.125 | 0.529 | ND | 0.986 | 0.868 | >25 | >25 | >25 | >25 | >25 | ND | ND | ND | ND | ND |

**Table S1. IC50 and IC80 scores for the AMP V703 and V704 study sequences. A.** IC50 AMP V703. **B.** IC50 AMP V704. **C.** IC80 AMP V703. **D.** IC80 AMP V704.When more than one transmitted founder lineage was found in an individual, the set of TFLs are grouped in these tables. For designing SHEP-T2 and class specific sensitive panels, we used the most sensitive variant overall to represent an individual, and those are marked in grey on the left. For the REP-

T2 the least sensitive from an individual was used, are these are marked in yellow. "ND" means that the assay was not done.

### Table S2 A. AMP V704 nomenclature key

| Accession number | Isolate name in Genbank, sequence name | NIH repository name | Plasmid_name | Subtype | Country | Sampling Year | Footnotes |
|---|---|---|---|---|---|---|---|
| OQ912888 | V704_0026_231_RE_NT_pblib001_3 | AMP1-P136 | H704_0026_231_RE_pbsga001_s | B | PERU | 2019 | |
| ON980817 | V704_0128_220_RE_NT_pblib001_52 | AMP1-P101.1 | H704_0128_220_RE_pb001_s | BF1 | PERU | 2019 | |
| ON980818 | V704_0128_220_RE_NT_pblib002_40 | AMP1-P101.2 | V704_0128_220_RE_pblib002_s | BF1 | PERU | 2019 | |
| OQ912889 | V704_0372_250_RE_NT_pblib001_7 | AMP1-P137.1 | V704_0372_250_RE_pblib001_s | B | PERU | 2019 | |
| OQ912890 | V704_0372_250_RE_NT_pblib002_6 | AMP1-P137.2 | V704_0372_250_RE_pblib002_s | B | PERU | 2019 | |
| ON980823 | V704_0445_180_RE_NT_pblib001_22 | AMP1-P102 | H704_0445_180_RE_con_s | B | PERU | 2019 | |
| ON980826 | V704_0496_040_RE_NT_pblib001_37 | AMP1-P103 | H704_0496_040sN | F1 | BRAZIL | 2017 | |
| ON980827 | V704_0513_150_RE_NT_pblib001_34 | AMP1-P104.1 | H704_0513_150_eN01T | B | PERU | 2018 | |
| ON980828 | V704_0513_150_RE_NT_pblib002_10 | AMP1-P104.2 | H704_0513_150_eN02C | B | PERU | 2018 | |
| ON980829 | V704_0513_150_RE_NT_pblib007_2 | AMP1-P104.3 | H704_0513_150_RE_pblib003_s | B | PERU | 2018 | |
| ON980835 | V704_0575_060_RE_NT_con02_s | AMP1-P105.2 | V704_0575_060_RE_NT_con02_s | B | PERU | 2018 | 2 |
| ON980834 | V704_0575_060_RE_NT_pblib001_60 | AMP1-P105.1 | H704_0575_060_EsN1A | B | PERU | 2018 | |
| ON980836 | V704_0575_060_RE_NT_pblib002_16 | AMP1-P105.3 | H704_0575_060_RE_p002s | B | PERU | 2018 | |
| ON980839 | V704_0644_060_RE_NT_pblib001_216 | AMP1-P106 | H704_0644_060sN_prelimSeq | B | PERU | 2018 | |
| ON980844 | V704_0726_080_RE_NT_pblib001_26 | AMP1-P107 | H704_0726_080sN | B | UNITED STATES | 2017 | |
| ON980846 | V704_0746_760_RE_NT_pblib001_37 | AMP1-P108.2 | H704_0746_760_RE_p002s | B | PERU | 2018 | |
| ON980847 | V704_0746_760_RE_NT_pblib002_36 | AMP1-P108.3 | H704_0746_760_RE_p003s | B | PERU | 2018 | |
| ON980845 | V704_0746_760_RE_NT_pblib003_34 | AMP1-P108.1 | H704_0746_760_RE_p001s | B | PERU | 2018 | |
| ON980852 | V704_0847_030_RE_NT_pblib001_34 | AMP1-P109.1 | H704_0847_030_EsN_01T | B | PERU | 2018 | |
| ON980853 | V704_0847_030_RE_NT_pblib002_23 | AMP1-P109.2 | H704_0847_030_EsN_02C | B | PERU | 2018 | |
| ON980854 | V704_0855_080_RE_NT_pblib001_187 | AMP1-P110 | H704_0855_080_EsN | B | PERU | 2018 | |
| OQ912891 | V704_0856_240_RE_NT_pblib001_9 | AMP1-P138 | H704_0856_240_RE_pb001_s | B | UNITED STATES | 2019 | |
| OQ912892 | V704_0886_250_RE_NT_pblib001_10 | AMP1-P139 | H704_0886_250_RE_p001s | B | PERU | 2019 | |
| ON980862 | V704_0907_130_RE_NT_pblib001_32 | AMP1-P111 | H704_0907_130sN | B | UNITED STATES | 2017 | |
| ON980863 | V704_0911_150_RE_NT_pblib001_58 | AMP1-P112 | H704_0911_150sN | B | UNITED STATES | 2017 | |
| ON980866 | V704_0944_180_RE_NT_pblib001_76 | AMP1-P113 | H704_0944_180_RE_cs | B | PERU | 2019 | |
| ON980868 | V704_1109_140_RE_NT_pblib001_191 | AMP1-P114 | H704_1109_140_RE_cs | BF1 | PERU | 2019 | |
| ON980872 | V704_1180_070_RE_NT_pblib001_87 | AMP1-P115.2 | H704_1180_070EsN | A1B | PERU | 2018 | |
| ON980871 | V704_1180_070_RE_NT_pblib027_1 | AMP1-P115.1 | H704_1180_070_RE_pblib027_s | A1B | PERU | 2018 | |
| ON980873 | V704_1183_220_RE_NT_pblib001_70 | AMP1-P116 | H704_1183_220EsN | B | UNITED STATES | 2018 | |
| ON980885 | V704_1429_090_RE_NT_pblib001_111 | AMP1-P117.2 | H704_1429_090_eN2G | F2 | SWITZERLAND | 2017 | |
| ON980884 | V704_1429_090_RE_NT_pblib002_75 | AMP1-P117.1 | H704_1429_090_eN1A | F2 | SWITZERLAND | 2017 | |
| OQ912893 | V704_1528_240_RE_NT_pblib001_17 | AMP1-P140 | H704_1528_240_RE_pblib_001_s | B | PERU | 2019 | |
| ON980889 | V704_1535_030_RE_NT_pblib001_471 | AMP1-P118 | H704_1535_030sN | B | UNITED STATES | 2017 | |
| ON980897 | V704_1706_040_RE_NT_pblib001_26 | AMP1-P119 | H704_1706_050sN | B | PERU | 2018 | |
| ON980906 | V704_1775_030_RE_NT_pblib001_135 | AMP1-P120 | H704_1775_030cN_SynGtoA_567 | F1 | PERU | 2018 | |
| ON980907 | V704_1783_150_RE_NT_pblib001_144 | AMP1-P121 | H704_1783_150_RE_cs | B | PERU | 2019 | |
| ON980910 | V704_1835_150_RE_NT_pblib001_10 | AMP1-P122.1 | H704_1835_150_RE_p001s_2484A | B | PERU | 2018 | |
| ON980911 | V704_1835_150_RE_NT_pblib002_9 | AMP1-P122.2 | H704_1835_150_RE_p002s_2484A | B | PERU | 2018 | |
| ON980913 | V704_1930_170_RE_NT_pblib001_234 | AMP1-P123.2 | H704_1930_170_RE_c02s_1523G | BF1 | PERU | 2019 | |
| ON980912 | V704_1930_170_RE_NT_pblib002_114 | AMP1-P123.1 | H704_1930_170_RE_c01s_1523A | BF1 | PERU | 2019 | |
| ON980917 | V704_1991_230_RE_NT_pblib001_38 | AMP1-P124.1 | H704_1991_230_RE_p001s_1194T | B | PERU | 2018 | |
| ON980918 | V704_1991_230_RE_NT_pblib002_19 | AMP1-P124.2 | H704_1991_230_RE_p002s_1194T | B | PERU | 2018 | |
| ON980920 | V704_2065_060_RE_NT_pblib001_47 | AMP1-P125.1 | H704_2065_060_RE_p001s | B | PERU | 2018 | |
| ON980921 | V704_2065_060_RE_NT_pblib002_42 | AMP1-P125.2 | H704_2065_060_RE_p002s | B | PERU | 2018 | |
| ON980922 | V704_2095_130_RE_NT_pblib001_136 | AMP1-P126 | H704_2095_130_RE_cs | B | PERU | 2019 | |
| OQ912894 | V704_2448_240_RE_NT_pblib001_288 | AMP1-P141 | H704_2448_240_RE_cs | B | PERU | 2018 | |
| ON980937 | V704_2541_080_RE_NT_pblib003_5 | AMP1-P127.3 | H704_2541_080EsN | B | UNITED STATES | 2018 | 3 |
| ON980939 | V704_2541_080_RE_NT_pblib042_1 | AMP1-P127.2 | H704_2541_080_RE_p003s | B | UNITED STATES | 2018 | 3 |
| ON980938 | V704_2541_080_RE_NT_pblib001_13 | AMP1-P127.1 | H704_2541_080_RE_p001s | B | UNITED STATES | 2018 | 3 |
| ON980941 | V704_2544_140_RE_NT_pblib001_11 | AMP1-P128.2 | H704_2544_140eN02 | B | UNITED STATES | 2018 | 4 |
| ON980940 | V704_2544_140_RE_NT_pblib002_10 | AMP1-P128.1 | H704_2544_140eN01 | B | UNITED STATES | 2018 | 4 |
| ON980942 | V704_2544_150_RE_NT_pblib001_33 | AMP1-P128.3 | H704_2544_150_RE_p001s | B | UNITED STATES | 2018 | 4 |
| OQ912895 | V704_2555_240_RE_NT_pblib001_231 | AMP1-P142 | H704_2555_240_RE_con_s_vpuG | B | PERU | 2019 | 5 |
| ON980946 | V704_2684_181_RE_NT_pblib001_13 | AMP1-P129.1 | H704_2684_181_RE_p001s | B | PERU | 2018 | |
| ON980947 | V704_2684_181_RE_NT_pblib006_3 | AMP1-P129.2 | H704_2684_181_RE_pblib006_s | B | PERU | 2018 | |
| ON980949 | V704_2767_070_RE_NT_pblib001_63 | AMP1-P130 | H704_2767_070sN | B | PERU | 2017 | |
| ON980952 | V704_2788_060_RE_NT_pblib001_96 | AMP1-P131.1 | H704_2788_060eN_04 | BF1 | PERU | 2017 | |
| ON980953 | V704_2788_060_RE_NT_pblib002_20 | AMP1-P131.2 | H704_2788_060eN_12 | BF1 | PERU | 2017 | |
| ON980954 | V704_2834_210_RE_NT_pblib001_44 | AMP1-P132.1 | H704_2834_210_RE_pb001_s | B | BRAZIL | 2019 | |
| ON980955 | V704_2834_210_RE_NT_pblib002_26 | AMP1-P132.2 | H704_2834_210_RE_pb002_s | B | BRAZIL | 2019 | |
| ON980956 | V704_2839_140_RE_NT_pblib001_208 | AMP1-P133 | H704_2839_140_RE_cs | B | PERU | 2019 | |
| ON980961 | V704_2981_150_RE_NT_pblib001_33 | AMP1-P134.1 | H704_2981_150_RE_p001s_2559A | B | UNITED STATES | 2018 | |
| ON980962 | V704_2981_150_RE_NT_pblib002_11 | AMP1-P134.2 | H704_2981_150_RE_p002s_2559A | B | UNITED STATES | 2018 | |
| ON980963 | V704_2981_150_RE_NT_pblib003_4 | AMP1-P134.3 | H704_2981_150_RE_pblib003_s | B | UNITED STATES | 2018 | |
| ON980964 | V704_2981_150_RE_NT_pblib004_2 | AMP1-P134.4 | H704_2981_150_RE_pblib005_s | B | UNITED STATES | 2018 | |
| OQ912896 | V704_3000_240_RE_NT_pblib001_8 | AMP1-P143.1 | H704_3000_240_RE_pbsga001_s | B | PERU | 2018 | |
| OQ912897 | V704_3000_240_RE_NT_pblib019_1 | AMP1-P143.2 | H704_3000_240_RE_pbsga002_s | B | PERU | 2018 | |
| ON980965 | V704_3008_040_RE_NT_pblib001_113 | AMP1-P135 | H704_3008_040EsN | B | PERU | 2018 | |

### Table S2 B. AMP V703 nomenclature key

| Accession number | Isolate name in Genbank, sequence name | NIH repository name | Plasmid_name | Subtype | Country | Sampling Year | Footnotes |
|---|---|---|---|---|---|---|---|
| ON890939 | V703_0013_090_RE_con_s | AMP1-P001 | H703_0013_090Es | C | SOUTH AFRICA | 2019 | |
| ON890946 | V703_0203_081_RE_pblib002_s | AMP1-P002.1 | V703_0203_081_RE_pblib002_s | C | MALAWI | 2019 | |
| ON890947 | V703_0203_081_RE_sga8A5_s | AMP1-P002.2 | H703_0203_081_RE_e8A5s | C | MALAWI | 2019 | |
| ON890948 | V703_0217_050_RE_pblib002_s | AMP1-P003.1 | V703_0217_050_RE_pblib002_s | C | ZAMBIA | 2017 | |
| ON890949 | V703_0217_050_RE_sga2A3_s | AMP1-P003.2 | H703_0217_050e_2A3 | C | ZAMBIA | 2017 | |
| ON890950 | V703_0279_110_RE_con_s-modified | AMP1-P004.1 | H703_0279_110s | C | SOUTH AFRICA | 2018 | |
| ON890951 | V703_0279_110_RE_pblib002_s | AMP1-P004.2 | V703_0279_110_RE_pblib002_s | C | SOUTH AFRICA | 2018 | |
| ON890962 | V703_0472_030_RE_con_s | AMP1-P005 | H703_0472_030s | C | MALAWI | 2017 | |
| ON890973 | V703_0537_110_RE_pblib003_s | AMP1-P006.1 | V703_0537_110_RE_pblib003_s | C | SOUTH AFRICA | 2017 | |
| ON890974 | V703_0537_110_RE_sga2B5_s | AMP1-P006.2 | H703_0537_110s_2B5 | C | SOUTH AFRICA | 2017 | |
| ON890975 | V703_0537_110_RE_sga4H1_s | AMP1-P006.3 | H703_0537_110s_4H1 | C | SOUTH AFRICA | 2017 | |
| ON890976 | V703_0566_160_RE_con_s | AMP1-P007.1 | H703_0566_160s | C | SOUTH AFRICA | 2017 | |
| ON890977 | V703_0566_160_RE_pblib002_s | AMP1-P007.2 | V703_0566_160_RE_pblib002_s | C | SOUTH AFRICA | 2017 | |
| ON890982 | V703_0629_150_RE_pblib002_s | AMP1-P008.1 | V703_0629_150_RE_pblib002_s | C | BOTSWANA | 2018 | |
| ON890983 | V703_0629_150_RE_pblib003_s | AMP1-P008.2 | V703_0629_150_RE_pblib003_s | C | BOTSWANA | 2018 | |
| ON890984 | V703_0629_150_RE_pblib004_s | AMP1-P008.3 | V703_0629_150_RE_pblib004_s | C | BOTSWANA | 2018 | |
| ON890985 | V703_0629_150_RE_sga5A2_s | AMP1-P008.4 | H703_0629_150_RE_e5A2s | C | BOTSWANA | 2018 | |
| ON890987 | V703_0646_051_RE_con_s | AMP1-P009 | H703_0646_051sN | C | SOUTH AFRICA | 2017 | |
| ON890990 | V703_0739_110_RE_con_s-modified | AMP1-P010 | H703_0739_110s | C | SOUTH AFRICA | 2017 | |
| ON890997 | V703_0842_200_RE_con_s | AMP1-P011 | H703_0842_200Es | C | ZAMBIA | 2018 | |
| ON890999 | V703_0865_070_RE_con_s | AMP1-P012 | H703_0865_070s | G | KENYA | 2017 | |
| ON891001 | V703_0926_070_RE_sga2H2_s | AMP1-P013 | H703_0926_070s_2H2 | C | SOUTH AFRICA | 2017 | |
| ON891010 | V703_1060_080_RE_con_s | AMP1-P014 | H703_1060_080s | C | SOUTH AFRICA | 2017 | |
| ON891011 | V703_1104_100_RE_pblib001_s | AMP1-P015.1 | V703_1104_100_RE_pblib001_s | C | ZAMBIA | 2017 | |
| ON891012 | V703_1104_100_RE_sga10A5_s | AMP1-P015.2 | H703_1104_100_RE_e10A5s | C | ZAMBIA | 2017 | |
| ON891013 | V703_1104_100_RE_sga10G5_s | AMP1-P015.3 | H703_1104_100_RE_e10G5s | C | ZAMBIA | 2017 | |
| ON891026 | V703_1313_040_RE_con_s | AMP1-P016 | H703_1313_040s | C | SOUTH AFRICA | 2017 | |
| ON891037 | V703_1471_190_RE_con_s | AMP1-P017 | H703_1471_190s | C | SOUTH AFRICA | 2017 | |
| ON891047 | V703_1675_080_RE_con_s | AMP1-P018.2 | H703_1675_080s | C | SOUTH AFRICA | 2017 | |
| ON891046 | V703_1675_080_RE_con_s-modified | AMP1-P018.1 | H703_1675_080s_G613S | C | SOUTH AFRICA | 2017 | |
| ON891053 | V703_1750_140_RE_con_s | AMP1-P019 | H703_1750_140Es | C | SOUTH AFRICA | 2018 | |
| ON891055 | V703_1764_250_RE_con_s | AMP1-P020.1 | H703_1764_250_RE_cs | C | SOUTH AFRICA | 2019 | |
| ON891056 | V703_1764_250_RE_pblib002_s | AMP1-P020.2 | V703_1764_250_RE_pblib002_s | C | SOUTH AFRICA | 2019 | |
| ON891058 | V703_1789_230_RE_sga3A3_s | AMP1-P021 | H703_1789_230_RE_e3A3s | C | MALAWI | 2018 | |
| ON891062 | V703_1848_190_RE_pblib002_s | AMP1-P022.1 | V703_1848_190_RE_pblib002_s | C | MALAWI | 2019 | |
| ON891063 | V703_1848_190_RE_sga6D1_s | AMP1-P022.2 | H703_1848_190_RE_e6D1s | C | MALAWI | 2019 | |
| ON891068 | V703_1889_100_RE_con_s | AMP1-P023 | H703_1889_100Es | C | SOUTH AFRICA | 2018 | |
| ON891069 | V703_1915_250_RE_sgaA3_s | AMP1-P024.1 | V703_1915_250_RE_sgaA3_s | C | SOUTH AFRICA | 2019 | |
| ON891070 | V703_1915_250_RE_sgaC2a_s | AMP1-P024.2 | V703_1915_250_RE_sgaC2a_s | C | SOUTH AFRICA | 2019 | |
| ON891075 | V703_2018_240_RE_sga6A1_s | AMP1-P025 | H703_2018_240_RE_e6A1s | C | SOUTH AFRICA | 2019 | |
| ON891077 | V703_2117_110_RE_sga2A10_s | AMP1-P026 | H703_2117_110_RE_e2A10s | C | SOUTH AFRICA | 2019 | |
| ON891079 | V703_2149_060_RE_sgaB10_s | AMP1-P027 | H703_2149_060_RE_eB10s | C | MALAWI | 2018 | |
| ON891080 | V703_2304_150_RE_con_s | AMP1-P028 | H703_2304_150_RE_cs | C | SOUTH AFRICA | 2019 | |
| ON891082 | V703_2539_070_RE_pblib002_s | AMP1-P029.1 | V703_2539_070_RE_pblib002_s | C | SOUTH AFRICA | 2018 | 1 |
| ON891083 | V703_2539_070_RE_sga6F6_s | AMP1-P029.2 | H703_2539_070_RE_e6F6s | C | SOUTH AFRICA | 2018 | 1 |
| ON891084 | V703_2631_150_RE_sga2F8_s | AMP1-P030 | H703_2631_150_RE_e2F8s | C | MALAWI | 2019 | |
| ON891087 | V703_2788_030_RE_con_s | AMP1-P031 | H703_2788_030Es_B1 | C | MALAWI | 2018 | |
| ON891088 | V703_2805_080_RE_con_s | AMP1-P032.1 | H703_2805_080Es | C | MOZAMBIQUE | 2018 | |
| ON891089 | V703_2805_080_RE_pblib002_s | AMP1-P032.2 | V703_2805_080_RE_pblib002_s | C | MOZAMBIQUE | 2018 | |

**Footnotes:**

1. V703_2539 also called V703_1039
2. Synthesized gene, no exact match in vivo
3. V704_2541 also called V704_1041
4. V704_2544 also called V704_1044
5. V704_2555 also called V704_1055

**Table S2. Nomenclature key and critical sequence metadata. A.** V704 study sequences. **B.** V703 study sequences.

### Table S3

| | | | | IC50 | | | | | | | | | | | | |
|---|---|---|---|---|---|---|---|---|---|---|---|---|---|---|---|---|
| | | | | CD4bs | | | | | V3 | | V2 | | FP | | | MPER |
| | Sequence Name | IC50 Rank Mean Rank | IC50 Mean Rank | 1-18 | VRC07-523 LS | 3BNC117-LS | VRC01 | CH235.12 | PGT121.414 LS | 10-1074 LS | PGDM1400 | CAP256-VRC26.25 LS | PGT151 | VRC34.01 | ACS202 | 10E8v4 |
| 1 | V704_0128_220_RE_NT_pblib001_52 | 10.21 | 1 | 0.023 | 0.006 | 0.026 | 0.04 | 0.021 | 0.018 | 0.059 | 0.008 | 0.002 | 25 | 0.19 | 22.918 | 1.221 |
| 2 | V704_0855_080_RE_NT_pblib001_187 | 11.47 | 2 | 0.012 | 0.003 | 0.009 | 0.013 | 25 | 0.05 | 0.061 | 25 | 25 | 0.007 | 25 | 25 | 0.134 |
| 3 | V704_1180_070_RE_NT_pblib001_87 | 12.59 | 3 | 0.005 | 0.045 | 0.016 | 0.17 | 0.145 | 25 | 25 | 0.188 | 25 | 0.009 | 0.28 | 0.273 | 0.32 |
| 4 | V703_0472_030_RE_con_s | 13.75 | 4 | 0.005 | 0.01 | 25 | 0.021 | 0.05 | 25 | 7.94 | 0.1 | 0.01 | 0.028 | 5.67 | 10 | 0.42 |
| 5 | V703_2304_150_RE_con_s | 13.77 | 5 | 0.06 | 0.02 | 0.06 | 0.1135 | 0.08 | 0.07 | 0.06 | 0.01 | 0.01 | 10 | 0.03 | 10 | 0.36 |
| 6 | V703_2805_080_RE_con_s | 15.12 | 6 | 0.027 | 0.01 | 0.02 | 0.2668 | 0.24 | 25 | 25 | 0.0019 | 0.0004 | 0.06 | 0.23 | 10 | 1.22 |
| 7 | V704_1535_030_RE_NT_pblib001_471 | 15.30 | 7 | 0.052 | 0.068 | 0.019 | 0.088 | 25 | 0.011 | 0.071 | 1.109 | 25 | 0.197 | 0.39 | 0.512 | 0.015 |
| 8 | V704_1528_240_RE_NT_pblib001_17 | 15.69 | 9 | 0.06 | 0.439 | 0.092 | 2.39 | 7.107 | 0.003 | 0.004 | 25 | 25 | 1.418 | 0.04 | 0.041 | 0.243 |
| 9 | V704_0907_130_RE_NT_pblib001_32 | 16.01 | 10 | 0.016 | 0.193 | 0.199 | 0.364 | 15.857 | 0.002 | 0.006 | 25 | 25 | 0.017 | 0.25 | 0.236 | 0.655 |
| 10 | V703_1313_040_RE_con_s | 17.08 | 12 | 0.046 | 0.04 | 0.11 | 0.686 | 10 | 0.01 | 0.06 | 0.004 | 0.0018 | 10 | 10 | 10 | 0.01 |
| 11 | V703_2117_110_RE_sga2A10_s | 20.38 | 17 | 0.17 | 0.04 | 0.11 | 0.2285 | 0.52 | 25 | 25 | 0.0007 | 0.0001 | 10 | 0.81 | 10 | 0.88 |
| 12 | V704_0445_180_RE_NT_pblib001_22 | 26.09 | 25 | 0.012 | 0.063 | 25 | 82.985 | 0.205 | 0.291 | 0.086 | 0.285 | 25 | 0.083 | 25 | 0.08 | 1.49 |
| replaced | V703_0537_110_RE_sga4H1_s | 15.32 | 8 | 0.01 | 0.01 | 0.02 | 0.0899 | 1.14 | 0.01 | 0.08 | 0.09 | 0.01 | 10 | 10 | 10 | 0.68 |
| replaced | V704_2684_181_RE_NT_pblib001_13 | 16.32 | 11 | 0.01 | 0.044 | 0.051 | 0.09 | 25 | 0.006 | 0.019 | 25 | 25 | 25 | 25 | 0.913 | 0.03 |
| | V704_2541_080_RE_NT_pblib001_13 | 18.13 | 13 | 0.037 | 0.056 | 0.032 | 0.305 | 0.288 | 0.026 | 0.008 | 2.163 | 25 | 0.033 | 0.76 | 25 | 0.25 |
| | V704_1775_030_RE_NT_pblib001_135 | 18.69 | 14 | 0.014 | 0.029 | 0.109 | 0.09 | 0.05 | 25 | 25 | 0.002 | 0.028 | 0.112 | 25 | 25 | 0.291 |
| | V703_0566_160_RE_con_s | 19.12 | 15 | 0.04 | 0.09 | 0.2 | 1.3324 | 0.48 | 0.01 | 0.01 | 25 | 25 | 0.007 | 0.08 | 10 | 0.92 |
| | V703_1675_080_RE_con_s-modified | 20.36 | 16 | 0.02 | 0.08 | 0.06 | 0.3446 | 0.15 | 0.98 | 1.42 | 0.01 | 0.0001 | 0.59 | 10 | 4.47 | 18.53 |
| | V703_1471_190_RE_con_s | 20.58 | 18 | 0.05 | 0.08 | 0.06 | 0.1608 | 0.6 | 0.01 | 0.1 | 0.02 | 0.01 | 0.01 | 10 | 10 | 4.92 |
| | V703_0739_110_RE_con_s-modified | 21.21 | 19 | 0.025 | 0.05 | 0.07 | 0.1431 | 0.56 | 0.67 | 0.16 | 3.23 | 0.0009 | 10 | 4.86 | 0.09 | 1.94 |
| | V703_2149_060_RE_sgaB10_s | 21.51 | 20 | 0.0324 | 0.07 | 25 | 0.205 | 10 | 0.11 | 0.07 | 0.0012 | 0.0001 | 9.17 | 10 | 10 | 0.98 |
| | V704_1783_150_RE_NT_pblib001_144 | 22.92 | 21 | 0.006 | 0.064 | 0.036 | 0.135 | 25 | 0.248 | 0.485 | 25 | 25 | 0.06 | 25 | 0.353 | 0.388 |
| | V703_2631_150_RE_sga2F8_s | 23.90 | 22 | 0.02 | 0.08 | 0.07 | 0.4631 | 1.02 | 25 | 25 | 0.0019 | 0.0004 | 10 | 0.97 | 10 | 1.23 |
| | V703_0629_150_RE_pblib002_s | 24.36 | 23 | 0.09 | 0.22 | 0.39 | 0.7194 | 0.2 | 4.72 | 4.4 | 0.42 | 0.05 | 0.0032 | 0.16 | 10 | 4.27 |
| | V704_0026_231_RE_NT_pblib001_3 | 25.62 | 24 | 0.097 | 0.081 | 0.103 | 0.46 | 25 | 0.006 | 0.017 | 0.075 | 25 | 25 | 25 | 25 | 0.07 |
| | V704_1930_170_RE_NT_pblib002_114 | 26.24 | 26 | 0.032 | 0.025 | 0.028 | 0.11 | 0.087 | 25 | 25 | 0.014 | 25 | 25 | 25 | 25 | 1 |
| | V704_0856_240_RE_NT_pblib001_9 | 27.10 | 27 | 0.095 | 0.063 | 0.06 | 0.524 | 0.503 | 0.011 | 0.016 | 25 | 25 | 1.04 | 7.85 | 25 | 0.54 |
| | V703_1848_190_RE_pblib002_s | 27.57 | 28 | 0.0517 | 0.15 | 0.06 | 1.0367 | 0.92 | 25 | 6.74 | 0.01 | 0.01 | 0.06 | 10 | 10 | 0.28 |
| | V704_2788_060_RE_NT_pblib001_96 | 28.86 | 29 | 0.075 | 0.173 | 0.167 | 1.36 | 0.694 | 0.171 | 0.201 | 0.003 | 0.0003 | 2.78 | 25 | 25 | 0.43 |
| | V704_2555_240_RE_NT_pblib001_231 | 28.95 | 30 | 0.092 | 0.052 | 0.075 | 0.424 | 25 | 0.012 | 0.03 | 25 | 25 | 25 | 2.81 | 0.151 | 25 |
| | V703_1764_250_RE_con_s | 28.99 | 31 | 0.03 | 0.1 | 0.1 | 0.2402 | 0.1 | 0.2 | 0.13 | 25 | 25 | 0.18 | 10 | 10 | 1.41 |
| | V703_2018_240_RE_sga6A1_s | 29.25 | 32 | 0.16 | 0.3 | 1.13 | 1.2489 | 6.38 | 0.05 | 0.07 | 0.0031 | 0.0001 | 10 | 10 | 10 | 2.85 |
| | V704_1183_220_RE_NT_pblib001_70 | 29.30 | 33 | 0.041 | 0.347 | 0.065 | 0.39 | 1.185 | 0.032 | 0.051 | 25 | 25 | 3.666 | 25 | 0.228 | 0.495 |
| | V704_1991_230_RE_NT_pblib001_38 | 30.11 | 34 | 0.093 | 0.284 | 0.242 | 1.51 | 0.303 | 25 | 25 | 0.026 | 0.0003 | 0.373 | 25 | 0.128 | 0.5 |
| | V703_1750_140_RE_con_s | 30.23 | 35 | 0.11 | 0.18 | 0.23 | 0.8813 | 1.35 | 0.05 | 0.08 | 25 | 8.59 | 0.05 | 0.34 | 10 | 0.5 |
| | V704_2065_060_RE_NT_pblib001_47 | 30.69 | 36 | 0.069 | 0.146 | 0.047 | 0.22 | 0.052 | 25 | 25 | 25 | 25 | 25 | 25 | 25 | 0.1 |
| | V704_1835_150_RE_NT_pblib001_10 | 31.23 | 37 | 0.027 | 21.596 | 25 | 99 | 25 | 3.127 | 0.095 | 25 | 25 | 0.112 | 0.41 | 0.113 | 0.147 |
| | V704_0575_060_RE_NT_pblib002_16 | 32.08 | 38 | 0.244 | 0.989 | 1.294 | 1.934 | 2.286 | 25 | 25 | 0.051 | 25 | 25 | 0.24 | 0.039 | 0.57 |
| | V704_0944_180_RE_NT_pblib001_76 | 32.33 | 39 | 0.023 | 0.02 | 0.061 | 0.074 | 25 | 25 | 25 | 25 | 25 | 0.367 | 25 | 25 | 25 |
| | V703_1060_080_RE_con_s | 32.84 | 40 | 0.09 | 0.13 | 0.8 | 1.7123 | 1.17 | 25 | 3.63 | 25 | 0.31 | 0.05 | 10 | 10 | 0.08 |
| | V704_2095_130_RE_NT_pblib001_136 | 32.92 | 41 | 0.079 | 0.096 | 0.053 | 0.312 | 0.283 | 2.167 | 0.085 | 25 | 25 | 0.147 | 25 | 25 | 0.51 |
| | V703_0646_051_RE_con_s | 33.47 | 42 | 0.18 | 0.45 | 1.54 | 4.8972 | 10 | 25 | 25 | 0.01 | 0.01 | 0.07 | 0.07 | 10 | 1.56 |
| | V704_0847_030_RE_NT_pblib001_34 | 33.49 | 43 | 6.634 | 0.19 | 25 | 2.048 | 25 | 0.01 | 0.003 | 25 | 25 | 25 | 25 | 25 | 0.221 |
| | V703_1889_100_RE_con_s | 34.72 | 44 | 0.09 | 2.3 | 25 | 25 | 5.78 | 0.09 | 0.23 | 0.01 | 0.0001 | 10 | 10 | 10 | 25 |
| | V704_2448_240_RE_NT_pblib001_288 | 35.69 | 45 | 0.133 | 0.108 | 0.058 | 0.14 | 0.146 | 25 | 25 | 25 | 25 | 25 | 25 | 7.416 | 25 |
| | V703_2539_070_RE_sga6F6_s | 36.03 | 46 | 0.74 | 0.39 | 25 | 1.7655 | 10 | 0.01 | 0.04 | 25 | 25 | 4.57 | 10 | 10 | 0.23 |
| | V704_3000_240_RE_NT_pblib001_8 | 36.05 | 47 | 0.037 | 0.11 | 0.156 | 0.523 | 25 | 25 | 25 | 8.698 | 25 | 25 | 0.28 | 25 | 0.111 |
| | V703_2788_030_RE_con_s | 36.41 | 48 | 1.56 | 0.24 | 0.09 | 0.6812 | 4.9 | 0.3 | 0.09 | 1.2 | 0.01 | 1.84 | 10 | 10 | 5.38 |
| | V704_0496_040_RE_NT_pblib001_37 | 36.41 | 49 | 0.037 | 0.242 | 25 | 4.94 | 1.236 | 0.142 | 0.078 | 0.011 | 0.011 | 25 | 25 | 25 | 0.289 |
| | V704_0372_250_RE_NT_pblib001_7 | 36.55 | 50 | 0.03 | 0.167 | 0.199 | 0.415 | 0.191 | 25 | 25 | 25 | 25 | 25 | 25 | 25 | 0.087 |
| | V703_0279_110_RE_con_s-modified | 36.56 | 51 | 0.08 | 0.25 | 0.44 | 1.391 | 5.34 | 0.77 | 0.49 | 21.61 | 25 | 10 | 10 | 10 | 0.08 |
| | V704_1706_040_RE_NT_pblib001_26 | 37.60 | 52 | 0.2 | 0.058 | 0.164 | 0.489 | 0.186 | 0.457 | 0.301 | 1.986 | 25 | 25 | 25 | 25 | 0.488 |
| | V703_1298_080_RE_pblib002_s | 38.21 | 53 | 0.15 | 1.27 | 25 | 25 | 0.16 | 25 | 25 | 0.09 | 0.01 | 0.1 | 10 | 10 | 0.84 |
| | V703_1915_250_RE_sgaA3_s | 38.62 | 54 | 0.0841 | 0.36 | 1.6 | 0.5291 | 5.6 | 0.32 | 0.84 | 25 | 3.48 | 0.04 | 10 | 10 | 3.83 |
| | V704_2981_150_RE_NT_pblib004_2 | 39.81 | 55 | 0.392 | 0.414 | 0.129 | 2.43 | 0.279 | 1.271 | 0.702 | 0.262 | 25 | 25 | 9.1 | 25 | 0.23 |
| | V704_0886_250_RE_NT_pblib001_10 | 41.06 | 56 | 0.173 | 0.177 | 0.516 | 1.745 | 25 | 0.338 | 0.185 | 25 | 25 | 25 | 25 | 0.498 | 0.171 |
| | V704_0746_760_RE_NT_pblib001_37 | 41.31 | 57 | 0.19 | 0.397 | 0.255 | 0.87 | 5.201 | 0.015 | 0.014 | 25 | 25 | 25 | 25 | 25 | 3.08 |
| | V704_2834_210_RE_NT_pblib002_26 | 41.47 | 58 | 0.056 | 0.199 | 0.059 | 2.039 | 25 | 7.78 | 3.385 | 25 | 25 | 25 | 1.18 | 25 | 0.397 |
| | V704_0644_060_RE_NT_pblib001_216 | 42.02 | 59 | 0.084 | 0.155 | 0.29 | 1.02 | 3.275 | 6.348 | 0.454 | 25 | 25 | 25 | 5.73 | 25 | 0.297 |
| | V704_1429_090_RE_NT_pblib002_75 | 42.07 | 60 | 0.111 | 0.097 | 1.216 | 0.235 | 25 | 25 | 25 | 0.079 | 11.679 | 4.01 | 25 | 25 | 0.537 |
| | V704_2767_070_RE_NT_pblib001_63 | 42.11 | 61 | 0.217 | 0.126 | 0.796 | 2.06 | 1.275 | 0.066 | 0.241 | 25 | 25 | 0.038 | 25 | 25 | 2.03 |
| | V704_0911_150_RE_NT_pblib001_58 | 42.39 | 62 | 0.437 | 0.538 | 0.622 | 1.391 | 10.111 | 0.013 | 0.018 | 25 | 25 | 3.99 | 25 | 25 | 0.869 |
| | V703_1104_100_RE_pblib001_s | 42.47 | 63 | 0.18 | 1.81 | 2.57 | 4.0481 | 10 | 25 | 2.98 | 0.33 | 0.14 | 10 | 10 | 0.19 | 0.92 |
| | V704_3008_040_RE_NT_pblib001_113 | 42.64 | 64 | 0.067 | 0.431 | 0.163 | 3.02 | 25 | 0.017 | 0.178 | 25 | 25 | 25 | 25 | 25 | 0.184 |
| | V703_0203_081_RE_pblib002_s | 42.66 | 65.5 | 0.38 | 0.45 | 25 | 3.8721 | 7.6 | 0.04 | 0.03 | 25 | 25 | 10 | 10 | 10 | 0.67 |
| | V703_0217_050_RE_pblib002_s | 42.66 | 65.5 | 0.38 | 0.45 | 25 | 3.8721 | 7.6 | 0.04 | 0.03 | 25 | 25 | 10 | 10 | 10 | 0.67 |
| | V703_0842_200_RE_con_s | 42.66 | 67 | 10 | 1.44 | 2.13 | 20.8948 | 6.36 | 7.23 | 1.56 | 0.11 | 0.01 | 0.21 | 10 | 10 | 0.73 |
| | V703_0926_070_RE_sga2H2_s | 42.93 | 68 | 2.02 | 0.81 | 2.04 | 7.8027 | 2.34 | 0.04 | 0.1 | 1.04 | 25 | 10 | 10 | 10 | 0.56 |
| | V703_0013_090_RE_con_s | 43.48 | 69 | 0.52 | 0.42 | 25 | 3.4725 | 10 | 7.97 | 18.41 | 0.01 | 0.01 | 10 | 10 | 10 | 1.61 |
| | V704_0513_150_RE_NT_pblib007_2 | 45.17 | 70 | 0.413 | 0.263 | 0.29 | 1.08 | 17.817 | 25 | 25 | 25 | 25 | 0.047 | 1.47 | 25 | 0.98 |
| | V704_2544_140_RE_NT_pblib002_10 | 45.54 | 71 | 0.431 | 0.122 | 0.108 | 0.922 | 4.104 | 0.493 | 0.611 | 25 | 25 | 25 | 25 | 25 | 0.4 |
| | V704_2839_140_RE_NT_pblib001_208 | 46.10 | 72 | 0.116 | 0.183 | 1.426 | 1.58 | 25 | 25 | 25 | 25 | 25 | 25 | 1.77 | 1.486 | 0.63 |
| | V703_1789_230_RE_sga3A3_s | 50.42 | 73 | 0.09 | 0.99 | 25 | 17.8963 | 10 | 0.83 | 0.34 | 25 | 25 | 10 | 10 | 10 | 2.43 |
| | V704_0726_080_RE_NT_pblib001_26 | 52.86 | 74 | 0.199 | 1.02 | 0.537 | 5.636 | 3.445 | 0.36 | 0.734 | 0.421 | 25 | 25 | 25 | 25 | 2.473 |
| | V704_1109_140_RE_NT_pblib001_191 | 58.55 | 75 | 25 | 0.283 | 25 | 3.36 | 25 | 25 | 25 | 0.201 | 25 | 25 | 25 | 25 | 1.25 |

**Table S3. Ranking of the AMP viruses for creating the SHEP-T2 bNab sensitive screening panel.** Note that only 13 of the 15 antibodies tested (see Table S1) were included here. This is because the full neutralization data for the antibody VRC01-23 LS was not available at the time this panel was first designed and pseudovirus evaluation was initiated, and CAP248-2B AMP data indicated that it was not a broadly neutralizing antibody, so both were excluded it from consideration. We began the work to characterize this panel and to share the panel reagents prior to the complete VRC01-23 LS data being obtained, but its data is included in the SHEP-T2 panel data and is shown in Fig. 3. Fortunately, and as expected, the 12 viruses selected for the panel were also very sensitive to VRC01-23 LS, and the two viruses most sensitive to VRC01-23 LS viruses were already included in the panel. The color key for the neutralizing antibody responses is provided in Table S1 and Fig. 3. The IC50 mean rank orders the viruses according to their overall sensitivity to this panel of antibodies. In addition, the 3 lowest antibody scores for each antibody are written in white and underlined, and we required that at least 2 of these 3 were included the SHEP-T2 panel, which is why two of the lower ranked viruses (ranks 8 and 11) were replaced with viruses that were ranked higher and less sensitive overall (ranks 17 and 25), as these viruses were particularly sensitive to the V2 apex antibodies and to FP antibody ACS202.

### Table S4 A.

| Virus Name | Backbone | Host Cell | Tier | Clade | Virus ID# | Panel | SHEP-T2 panel | MPER sensitive | V3 sensitive | V2 sensitive | FP sensitive | CD4bs sensitive |
|---|---|---|---|---|---|---|---|---|---|---|---|---|
| V704_1535_030_RE_NT_pblib001_471 | SG3 Δenv | 293T/17 | 2 | B | AMP322 | SHEP-T2 | Y | Y | N | N | N | Y |
| V704_0128_220_RE_NT_pblib001_52 | SG3 Δenv | 293T/17 | 2 | F1/B | AMP228 | SHEP-T2 | Y | N | N | Y | N | Y |
| V704_0907_130_RE_NT_pblib001_32 | SG3 Δenv | 293T/17 | 2 | B | AMP17 | SHEP-T2 | Y | N | Y | N | Y | N |
| V704_1528_240_RE_NT_pblib001_17 | SG3 Δenv | 293T/17 | 2 | B | AMP253 | SHEP-T2 | Y | N | Y | N | Y | N |
| V704_0855_080_RE_NT_pblib001_187 | SG3 Δenv | 293T/17 | 2 | B | AMP98 | SHEP-T2 | Y | Y | N | N | N | Y |
| V704_1180_070_RE_NT_pblib001_87 | SG3 Δenv | 293T/17 | 2 | A1/B | 11050 | SHEP-T2 | Y | N | N | N | Y | Y |
| V704_0445_180_RE_NT_pblib001_22 | SG3 Δenv | 293T/17 | 2 | B | AMP218 | SHEP-T2 | Y | N | N | N | Y | N |
| V703_2117_110_RE_sga2A10_s | SG3 Δenv | 293T/17 | 2 | C | AMP333 | SHEP-T2 | Y | N | N | Y | N | N |
| V703_2117_110_RE_sga2A10_s | SG3 Δenv | 293T/17 | 2 | C | 11105 | SHEP-T2 | Y | N | N | Y | Y | Y |
| V703_1313_040_RE_con_s | SG3 Δenv | 293T/17 | 2 | C | 11065 | SHEP-T2 | Y | Y | Y | Y | N | N |
| V703_2304_150_RE_con_s | SG3 Δenv | 293T/17 | 2 | C | AMP328 | SHEP-T2 | Y | N | N | N | Y | N |
| V703_0472_030_RE_con_s | SG3 Δenv | 293T/17 | 2 | C | 11072 | SHEP-T2 | Y | N | N | N | N | Y |
| V704_2684_181_RE_NT_pblib001_13 | SG3 Δenv | 293T/17 | 2 | B | AMP205 | V3 MPER | N | Y | Y | N | N | N |
| V704_0026_231_RE_NT_pblib001_3 | SG3 Δenv | 293T/17 | 2 | B | AMP245 | V3 MPER | N | Y | Y | N | N | N |
| V703_1060_080_RE_con_s | SG3 Δenv | 293T/17 | 2 | C | AMP38 | MPER | N | Y | N | N | N | N |
| V703_0279_110_RE_con_s-modified | SG3 Δenv | 293T/17 | 2 | C | AMP339 | MPER | N | Y | N | N | N | N |
| V704_0372_250_RE_NT_pblib001_7 | SG3 Δenv | 293T/17 | 2 | B | AMP312 | MPER | N | Y | N | N | N | N |
| V704_0847_030_RE_NT_pblib001_34 | SG3 Δenv | 293T/17 | 2 | B | 11132 | V3 | N | N | Y | N | N | N |
| V703_0566_160_RE_con_s | SG3 Δenv | 293T/17 | 2 | C | AMP337 | V3 FP | N | N | Y | N | Y | N |
| V704_2541_080_RE_NT_pblib001_13 | SG3 Δenv | 293T/17 | 2 | B | AMP201 | V3 | N | N | Y | N | N | N |
| V703_0629_150_RE_pblib002_s | SG3 Δenv | 293T/17 | 2 | C | AMP345 | FP | N | N | N | N | Y | N |
| V704_0575_060_RE_NT_pblib002_16 | SG3 Δenv | 293T/17 | 2 | B | AMP181 | FP | N | N | N | N | Y | N |
| V703_0646_051_RE_con_s | SG3 Δenv | 293T/17 | 2 | C | AMP350 | FP | N | N | N | N | Y | N |
| V704_1835_150_RE_NT_pblib001_10 | SG3 Δenv | 293T/17 | 2 | B | AMP192 | FP | N | N | N | N | Y | N |
| V703_0739_110_RE_con_s-modified | SG3 Δenv | 293T/17 | 2 | C | AMP357 | FP | N | N | N | N | Y | N |
| V703_2149_060_RE_sgaB10_s | SG3 Δenv | 293T/17 | 2 | C | 11149 | V2 | N | N | N | Y | N | N |
| V703_2018_240_RE_sga6A1_s | SG3 Δenv | 293T/17 | 2 | C | AMP341 | V2 | N | N | N | Y | N | N |
| V703_2631_150_RE_sga2F8_s | SG3 Δenv | 293T/17 | IB | C | AMP343 | V2 | N | N | N | N | N | N |
| V703_1675_080_RE_con_s-modified | SG3 Δenv | 293T/17 | 2 | C | 11112 | V2 | N | N | N | Y | N | N |
| V703_0537_110_RE_sga4H1_s | SG3 Δenv | 293T/17 | 2 | C | AMP348 | CD4bs | N | N | N | N | N | Y |
| V704_1775_030_RE_NT_pblib001_135 | SG3 Δenv | 293T/17 | 2 | F1 | AMP93 | CD4bs V2 | N | N | N | Y | N | Y |
| V704_1783_150_RE_NT_pblib001_144 | SG3 Δenv | 293T/17 | 2 | B | AMP167 | CD4bs | N | N | N | N | N | Y |

### Table S4 B.

| | VRC26UCA_4A/293I | | AbPG9_RUA/293I | | AbPG16_RUA/293I | | glPGT145/293I | | PGT121tkUCA.V2/293I | | DH270UCA3_G1.4A/293I | | Antibody |
|---|---|---|---|---|---|---|---|---|---|---|---|---|---|
| | 191022PPF | | 200221PPF | | 200224PPF | | 19121PPF | | 200225PPF | | 191107PPF | | Lot |
| | V2-apex | | V2-apex | | V2-apex | | Apex | | V3-glycan | | V3-glycan | | Epitope |
| Virus Name | ID50 | ID80 | ID50 | ID80 | ID50 | ID80 | ID50 | ID80 | ID50 | ID80 | ID50 | ID80 | Measure |
| V704_1535_030_RE_NT_pblib001_471 | >50 | >50 | >50 | >50 | >50 | >50 | >50 | >50 | >50 | >50 | >50 | >50 | |
| V704_0128_220_RE_NT_pblib001_52 | >50 | >50 | >50 | >50 | >50 | >50 | >50 | >50 | >50 | >50 | >50 | >50 | |
| V704_0907_130_RE_NT_pblib001_32 | >50 | >50 | >50 | >50 | >50 | >50 | >50 | >50 | >50 | >50 | >50 | >50 | |
| V704_1528_240_RE_NT_pblib001_17 | >50 | >50 | >50 | >50 | >50 | >50 | >50 | >50 | >50 | >50 | >50 | >50 | |
| V704_0855_080_RE_NT_pblib001_187 | >50 | >50 | >50 | >50 | >50 | >50 | >50 | >50 | >50 | >50 | >50 | >50 | |
| V704_1180_070_RE_NT_pblib001_87 | >50 | >50 | >50 | >50 | >50 | >50 | >50 | >50 | >50 | >50 | >50 | >50 | |
| V704_0445_180_RE_NT_pblib001_22 | >50 | >50 | >50 | >50 | >50 | >50 | >50 | >50 | >50 | >50 | >50 | >50 | |
| V703_2117_110_RE_sga2A10_s | >50 | >50 | >50 | >50 | >50 | >50 | >50 | >50 | >50 | >50 | >50 | >50 | |
| V703_2117_110_RE_sga2A10_s | >50 | >50 | >50 | >50 | >50 | >50 | >50 | >50 | >50 | >50 | >50 | >50 | |
| V703_1313_040_RE_con_s | >50 | >50 | >50 | >50 | >50 | >50 | >50 | >50 | >50 | >50 | >50 | >50 | |
| V703_2304_150_RE_con_s | >50 | >50 | >50 | >50 | >50 | >50 | >50 | >50 | >50 | >50 | >50 | >50 | |
| V703_0472_030_RE_con_s | >50 | >50 | >50 | >50 | >50 | >50 | >50 | >50 | >50 | >50 | >50 | >50 | |
| V704_2684_181_RE_NT_pblib001_13 | >50 | >50 | >50 | >50 | >50 | >50 | >50 | >50 | >50 | >50 | >50 | >50 | |
| V704_0026_231_RE_NT_pblib001_3 | >50 | >50 | >50 | >50 | >50 | >50 | >50 | >50 | >50 | >50 | >50 | >50 | |
| V703_1060_080_RE_con_s | >50 | >50 | >50 | >50 | >50 | >50 | >50 | >50 | >50 | >50 | >50 | >50 | |
| V703_0279_110_RE_con_s-modified | >50 | >50 | >50 | >50 | >50 | >50 | >50 | >50 | >50 | >50 | >50 | >50 | |
| V704_0372_250_RE_NT_pblib001_7 | >50 | >50 | >50 | >50 | >50 | >50 | >50 | >50 | >50 | >50 | >50 | >50 | |
| V704_0847_030_RE_NT_pblib001_34 | >50 | >50 | >50 | >50 | >50 | >50 | >50 | >50 | >50 | >50 | >50 | >50 | |
| V703_0566_160_RE_con_s | >50 | >50 | >50 | >50 | >50 | >50 | >50 | >50 | >50 | >50 | >50 | >50 | |
| V704_2541_080_RE_NT_pblib001_13 | >50 | >50 | >50 | >50 | >50 | >50 | >50 | >50 | >50 | >50 | >50 | >50 | |
| V703_0629_150_RE_pblib002_s | >50 | >50 | >50 | >50 | >50 | >50 | >50 | >50 | >50 | >50 | >50 | >50 | |
| V704_0575_060_RE_NT_pblib002_16 | >50 | >50 | >50 | >50 | >50 | >50 | >50 | >50 | >50 | >50 | >50 | >50 | |
| V703_0646_051_RE_con_s | >50 | >50 | >50 | >50 | >50 | >50 | >50 | >50 | >50 | >50 | >50 | >50 | |
| V704_1835_150_RE_NT_pblib001_10 | >50 | >50 | >50 | >50 | >50 | >50 | >50 | >50 | >50 | >50 | >50 | >50 | |
| V703_0739_110_RE_con_s-modified | >50 | >50 | >50 | >50 | >50 | >50 | >50 | >50 | >50 | >50 | >50 | >50 | |
| V703_2149_060_RE_sgaB10_s | >50 | >50 | >50 | >50 | >50 | >50 | >50 | >50 | >50 | >50 | >50 | >50 | |
| V703_2018_240_RE_sga6A1_s | >50 | >50 | >50 | >50 | >50 | >50 | >50 | >50 | >50 | >50 | >50 | >50 | |
| V703_2631_150_RE_sga2F8_s | >50 | >50 | >50 | >50 | >50 | >50 | >50 | >50 | >50 | >50 | >50 | >50 | |
| V703_1675_080_RE_con_s-modified | >50 | >50 | >50 | >50 | >50 | >50 | >50 | >50 | >50 | >50 | >50 | >50 | |
| V703_0537_110_RE_sga4H1_s | >50 | >50 | >50 | >50 | >50 | >50 | >50 | >50 | >50 | >50 | >50 | >50 | |
| V704_1775_030_RE_NT_pblib001_135 | >50 | >50 | >50 | >50 | >50 | >50 | >50 | >50 | >50 | >50 | >50 | >50 | |
| V704_1783_150_RE_NT_pblib001_144 | >50 | >50 | >50 | >50 | >50 | >50 | >50 | >50 | >50 | >50 | >50 | >50 | |

### Table S4 B, continued.

| | PGT128_gl/293I | | VRC03gHvgLv/293I | | VRC04gHvgLv/293I | | VRC07gHvgLv/293I | | VRC18bgHvgLv | | VRC20gHvgLv | | Antibody |
|---|---|---|---|---|---|---|---|---|---|---|---|---|---|
| | 200226PPF | | 191106PPF | | 191023PPF | | 191024PPF | | 191114PPF | | 191029PPF | | Lot |
| | V3-glycan | | CD4bs | | CD4bs | | CD4bs | | CD4bs | | CD4bs | | Epitope |
| Virus Name | ID50 | ID80 | ID50 | ID80 | ID50 | ID80 | ID50 | ID80 | ID50 | ID80 | ID50 | ID80 | Measure |
| V704_1535_030_RE_NT_pblib001_471 | >50 | >50 | >50 | >50 | >50 | >50 | >50 | >50 | >50 | >50 | >50 | >50 | |
| V704_0128_220_RE_NT_pblib001_52 | >50 | >50 | >50 | >50 | >50 | >50 | >50 | >50 | >50 | >50 | >50 | >50 | |
| V704_0907_130_RE_NT_pblib001_32 | >50 | >50 | >50 | >50 | >50 | >50 | >50 | >50 | >50 | >50 | >50 | >50 | |
| V704_1528_240_RE_NT_pblib001_17 | >50 | >50 | >50 | >50 | >50 | >50 | >50 | >50 | >50 | >50 | >50 | >50 | |
| V704_0855_080_RE_NT_pblib001_187 | >50 | >50 | >50 | >50 | >50 | >50 | >50 | >50 | >50 | >50 | >50 | >50 | |
| V704_1180_070_RE_NT_pblib001_87 | >50 | >50 | >50 | >50 | >50 | >50 | >50 | >50 | >50 | >50 | >50 | >50 | |
| V704_0445_180_RE_NT_pblib001_22 | >50 | >50 | >50 | >50 | >50 | >50 | >50 | >50 | >50 | >50 | >50 | >50 | |
| V703_2117_110_RE_sga2A10_s | >50 | >50 | >50 | >50 | >50 | >50 | >50 | >50 | >50 | >50 | >50 | >50 | |
| V703_2117_110_RE_sga2A10_s | >50 | >50 | >50 | >50 | >50 | >50 | >50 | >50 | >50 | >50 | >50 | >50 | |
| V703_1313_040_RE_con_s | >50 | >50 | >50 | >50 | >50 | >50 | >50 | >50 | >50 | >50 | >50 | >50 | |
| V703_2304_150_RE_con_s | >50 | >50 | >50 | >50 | >50 | >50 | >50 | >50 | >50 | >50 | >50 | >50 | |
| V703_0472_030_RE_con_s | >50 | >50 | >50 | >50 | >50 | >50 | >50 | >50 | >50 | >50 | >50 | >50 | |
| V704_2684_181_RE_NT_pblib001_13 | >50 | >50 | >50 | >50 | >50 | >50 | >50 | >50 | >50 | >50 | >50 | >50 | |
| V704_0026_231_RE_NT_pblib001_3 | >50 | >50 | >50 | >50 | >50 | >50 | >50 | >50 | >50 | >50 | >50 | >50 | |
| V703_1060_080_RE_con_s | >50 | >50 | >50 | >50 | >50 | >50 | >50 | >50 | >50 | >50 | >50 | >50 | |
| V703_0279_110_RE_con_s-modified | >50 | >50 | >50 | >50 | >50 | >50 | >50 | >50 | >50 | >50 | >50 | >50 | |
| V704_0372_250_RE_NT_pblib001_7 | >50 | >50 | >50 | >50 | >50 | >50 | >50 | >50 | >50 | >50 | >50 | >50 | |
| V704_0847_030_RE_NT_pblib001_34 | >50 | >50 | >50 | >50 | >50 | >50 | >50 | >50 | >50 | >50 | >50 | >50 | |
| V703_0566_160_RE_con_s | >50 | >50 | >50 | >50 | >50 | >50 | >50 | >50 | >50 | >50 | >50 | >50 | |
| V704_2541_080_RE_NT_pblib001_13 | >50 | >50 | >50 | >50 | >50 | >50 | >50 | >50 | >50 | >50 | >50 | >50 | |
| V703_0629_150_RE_pblib002_s | >50 | >50 | >50 | >50 | >50 | >50 | >50 | >50 | >50 | >50 | >50 | >50 | |
| V704_0575_060_RE_NT_pblib002_16 | >50 | >50 | >50 | >50 | >50 | >50 | >50 | >50 | >50 | >50 | >50 | >50 | |
| V703_0646_051_RE_con_s | >50 | >50 | >50 | >50 | >50 | >50 | >50 | >50 | >50 | >50 | >50 | >50 | |
| V704_1835_150_RE_NT_pblib001_10 | >50 | >50 | >50 | >50 | >50 | >50 | >50 | >50 | >50 | >50 | >50 | >50 | |
| V703_0739_110_RE_con_s-modified | >50 | >50 | >50 | >50 | >50 | >50 | >50 | >50 | >50 | >50 | >50 | >50 | |
| V703_2149_060_RE_sgaB10_s | >50 | >50 | >50 | >50 | >50 | >50 | >50 | >50 | >50 | >50 | >50 | >50 | |
| V703_2018_240_RE_sga6A1_s | >50 | >50 | >50 | >50 | >50 | >50 | >50 | >50 | >50 | >50 | >50 | >50 | |
| V703_2631_150_RE_sga2F8_s | >50 | >50 | >50 | >50 | >50 | >50 | >50 | >50 | >50 | >50 | >50 | >50 | |
| V703_1675_080_RE_con_s-modified | >50 | >50 | >50 | >50 | >50 | >50 | >50 | >50 | >50 | >50 | >50 | >50 | |
| V703_0537_110_RE_sga4H1_s | >50 | >50 | >50 | >50 | >50 | >50 | >50 | >50 | >50 | >50 | >50 | >50 | |
| V704_1775_030_RE_NT_pblib001_135 | >50 | >50 | >50 | >50 | >50 | >50 | >50 | >50 | >50 | >50 | >50 | >50 | |
| V704_1783_150_RE_NT_pblib001_144 | >50 | >50 | >50 | >50 | >50 | >50 | >50 | >50 | >50 | >50 | >50 | >50 | |

| | VRC23.gl/293I | | RC01_UCAtk_4A/293 | | CH3X_UCAI (gl-CH3 | | CH103_UCA_4A/293I | | 12A12gl/293I | | glPGV04/293I | | Antibody |
|---|---|---|---|---|---|---|---|---|---|---|---|---|---|
| | 191031PPF | | 191120PPF-1 | | 191112PPF | | 191101PPF | | 191030PPF | | 191119PPF | | Lot |
| | CD4bs | | CD4bs | | CD4bs | | CD4bs | | CD4bs | | CD4bs | | Epitope |
| Virus Name | ID50 | ID80 | ID50 | ID80 | ID50 | ID80 | ID50 | ID80 | ID50 | ID80 | ID50 | ID80 | Measure |
| V704_1535_030_RE_NT_pblib001_471 | >50 | >50 | >50 | >50 | >50 | >50 | >50 | >50 | >50 | >50 | >50 | >50 | |
| V704_0128_220_RE_NT_pblib001_52 | >50 | >50 | >50 | >50 | >50 | >50 | >50 | >50 | >50 | >50 | >50 | >50 | |
| V704_0907_130_RE_NT_pblib001_32 | >50 | >50 | >50 | >50 | >50 | >50 | >50 | >50 | >50 | >50 | >50 | >50 | |
| V704_1528_240_RE_NT_pblib001_17 | >50 | >50 | >50 | >50 | >50 | >50 | >50 | >50 | >50 | >50 | >50 | >50 | |
| V704_0855_080_RE_NT_pblib001_187 | >50 | >50 | >50 | >50 | >50 | >50 | >50 | >50 | >50 | >50 | >50 | >50 | |
| V704_1180_070_RE_NT_pblib001_87 | >50 | >50 | >50 | >50 | >50 | >50 | >50 | >50 | >50 | >50 | >50 | >50 | |
| V704_0445_180_RE_NT_pblib001_22 | >50 | >50 | >50 | >50 | >50 | >50 | >50 | >50 | >50 | >50 | >50 | >50 | |
| V703_2117_110_RE_sga2A10_s | >50 | >50 | >50 | >50 | >50 | >50 | >50 | >50 | >50 | >50 | >50 | >50 | |
| V703_2117_110_RE_sga2A10_s | >50 | >50 | >50 | >50 | >50 | >50 | >50 | >50 | >50 | >50 | >50 | >50 | |
| V703_1313_040_RE_con_s | >50 | >50 | >50 | >50 | >50 | >50 | >50 | >50 | >50 | >50 | >50 | >50 | |
| V703_2304_150_RE_con_s | >50 | >50 | >50 | >50 | >50 | >50 | >50 | >50 | >50 | >50 | >50 | >50 | |
| V703_0472_030_RE_con_s | >50 | >50 | >50 | >50 | >50 | >50 | >50 | >50 | >50 | >50 | >50 | >50 | |
| V704_2684_181_RE_NT_pblib001_13 | >50 | >50 | >50 | >50 | >50 | >50 | >50 | >50 | >50 | >50 | >50 | >50 | |
| V704_0026_231_RE_NT_pblib001_3 | >50 | >50 | >50 | >50 | >50 | >50 | >50 | >50 | >50 | >50 | >50 | >50 | |
| V703_1060_080_RE_con_s | >50 | >50 | >50 | >50 | >50 | >50 | >50 | >50 | >50 | >50 | >50 | >50 | |
| V703_0279_110_RE_con_s-modified | >50 | >50 | >50 | >50 | *failed* | *failed* | >50 | >50 | >50 | >50 | >50 | >50 | |
| V704_0372_250_RE_NT_pblib001_7 | >50 | >50 | >50 | >50 | >50 | >50 | >50 | >50 | >50 | >50 | >50 | >50 | |
| V704_0847_030_RE_NT_pblib001_34 | >50 | >50 | >50 | >50 | >50 | >50 | >50 | >50 | >50 | >50 | >50 | >50 | |
| V703_0566_160_RE_con_s | >50 | >50 | >50 | >50 | >50 | >50 | >50 | >50 | >50 | >50 | >50 | >50 | |
| V704_2541_080_RE_NT_pblib001_13 | >50 | >50 | >50 | >50 | >50 | >50 | >50 | >50 | >50 | >50 | >50 | >50 | |
| V703_0629_150_RE_pblib002_s | >50 | >50 | >50 | >50 | >50 | >50 | >50 | >50 | >50 | >50 | >50 | >50 | |
| V704_0575_060_RE_NT_pblib002_16 | >50 | >50 | >50 | >50 | >50 | >50 | >50 | >50 | >50 | >50 | >50 | >50 | |
| V703_0646_051_RE_con_s | >50 | >50 | >50 | >50 | >50 | >50 | >50 | >50 | >50 | >50 | >50 | >50 | |
| V704_1835_150_RE_NT_pblib001_10 | >50 | >50 | >50 | >50 | >50 | >50 | >50 | >50 | >50 | >50 | >50 | >50 | |
| V703_0739_110_RE_con_s-modified | >50 | >50 | >50 | >50 | >50 | >50 | >50 | >50 | >50 | >50 | >50 | >50 | |
| V703_2149_060_RE_sgaB10_s | >50 | >50 | >50 | >50 | >50 | >50 | >50 | >50 | >50 | >50 | >50 | >50 | |
| V703_2018_240_RE_sga6A1_s | >50 | >50 | >50 | >50 | >50 | >50 | >50 | >50 | >50 | >50 | >50 | >50 | |
| V703_2631_150_RE_sga2F8_s | >50 | >50 | >50 | >50 | >50 | >50 | >50 | >50 | >50 | >50 | >50 | >50 | |
| V703_1675_080_RE_con_s-modified | >50 | >50 | >50 | >50 | >50 | >50 | >50 | >50 | >50 | >50 | >50 | >50 | |
| V703_0537_110_RE_sga4H1_s | >50 | >50 | >50 | >50 | >50 | >50 | >50 | >50 | >50 | >50 | >50 | >50 | |
| V704_1775_030_RE_NT_pblib001_135 | >50 | >50 | >50 | >50 | >50 | >50 | >50 | >50 | >50 | >50 | >50 | >50 | |
| V704_1783_150_RE_NT_pblib001_144 | >50 | >50 | >50 | >50 | >50 | >50 | >50 | >50 | >50 | >50 | >50 | >50 | |

**Table S4 B, continued.**

| Virus Name | gl1NC9/293i | | gl3BNC60/293i | | glNIH45-46/293i | | glIOMA/293i | | VRC01gHvgLv/293i | | DH235UCAtk_v3_4A/293i | | Antibody |
|---|---|---|---|---|---|---|---|---|---|---|---|---|---|
| | 200227PPF | | 200115PPF | | 200228PPF | | 5Mar202PPFJCA-1 | | 191105PPF | | 200114PPF | | Lot |
| | CD4bs | | CD4bs | | CD4bs | | CD4bs | | CD4bs | | CD4bs | | Epitope |
| | ID50 | ID80 | ID50 | ID80 | ID50 | ID80 | ID50 | ID80 | ID50 | ID80 | ID50 | ID80 | Measure |
| V704_1535_030_RE_NT_pblib001_471 | >50 | >50 | >50 | >50 | >50 | >50 | **32** | >50 | >50 | >50 | >50 | >50 | |
| V704_0128_220_RE_NT_pblib001_52 | >50 | >50 | >50 | >50 | >50 | >50 | **34** | >50 | >50 | >50 | >50 | >50 | |
| V704_0907_130_RE_NT_pblib001_32 | >50 | >50 | >50 | >50 | >50 | >50 | **33** | >50 | >50 | >50 | >50 | >50 | |
| V704_1528_240_RE_NT_pblib001_17 | >50 | >50 | >50 | >50 | >50 | >50 | **22** | >50 | >50 | >50 | >50 | >50 | |
| V704_0855_080_RE_NT_pblib001_187 | >50 | >50 | >50 | >50 | >50 | >50 | >50 | >50 | >50 | >50 | >50 | >50 | |
| V704_1180_070_RE_NT_pblib001_87 | >50 | >50 | >50 | >50 | >50 | >50 | >50 | >50 | >50 | >50 | >50 | >50 | |
| V704_0445_180_RE_NT_pblib001_22 | >50 | >50 | >50 | >50 | >50 | >50 | >50 | >50 | >50 | >50 | >50 | >50 | |
| V703_2117_110_RE_sga2A10_s | >50 | >50 | >50 | >50 | >50 | >50 | >50 | >50 | >50 | >50 | >50 | >50 | |
| V703_2117_110_RE_sga2A10_s | >50 | >50 | >50 | >50 | >50 | >50 | **27** | >50 | >50 | >50 | >50 | >50 | |
| V703_1313_040_RE_con_s | >50 | >50 | >50 | >50 | >50 | >50 | **37** | >50 | >50 | >50 | >50 | >50 | |
| V703_2304_150_RE_con_s | >50 | >50 | >50 | >50 | >50 | >50 | >50 | >50 | >50 | >50 | >50 | >50 | |
| V703_0472_030_RE_con_s | >50 | >50 | >50 | >50 | >50 | >50 | >50 | >50 | >50 | >50 | >50 | >50 | |
| V704_2684_181_RE_NT_pblib001_13 | >50 | >50 | >50 | >50 | >50 | >50 | >50 | >50 | >50 | >50 | >50 | >50 | |
| V704_0026_231_RE_NT_pblib001_3 | >50 | >50 | >50 | >50 | >50 | >50 | >50 | >50 | >50 | >50 | >50 | >50 | |
| V703_1060_080_RE_con_s | >50 | >50 | >50 | >50 | >50 | >50 | >50 | >50 | >50 | >50 | >50 | >50 | |
| V703_0279_110_RE_con_s-modified | >50 | >50 | >50 | >50 | >50 | >50 | >50 | >50 | >50 | >50 | >50 | >50 | |
| V704_0372_250_RE_NT_pblib001_7 | >50 | >50 | >50 | >50 | >50 | >50 | >50 | >50 | >50 | >50 | >50 | >50 | |
| V704_0847_030_RE_NT_pblib001_34 | >50 | >50 | >50 | >50 | >50 | >50 | >50 | >50 | >50 | >50 | >50 | >50 | |
| V703_0566_160_RE_con_s | >50 | >50 | >50 | >50 | >50 | >50 | >50 | >50 | >50 | >50 | >50 | >50 | |
| V704_2541_080_RE_NT_pblib001_13 | >50 | >50 | >50 | >50 | >50 | >50 | >50 | >50 | >50 | >50 | >50 | >50 | |
| V703_0629_150_RE_pblib002_s | >50 | >50 | >50 | >50 | >50 | >50 | >50 | >50 | >50 | >50 | >50 | >50 | |
| V704_0575_060_RE_NT_pblib002_16 | >50 | >50 | >50 | >50 | >50 | >50 | >50 | >50 | >50 | >50 | >50 | >50 | |
| V703_0646_051_RE_con_s | >50 | >50 | >50 | >50 | >50 | >50 | >50 | >50 | >50 | >50 | >50 | >50 | |
| V704_1835_150_RE_NT_pblib001_10 | >50 | >50 | >50 | >50 | >50 | >50 | >50 | >50 | >50 | >50 | >50 | >50 | |
| V703_0739_110_RE_con_s-modified | >50 | >50 | >50 | >50 | >50 | >50 | >50 | >50 | >50 | >50 | >50 | >50 | |
| V703_2149_060_RE_sgaB10_s | >50 | >50 | >50 | >50 | >50 | >50 | >50 | >50 | >50 | >50 | >50 | >50 | |
| V703_2018_240_RE_sga6A1_s | >50 | >50 | >50 | >50 | >50 | >50 | >50 | >50 | >50 | >50 | >50 | >50 | |
| V703_2631_150_RE_sga2F8_s | >50 | >50 | >50 | >50 | >50 | >50 | >50 | >50 | >50 | >50 | >50 | >50 | |
| V703_1675_080_RE_con_s-modified | >50 | >50 | >50 | >50 | >50 | >50 | >50 | >50 | >50 | >50 | >50 | >50 | |
| V703_0537_110_RE_sga4H1_s | >50 | >50 | >50 | >50 | >50 | >50 | >50 | >50 | >50 | >50 | >50 | >50 | |
| V704_1775_030_RE_NT_pblib001_135 | >50 | >50 | >50 | >50 | >50 | >50 | >50 | >50 | >50 | >50 | >50 | >50 | |
| V704_1783_150_RE_NT_pblib001_144 | >50 | >50 | >50 | >50 | >50 | >50 | >50 | >50 | >50 | >50 | >50 | >50 | |

| Virus Name | CH01-RUA3_4A/293i | | glPGV19 | | Antibody |
|---|---|---|---|---|---|
| | 200305PPF | | | | Lot |
| | V1/V2 | | CD4bs | | Epitope |
| | ID50 | ID80 | ID50 | ID80 | Measure |
| V704_1535_030_RE_NT_pblib001_471 | >50 | >50 | >50 | >50 | |
| V704_0128_220_RE_NT_pblib001_52 | >50 | >50 | >50 | >50 | |
| V704_0907_130_RE_NT_pblib001_32 | >50 | >50 | >50 | >50 | |
| V704_1528_240_RE_NT_pblib001_17 | >50 | >50 | >50 | >50 | |
| V704_0855_080_RE_NT_pblib001_187 | >50 | >50 | >50 | >50 | |
| V704_1180_070_RE_NT_pblib001_87 | >50 | >50 | >50 | >50 | |
| V704_0445_180_RE_NT_pblib001_22 | >50 | >50 | >50 | >50 | |
| V703_2117_110_RE_sga2A10_s | >50 | >50 | >50 | >50 | |
| V703_2117_110_RE_sga2A10_s | >50 | >50 | >50 | >50 | |
| V703_1313_040_RE_con_s | >50 | >50 | >50 | >50 | |
| V703_2304_150_RE_con_s | >50 | >50 | >50 | >50 | |
| V703_0472_030_RE_con_s | >50 | >50 | >50 | >50 | |
| V704_2684_181_RE_NT_pblib001_13 | >50 | >50 | >50 | >50 | |
| V704_0026_231_RE_NT_pblib001_3 | >50 | >50 | >50 | >50 | |
| V703_1060_080_RE_con_s | >50 | >50 | >50 | >50 | |
| V703_0279_110_RE_con_s-modified | >50 | >50 | >50 | >50 | |
| V704_0372_250_RE_NT_pblib001_7 | >50 | >50 | >50 | >50 | |
| V704_0847_030_RE_NT_pblib001_34 | >50 | >50 | >50 | >50 | |
| V703_0566_160_RE_con_s | >50 | >50 | >50 | >50 | |
| V704_2541_080_RE_NT_pblib001_13 | >50 | >50 | >50 | >50 | |
| V703_0629_150_RE_pblib002_s | >50 | >50 | >50 | >50 | |
| V704_0575_060_RE_NT_pblib002_16 | >50 | >50 | >50 | >50 | |
| V703_0646_051_RE_con_s | >50 | >50 | >50 | >50 | |
| V704_1835_150_RE_NT_pblib001_10 | >50 | >50 | >50 | >50 | |
| V703_0739_110_RE_con_s-modified | >50 | >50 | >50 | >50 | |
| V703_2149_060_RE_sgaB10_s | >50 | >50 | >50 | >50 | |
| V703_2018_240_RE_sga6A1_s | >50 | >50 | >50 | >50 | |
| V703_2631_150_RE_sga2F8_s | >50 | >50 | >50 | >50 | |
| V703_1675_080_RE_con_s-modified | >50 | >50 | >50 | >50 | |
| V703_0537_110_RE_sga4H1_s | >50 | >50 | *failed* | *failed* | |
| V704_1775_030_RE_NT_pblib001_135 | >50 | >50 | >50 | >50 | |
| V704_1783_150_RE_NT_pblib001_144 | >50 | >50 | >50 | >50 | |

**Table S4.** Twenty six germline precursors antibodies that ultimately give rise to bnAbs show no IC50 against any of the 40 viruses included in the SHEP-T2 and class-specific antibody panels. **A.** Virus details and panel inclusion. **B.** IC50 and IC80 responses for each of the 26 antibodies tested; there was no detectable activity against any of the AMP viruses.

### Table S5 A and B

#### Table S5 A.

| Sequence Name HVTN 704 | HIV-1 M Env subtype | PGDM1400 | CAP256-VRC26.25 LS |
|---|---|---|---|
| V704_1991_230_RE_NT_pblib001_38 | B | 0.026 | 0.0003 |
| V704_1991_230_RE_NT_pblib002_19 | B | 0.052 | 0.0003 |
| V704_0372_250_RE_NT_pblib002_6 | B | >25 | 2.114 |
| V704_2981_150_RE_NT_pblib001_33 | B | 0.022 | >25 |
| V704_0026_231_RE_NT_pblib001_3 | B | 0.075 | >25 |
| V704_2544_150_RE_NT_pblib001_33 | B | 0.101 | >25 |
| V704_2541_080_RE_NT_pblib042_1 | B | 0.218 | >25 |
| V704_2981_150_RE_NT_pblib004_2 | B | 0.262 | >25 |
| V704_0445_180_RE_NT_pblib001_22 | B | 0.285 | >25 |
| V704_0726_080_RE_NT_pblib001_26 | B | 0.421 | >25 |
| V704_1535_030_RE_NT_pblib001_471 | B | 1.109 | >25 |
| V704_1706_040_RE_NT_pblib001_26 | B | 1.986 | >25 |
| V704_2541_080_RE_NT_pblib001_13 | B | 2.163 | >25 |
| V704_2541_080_RE_NT_pblib003_5 | B | 3.382 | >25 |
| V704_3000_240_RE_NT_pblib001_8 | B | 8.698 | >25 |
| V704_2544_140_RE_NT_pblib001_11 | B | 13.864 | >25 |
| V704_3000_240_RE_NT_pblib019_1 | B | 16.221 | >25 |
| V704_0513_150_RE_NT_pblib007_2 | B | >25 | >25 |
| V704_0513_150_RE_NT_pblib002_10 | B | >25 | >25 |
| V704_0513_150_RE_NT_pblib001_34 | B | >25 | >25 |
| V704_0746_760_RE_NT_pblib001_37 | B | >25 | >25 |
| V704_0746_760_RE_NT_pblib003_34 | B | >25 | >25 |
| V704_0746_760_RE_NT_pblib002_36 | B | >25 | >25 |
| V704_0847_030_RE_NT_pblib001_34 | B | >25 | >25 |
| V704_0847_030_RE_NT_pblib002_23 | B | >25 | >25 |
| V704_0855_080_RE_NT_pblib001_187 | B | >25 | >25 |
| V704_0856_240_RE_NT_pblib001_9 | B | >25 | >25 |
| V704_0886_250_RE_NT_pblib001_10 | B | >25 | >25 |
| V704_0907_130_RE_NT_pblib001_32 | B | >25 | >25 |
| V704_0911_150_RE_NT_pblib001_58 | B | >25 | >25 |
| V704_0944_180_RE_NT_pblib001_76 | B | >25 | >25 |
| V704_1528_240_RE_NT_pblib001_17 | B | >25 | >25 |
| V704_1783_150_RE_NT_pblib001_144 | B | >25 | >25 |
| V704_1835_150_RE_NT_pblib001_10 | B | >25 | >25 |
| V704_1835_150_RE_NT_pblib002_9 | B | >25 | >25 |
| V704_2065_060_RE_NT_pblib001_47 | B | >25 | >25 |
| V704_2065_060_RE_NT_pblib002_42 | B | >25 | >25 |
| V704_2095_130_RE_NT_pblib001_136 | B | >25 | >25 |
| V704_2448_240_RE_NT_pblib001_288 | B | >25 | >25 |
| V704_2544_140_RE_NT_pblib002_10 | B | >25 | >25 |
| V704_2555_240_RE_NT_pblib001_231 | B | >25 | >25 |
| V704_2684_181_RE_NT_pblib001_13 | B | >25 | >25 |
| V704_2684_181_RE_NT_pblib006_3 | B | >25 | >25 |
| V704_2767_070_RE_NT_pblib001_63 | B | >25 | >25 |
| V704_2834_210_RE_NT_pblib002_26 | B | >25 | >25 |
| V704_2834_210_RE_NT_pblib001_44 | B | >25 | >25 |
| V704_2839_140_RE_NT_pblib001_208 | B | >25 | >25 |
| V704_2981_150_RE_NT_pblib002_11 | B | >25 | >25 |
| V704_2981_150_RE_NT_pblib003_4 | B | >25 | >25 |
| V704_3008_040_RE_NT_pblib001_113 | B | >25 | >25 |
| V704_0372_250_RE_NT_pblib001_7 | B | >25 | >25 |

#### Table S5 B.

| Sequence Name HVTN 704 | HIV-1 M Env subtype | PGDM1400 | CAP256-VRC26.25 LS |
|---|---|---|---|
| V703_2117_110_RE_sga2A10_s | C | 0.0007 | 0.0001 |
| V703_2149_060_RE_sgaB10_s | C | 0.0012 | 0.0001 |
| V703_2018_240_RE_sga6A1_s | C | 0.0031 | 0.0001 |
| V703_1675_080_RE_con_s-modified | C | 0.0100 | 0.0001 |
| V703_1889_100_RE_con_s | C | 0.0100 | 0.0001 |
| V703_2631_150_RE_sga2F8_s | C | 0.0019 | 0.0004 |
| V703_2805_080_RE_con_s | C | 0.0019 | 0.0004 |
| V703_0739_110_RE_con_s-modified | C | 3.2300 | 0.0009 |
| V703_0217_050_RE_pblib002_s | C | 0.0028 | 0.0011 |
| V703_1313_040_RE_con_s | C | 0.0040 | 0.0018 |
| V703_0217_050_RE_sga2A3_s | C | 0.0040 | 0.0020 |
| V703_0013_090_RE_con_s | C | 0.0100 | 0.0100 |
| V703_0646_051_RE_con_s | C | 0.0100 | 0.0100 |
| V703_1675_080_RE_con_s | C | 0.0100 | 0.0100 |
| V703_1848_190_RE_pblib002_s | C | 0.0100 | 0.0100 |
| V703_2304_150_RE_con_s | C | 0.0100 | 0.0100 |
| V703_1471_190_RE_con_s | C | 0.0200 | 0.0100 |
| V703_0537_110_RE_sga4H1_s | C | 0.0900 | 0.0100 |
| V703_1298_080_RE_pblib002_s | C | 0.0900 | 0.0100 |
| V703_0472_030_RE_con_s | C | 0.1000 | 0.0100 |
| V703_0842_200_RE_con_s | C | 0.1100 | 0.0100 |
| V703_0537_110_RE_sga2B5_s | C | 0.2200 | 0.0100 |
| V703_0537_110_RE_pblib003_s | C | 1.0000 | 0.0100 |
| V703_2788_030_RE_con_s | C | 1.2000 | 0.0100 |
| V703_0629_150_RE_sga5A2_s | C | 0.4200 | 0.0500 |
| V703_0629_150_RE_pblib002_s | C | 0.3600 | 0.1100 |
| V703_1104_100_RE_sga10G5_s | C | 0.4700 | 0.1200 |
| V703_1104_100_RE_sga10A5_s | C | 0.9200 | 0.1200 |
| V703_1104_100_RE_pblib001_s | C | 0.3300 | 0.1400 |
| V703_0629_150_RE_pblib004_s | C | 0.1100 | 0.2200 |
| V703_1060_080_RE_con_s | C | >25 | 0.3100 |
| V703_1915_250_RE_sgaA3_s | C | >25 | 3.4800 |
| V703_2805_080_RE_pblib002_s | C | >25 | 5.9000 |
| V703_1750_140_RE_con_s | C | >25 | 8.5900 |
| V703_0926_070_RE_sga2H2_s | C | 1.0400 | >25 |
| V703_0629_150_RE_pblib003_s | C | 1.5100 | >25 |
| V703_1848_190_RE_sga6D1_s | C | 11.4600 | >25 |
| V703_0279_110_RE_con_s-modified | C | 21.6100 | >25 |
| V703_0203_081_RE_pblib002_s | C | >25 | >25 |
| V703_0203_081_RE_sga8A5_s | C | >25 | >25 |
| V703_0279_110_RE_pblib002_s | C | >25 | >25 |
| V703_0566_160_RE_con_s | C | >25 | >25 |
| V703_0566_160_RE_pblib002_s | C | >25 | >25 |
| V703_1764_250_RE_con_s | C | >25 | >25 |
| V703_1789_230_RE_sga3A3_s | C | >25 | >25 |
| V703_2539_070_RE_sga6F6_s | C | >25 | >25 |

#### Table S5 C

Table S5 C.

| Sequence Name HVTN 704 | HIV-1 M Env subtype | PGDM1400 | CAP256-VRC26.25LS |
|---|---|---|---|
| V704_0575_060_RE_NT_pblib002_16 | Atypical B | 0.051 | >25 |
| V704_0575_060_RE_NT_pblib001_60 | Atypical B | 0.047 | >25 |
| V704_0575_060_RE_NT_con02_s | Atypical B | 0.042 | >25 |
| V704_0644_060_RE_NT_pblib001_216 | Atypical B | >25 | >25 |
| V704_1183_220_RE_NT_pblib001_70 | B/F | >25 | >25 |
| V704_2788_060_RE_NT_pblib001_96 | B/F | 0.003 | 0.0003 |
| V704_2788_060_RE_NT_pblib002_20 | B/F | 0.011 | 0.011 |
| V704_1930_170_RE_NT_pblib002_114 | 12_BF/F1 | 0.014 | >25 |
| V704_1930_170_RE_NT_pblib001_234 | 12_BF/F1 | 0.018 | >25 |
| V704_1180_070_RE_NT_pblib001_87 | A1/B | 0.188 | >25 |
| V704_1180_070_RE_NT_pblib027_1 | A1/B | 0.405 | >25 |
| V704_0496_040_RE_NT_pblib001_37 | F | 0.011 | 0.011 |
| V704_1775_030_RE_NT_pblib001_135 | F1 | 0.002 | 0.028 |
| V704_0128_220_RE_NT_pblib001_52 | F1/B | 0.008 | 0.002 |
| V704_0128_220_RE_NT_pblib002_40 | F1/B | 0.031 | 0.0003 |
| V704_1109_140_RE_NT_pblib001_191 | F1/B | 0.201 | >25 |
| V704_1429_090_RE_NT_pblib002_75 | F2 | 0.079 | 11.679 |
| V704_1429_090_RE_NT_pblib001_111 | F2 | 0.141 | >25 |
| V703_0865_070_RE_con_s | G | ND | ND |

**Table S5. Clade specificity of V2 apex antibodies. A.** IC50s of V2 apex antibodies against B clade AMP viruses. **B.** C clade AMP viruses. **C.** Other subtypes and recombinant viruses found among the AMP placebo HIV-1 infections. The V2 apex bNab CAP256 VRC26.25 LS has substantial breadth and great potency among C clade viruses but has almost no neutralizing activity against B clade viruses. PGDM1400 shares this pattern but is less extreme, with a few modest B clade responses and less potent but still strong neutralization of C clade viruses. F clade viruses (in Table C) can be sensitivity to both antibodies and are generally more like C clade viruses than B clade. B/F recombinants, indicated by the forward slash, were each checked for recombination breakpoints, and all were found to be F-like in the relevant V2 apex epitope region.

### Table S6

| | CD4bs | | | | | | V3 | | V2 | | FP | | | MPER | | |
|---|---|---|---|---|---|---|---|---|---|---|---|---|---|---|---|---|
| Sequence Name | 1-18 | VRC07- | VRC01-23 | 3BNC117- | VRC01 | CH235.12 | PGT121- | 10-1074 | PGDM140 | CAP256 | PGT151 | VRC34.01 | ACS202 | 10E8v4 | CAP248- | Subtype |
| V703_2805_080_RE_pblib002_s | >10 | 1.87 | >10 | 5.17 | >25 | >10 | >25 | >25 | >25 | 5.9 | >10 | 0.84 | >10 | 2.38 | >10 | C |
| V704_1109_140_RE_NT_pblib001_191 | >25 | 0.283 | 2.455 | >25 | 3.360 | >25 | >25 | >25 | 0.201 | >25 | >25 | >25 | >25 | 1.25 | >25 | F1B |
| V703_0279_110_RE_pblib002_s | 0.17 | 0.18 | 0.14 | 0.33 | 1.509 | >10 | 0.94 | 1.19 | >25 | >25 | >10 | >10 | >10 | 0.34 | >10 | C |
| V704_0513_150_RE_NT_pblib001_34 | 0.612 | 0.38 | 0.329 | 0.276 | 1.490 | 6.07 | >25 | >25 | >25 | >25 | 0.49 | >25 | >25 | 1.13 | >25 | B |
| V704_0575_060_RE_NT_con02_s | 0.467 | 0.756 | 0.771 | 0.965 | 2.118 | 2.354 | >25 | >25 | 0.042 | >25 | >25 | >25 | 0.157 | 0.494 | >25 | B |
| V704_0644_060_RE_NT_pblib001_216 | 0.084 | 0.155 | 0.279 | 0.29 | 1.020 | 3.275 | 6.348 | 0.454 | >25 | >25 | >25 | 5.728 | >25 | 0.297 | >25 | B |
| V704_0726_080_RE_NT_pblib001_26 | 0.199 | 1.02 | 0.466 | 0.537 | 5.636 | 3.445 | 0.36 | 0.734 | 0.421 | >25 | >25 | >25 | >25 | 2.473 | >25 | B |
| V704_1429_090_RE_NT_pblib001_111 | 0.119 | 0.112 | 0.122 | 1.806 | 0.307 | >25 | >25 | >25 | 0.141 | >25 | 4.52 | >25 | >25 | 0.45 | >25 | F2 |
| V704_1835_150_RE_NT_pblib002_9 | 0.026 | 24.845 | 0.078 | >25 | >100 | >25 | 3.791 | 0.081 | >25 | >25 | 0.24 | 1.729 | 0.397 | 0.099 | >25 | B |
| V704_1044_140_RE_NT_pblib001_11 | 0.99 | 0.431 | 0.494 | 0.82 | 5.76 | 8.648 | >25 | 0.421 | >25 | >25 | 0.059 | >25 | >25 | 1.127 | 10 | B |
| V704_2834_210_RE_NT_pblib001_44 | 0.107 | 0.312 | 0.073 | 0.086 | 2.190 | >25 | 19.219 | 4.297 | >25 | >25 | >25 | 0.922 | >25 | 0.12 | >25 | B |
| V704_2839_140_RE_NT_pblib001_208 | 0.116 | 0.183 | 0.346 | 1.426 | 1.580 | >25 | >25 | >25 | >25 | >25 | >25 | 1.771 | 1.486 | 0.63 | >25 | B |
| V704_2981_150_RE_NT_pblib003_4 | 1.058 | 0.39 | 0.178 | 0.155 | 1.840 | 0.333 | >25 | >25 | >25 | >25 | >25 | >25 | >25 | 0.28 | >25 | B |
| V704_3000_240_RE_NT_pblib019_1 | 0.062 | 0.201 | 0.562 | 0.271 | 1.920 | >25 | >25 | >25 | 16.221 | >25 | >25 | 0.145 | >25 | 0.14 | >25 | B |
| V703_1789_230_RE_sga3A3_s | 0.09 | 0.99 | 0.55 | >25 | 17.896 | >10 | 0.83 | 0.34 | >25 | >25 | >10 | >10 | >10 | 2.43 | >10 | C |
| V703_1675_080_RE_con_s | 0.0279 | 0.05 | 0.0528 | 0.04 | 0.378 | 0.23 | 0.26 | 0.79 | 0.01 | 0.01 | >10 | >10 | 1.49 | 14.66 | >10 | C |
| V704_0128_220_RE_NT_pblib002_40 | 0.044 | 0.026 | 0.034 | 0.025 | 0.160 | 0.089 | 0.047 | 0.171 | 0.031 | 0.0003 | >25 | 2.044 | >25 | 1.33 | >25 | F1B |
| V704_0496_040_RE_NT_pblib001_37 | 0.037 | 0.242 | 0.601 | >25 | 4.940 | 1.236 | 0.142 | 0.078 | 0.011 | 0.011 | >25 | >25 | >25 | 0.289 | >25 | B |
| V704_1775_030_RE_NT_pblib001_135 | 0.014 | 0.029 | 0.031 | 0.109 | 0.090 | 0.05 | >25 | >25 | 0.002 | 0.028 | 0.112 | >25 | >25 | 0.291 | >25 | F1 |
| V704_2788_060_RE_NT_pblib002_20 | 0.192 | 0.26 | 0.284 | 0.223 | 1.409 | 1.144 | 0.134 | 0.144 | 0.011 | 0.011 | >10 | >10 | >10 | 0.082 | >10 | BF |
| V703_0472_030_RE_con_s | 0.0054 | 0.01 | 0.005 | >25 | 0.021 | 0.05 | >25 | 7.94 | 0.1 | 0.01 | 0.028 | 5.67 | >10 | 0.42 | >10 | C |
| V703_0537_110_RE_pblib003_s | 0.0138 | 0.02 | 0.005 | 0.03 | 0.097 | 0.19 | 0.05 | 0.21 | 1 | 0.01 | >10 | >10 | >10 | 0.26 | >10 | C |
| V703_0739_110_RE_con_s-modified | 0.025 | 0.05 | 0.023 | 0.07 | 0.143 | 0.56 | 0.67 | 0.16 | 3.23 | 0.0009 | >10 | 4.86 | 0.09 | 1.94 | >10 | C |
| V703_1313_040_RE_con_s | 0.046 | 0.04 | 0.07 | 0.11 | 0.686 | >10 | 0.01 | 0.06 | 0.004 | 0.0018 | >10 | >10 | >10 | 0.01 | >10 | C |
| V703_2304_150_RE_con_s | 0.06 | 0.02 | 0.0106 | 0.06 | 0.114 | 0.08 | 0.07 | 0.06 | 0.01 | 0.01 | >10 | 0.03 | >10 | 0.36 | >10 | C |
| V703_0842_200_RE_con_s | >10 | 1.44 | 1.82 | 2.13 | 20.895 | 6.36 | 7.23 | 1.56 | 0.11 | 0.01 | 0.21 | >10 | >10 | 0.73 | >10 | C |
| V703_0013_090_RE_con_s | 0.52 | 0.42 | 0.49 | >25 | 3.472 | >10 | 7.97 | 18.41 | 0.01 | 0.01 | >10 | >10 | >10 | 1.61 | >10 | C |
| V703_0646_051_RE_con_s | 0.18 | 0.45 | 0.63 | 1.54 | 4.897 | >10 | >25 | >25 | 0.01 | 0.01 | 0.07 | 0.07 | >10 | 1.56 | >10 | C |
| V703_1104_100_RE_sga10G5_s | 0.28 | 1.61 | 1.41 | 2.99 | 6.410 | >10 | >25 | 10.48 | 0.47 | 0.12 | >10 | >10 | 4.23 | 0.86 | >10 | C |
| V703_1298_080_RE_pblib002_s | 0.15 | 1.27 | 0.13 | >25 | >25 | 0.16 | >25 | >25 | 0.09 | 0.01 | 0.1 | >10 | >10 | 0.84 | >10 | C |
| V703_2788_030_RE_con_s | 1.56 | 0.24 | 4.11 | 0.09 | 0.6812 | 4.9 | 0.3 | 0.09 | 1.2 | 0.01 | 1.84 | >10 | >10 | 5.38 | >10 | C |
| V704_1783_150_RE_NT_pblib001_144 | 0.006 | 0.064 | 0.009 | 0.036 | 0.135 | >25 | 0.248 | 0.485 | >25 | >25 | 0.06 | >25 | 0.353 | 0.388 | >25 | B |
| V704_0855_080_RE_NT_pblib001_187 | 0.012 | 0.003 | 0.002 | 0.009 | 0.013 | >25 | 0.05 | 0.061 | >25 | >25 | 0.007 | >25 | >25 | 0.134 | >25 | B |
| V704_1180_070_RE_NT_pblib027_1 | 0.005 | 0.074 | 0.009 | 0.018 | 0.137 | 0.419 | >25 | >25 | 0.405 | >25 | 0.009 | 0.683 | 0.388 | 0.16 | >25 | A1B |
| V704_1535_030_RE_NT_pblib001_471 | 0.052 | 0.068 | 0.018 | 0.019 | 0.088 | >25 | 0.011 | 0.071 | 1.109 | >25 | 0.197 | 0.391 | 0.512 | 0.015 | 0.334 | B |
| V704_1041_080_RE_NT_pblib042_1 | 0.026 | 0.036 | 0.018 | 0.021 | 0.21 | 0.112 | 0.022 | 0.027 | 0.218 | >25 | 0.168 | 2.872 | >25 | 0.08 | >10 | B |
| V704_2767_070_RE_NT_pblib001_63 | 0.217 | 0.126 | 0.068 | 0.796 | 2.060 | 1.275 | 0.066 | 0.241 | >25 | >25 | 0.038 | >25 | >25 | 2.03 | >25 | B |
| V704_0445_180_RE_NT_pblib001_22 | 0.012 | 0.063 | 0.021 | >25 | >25 | 0.205 | 0.291 | 0.086 | 0.285 | >25 | 0.083 | >25 | 0.08 | 1.49 | >25 | B |
| V703_0566_160_RE_pblib002_s | 0.06 | 0.18 | 0.17 | 0.66 | 3.503 | 1.28 | 0.03 | 0.06 | >25 | >25 | 0.2 | 0.08 | >10 | 4.21 | >10 | C |
| V703_1750_140_RE_con_s | 0.11 | 0.18 | 0.13 | 0.23 | 0.881 | 1.35 | 0.05 | 0.08 | >25 | 8.59 | 0.05 | 0.34 | >10 | 0.5 | >10 | C |
| V703_1915_250_RE_sgaA3_s | 0.0841 | 0.36 | 0.0406 | 1.6 | 0.529 | 5.6 | 0.32 | 0.84 | >25 | 3.48 | 0.04 | >10 | >10 | 3.83 | >10 | C |
| V703_0217_050_RE_sga2A3_s | 0.008 | 0.03 | 0.04 | >25 | >25 | 0.29 | >25 | >25 | 0.004 | 0.002 | 0.01 | 0.0253 | 2.21 | 0.18 | 0.0247 | C |
| V704_0746_760_RE_NT_pblib002_36 | 0.224 | 0.447 | 0.189 | 0.282 | 1.250 | 6.796 | 0.02 | 0.015 | >25 | >25 | >25 | >25 | >25 | 1.96 | >25 | B |
| V704_0026_231_RE_NT_pblib001_3 | 0.097 | 0.081 | 0.088 | 0.103 | 0.460 | >25 | 0.006 | 0.017 | 0.075 | >25 | >25 | >25 | >25 | 0.07 | >25 | B |
| V704_0847_030_RE_NT_pblib002_23 | 9.694 | 0.204 | 0.161 | >25 | 3.118 | >25 | 0.011 | 0.006 | >25 | >25 | >25 | >25 | >25 | 0.27 | 1.448 | B |
| V704_0856_240_RE_NT_pblib001_9 | 0.095 | 0.063 | 0.055 | 0.06 | 0.524 | 0.503 | 0.011 | 0.016 | >25 | >25 | 1.04 | 7.845 | >25 | 0.54 | >25 | B |
| V704_0886_250_RE_NT_pblib001_10 | 0.173 | 0.177 | 0.099 | 0.516 | 1.745 | >25 | 0.338 | 0.185 | >25 | >25 | >25 | >25 | 0.498 | 0.171 | >25 | B |
| V704_0911_150_RE_NT_pblib001_58 | 0.437 | 0.538 | 1.525 | 0.622 | 1.391 | 10.111 | 0.013 | 0.018 | >25 | >25 | 3.99 | >25 | >25 | 0.869 | >25 | B |
| V704_1183_220_RE_NT_pblib001_70 | 0.041 | 0.347 | 0.038 | 0.065 | 0.390 | 1.185 | 0.032 | 0.051 | >25 | >25 | 3.666 | >25 | 0.228 | 0.495 | >25 | BF |
| V704_1055_240_RE_NT_pblib001_231 | 0.092 | 0.052 | 0.033 | 0.075 | 0.424 | 10 | 0.012 | 0.03 | >25 | >25 | >25 | 2.808 | 0.151 | >25 | >25 | B |
| V704_2684_181_RE_NT_pblib006_3 | 0.007 | 0.057 | 0.012 | 0.43 | 0.092 | >25 | 0.007 | 0.019 | >25 | >25 | >25 | >25 | 0.97 | 0.031 | >25 | B |
| V704_3008_040_RE_NT_pblib001_113 | 0.067 | 0.431 | 0.025 | 0.163 | 3.020 | >25 | 0.017 | 0.178 | >25 | >25 | >25 | >25 | >25 | 0.184 | >25 | B |
| V703_0203_081_RE_sga8A5_s | 0.64 | 0.41 | 0.91 | >10 | 4.151 | >10 | 0.03 | 0.02 | >10 | >10 | >10 | >10 | >10 | 0.13 | >10 | C |
| V703_0926_070_RE_sga2H2_s | 2.02 | 0.81 | 0.76 | 2.04 | 7.803 | 2.34 | 0.04 | 0.1 | 1.04 | >25 | >10 | >10 | >10 | 0.56 | >10 | C |
| V703_1039_070_RE_sga6F6_s | 0.74 | 0.39 | 0.27 | >10 | 1.7655 | >10 | 0.01 | 0.04 | >10 | >10 | 4.57 | >10 | >10 | 0.23 | >10 | C |
| V703_1471_190_RE_con_s | 0.05 | 0.08 | 0.0235 | 0.06 | 0.161 | 0.6 | 0.01 | 0.1 | 0.02 | 0.01 | 0.01 | >10 | >10 | 4.92 | 0.011 | C |
| V704_2095_130_RE_NT_pblib001_136 | 0.079 | 0.096 | 0.031 | 0.053 | 0.312 | 0.283 | 2.167 | 0.085 | >25 | >25 | 0.147 | >25 | >25 | 0.51 | >25 | B |
| V704_0944_180_RE_NT_pblib001_76 | 0.023 | 0.02 | 0.007 | 0.061 | 0.074 | >25 | >25 | >25 | >25 | >25 | 0.367 | >25 | >25 | >25 | >25 | B |
| V704_1706_040_RE_NT_pblib001_26 | 0.2 | 0.058 | 0.058 | 0.164 | 0.489 | 0.186 | 0.457 | 0.301 | 1.986 | >25 | >25 | >25 | >25 | 0.488 | >25 | B |
| V704_1930_170_RE_NT_pblib001_234 | 0.041 | 0.035 | 0.023 | 0.043 | 0.134 | 0.118 | >25 | >25 | 0.018 | >25 | >25 | >25 | >25 | 1.25 | >25 | 12.BF |
| V704_2065_060_RE_NT_pblib002_42 | 0.062 | 0.082 | 0.037 | 0.048 | 0.220 | 0.036 | >25 | >25 | >25 | >25 | >25 | >25 | >25 | 0.161 | >25 | B |
| V704_2448_240_RE_NT_pblib001_288 | 0.133 | 0.108 | 0.01 | 0.058 | 0.140 | 0.146 | >25 | >25 | >25 | >25 | >25 | >25 | 7.416 | >25 | >25 | B |
| V703_0629_150_RE_pblib003_s | 0.11 | 0.33 | 0.06 | 0.72 | 1.076 | 1.5 | 9.87 | 5.61 | 1.51 | >25 | 0.06 | >10 | >10 | 6.44 | >10 | C |
| V703_1060_080_RE_con_s | 0.09 | 0.13 | 0.28 | 0.8 | 1.712 | 1.17 | >25 | 3.63 | >25 | 0.31 | 0.05 | >10 | >10 | 0.08 | >10 | C |
| V703_1764_250_RE_con_s | 0.03 | 0.1 | 0.06 | 0.1 | 0.240 | 0.1 | 0.2 | 0.13 | >25 | >25 | 0.18 | >10 | >10 | 1.41 | >10 | C |
| V703_1848_190_RE_sga6D1_s | 0.09 | 0.31 | 0.2 | 0.24 | 1.054 | 0.24 | >25 | >25 | 11.46 | >25 | 0.02 | >10 | >10 | 2.76 | >10 | C |
| V703_2018_240_RE_sga6A1_s | 0.16 | 0.3 | 0.09 | 1.13 | 1.249 | 6.38 | 0.05 | 0.07 | 0.0031 | 0.00013 | >10 | >10 | >10 | 2.85 | >10 | C |
| V704_1991_230_RE_NT_pblib002_19 | 0.067 | 0.215 | 0.14 | 0.165 | 1.488 | 0.309 | >25 | >25 | 0.052 | 0.0003 | >25 | >25 | 0.12 | 0.317 | >25 | B |
| V703_1889_100_RE_con_s | 0.09 | 2.3 | 0.19 | >25 | >25 | 5.78 | 0.09 | 0.23 | 0.01 | 0.00013 | >10 | >10 | >10 | >25 | >10 | C |
| V703_2117_110_RE_sga2A10_s | 0.17 | 0.04 | 0.06 | 0.11 | 0.228 | 0.52 | >25 | >25 | 0.0007 | 0.00013 | >10 | 0.81 | >10 | 0.88 | >10 | C |
| V703_2149_060_RE_sgaB10_s | 0.0324 | 0.07 | 0.0208 | >25 | 0.205 | >10 | 0.11 | 0.07 | 0.0012 | 0.00013 | 9.17 | >10 | >10 | 0.98 | >10 | C |
| V703_2631_150_RE_sga2F8_s | 0.02 | 0.08 | 0.09 | 0.07 | 0.463 | 1.02 | >25 | >25 | 0.0019 | 0.0004 | >10 | 0.97 | >10 | 1.23 | >10 | C |
| V704_0907_130_RE_NT_pblib001_32 | 0.016 | 0.193 | 0.019 | 0.199 | 0.364 | 15.857 | 0.002 | 0.006 | >25 | >25 | 0.017 | 0.25 | 0.236 | 0.655 | >25 | B |
| V704_1528_240_RE_NT_pblib001_17 | 0.06 | 0.439 | 0.114 | 0.092 | 2.390 | 7.107 | 0.003 | 0.004 | >25 | >25 | 1.418 | 0.04 | 0.041 | 0.243 | 1.15 | B |

**Table S6. K-means clusters used to define the REP-T2 panel, the centroids of each cluster were used as panel members and are marked in pink.**

### Table S7

SHEP-T2 panel phenotyping

| | Tier 2 | Tier 2 | Tier 2 | Tier 2 | Tier 2 | Tier 2 | Tier 2 | Tier 2 | Tier 2 | Tier 2 | Tier 2 | Tier 2 |
|---|---|---|---|---|---|---|---|---|---|---|---|---|
| | H703_1313_040s | H704_1535_030sN | H704_0855_080_EsN | H704_0907_130sN | H704_1528_240_RE_pblib_001_s | H704_1180_070EsN | H704_0445_180_RE_con_s | H703_2304_150_RE_cs | H703_2805_080Es | H703_2117_110_RE_e2A10s | H704_0128_220_RE_pb001_s | H703_0472_030s |
| Specimen ID | Clade C | Clade B | Clade B | Clade B | Clade B | Clade A1/B | Clade B | Clade C | Clade C | Clade C | Clade F1/B | Clade C |
| SA-C10 | 1623 | 468 | 789 | 795 | 505 | 588 | >1666.67 | >1666.67 | >1666.67 | >1666.67 | >1666.67 | >1666.67 |
| SA-C48 | 421 | 136 | 919 | 1088 | 652 | 587 | >1666.67 | >1666.67 | >1666.67 | >1666.67 | >1666.67 | >1666.67 |
| SA-C72 | 461 | 168 | >1666.67 | >1666.67 | 186 | 821 | 753 | >1666.67 | 629 | 1381 | >1666.67 | 534 |
| SA-C74 | >1666.67 | 84 | >1666.67 | 367 | 152 | 783 | >1666.67 | 1403 | 1028 | 489 | >1666.67 | >1666.67 |
| SA-C90 | 1653 | 782 | 1150 | 375 | 145 | >1666.67 | 738 | 1192 | 822 | 1181 | 1508 | >1666.67 |
| 2219 | >16.667 | >16.667 | >16.667 | >16.667 | >16.667 | >16.667 | >16.667 | >16.667 | >16.667 | >16.667 | >16.667 | >16.667 |
| 2557 | >16.667 | >16.667 | >16.667 | >16.667 | >16.667 | >16.667 | >16.667 | >16.667 | >16.667 | >16.667 | >16.667 | >16.667 |
| 3074 | >16.667 | 13 | >16.667 | >16.667 | >16.667 | >16.667 | >16.667 | >16.667 | >16.667 | >16.667 | >16.667 | >16.667 |
| 3869 | >16.667 | >16.667 | >16.667 | >16.667 | >16.667 | >16.667 | >16.667 | >16.667 | >16.667 | >16.667 | >16.667 | >16.667 |
| 447-52D | >16.667 | >16.667 | >16.667 | >16.667 | >16.667 | >16.667 | >16.667 | >16.667 | >16.667 | >16.667 | >16.667 | >16.667 |
| 838-12D | >16.667 | >16.667 | >16.667 | >16.667 | >16.667 | >16.667 | >16.667 | >16.667 | >16.667 | >16.667 | >16.667 | >16.667 |
| 654-30D | >16.667 | >16.667 | >16.667 | >16.667 | >16.667 | >16.667 | >16.667 | >16.667 | >16.667 | >16.667 | >16.667 | >16.667 |
| 1008-30D | >16.667 | >16.667 | >16.667 | >16.667 | >16.667 | >16.667 | >16.667 | >16.667 | >16.667 | >16.667 | >16.667 | >16.667 |
| 729-30D | >16.667 | >16.667 | >16.667 | >16.667 | >16.667 | >16.667 | >16.667 | >16.667 | >16.667 | >16.667 | >16.667 | >16.667 |
| F105 | >16.667 | >16.667 | >16.667 | >16.667 | >16.667 | >16.667 | >16.667 | >16.667 | >16.667 | >16.667 | >16.667 | >16.667 |
| VRC01 | 1.25 | 0.21 | 0.09 | 0.96 | 3.31 | 0.28 | >6.667 | 1.89 | >6.667 | 0.3 | 0.2 | 0.308 |

REP-T2 panel phenotyping

| | Tier 2 | Tier 2 | Tier 2 | Tier 2 | Tier 2 | Tier 2 | Tier 2 | Tier 2 | Tier 2 | Tier 2 | Tier 2 | Tier 2 |
|---|---|---|---|---|---|---|---|---|---|---|---|---|
| | H703_0217_050e_2A3 | H703_0842_200Es | H703_1471_190s | H703_1675_080s | H703_2805_080_RE_pblib002_s | H703_2018_240_RE_e6A1s | H704_0746_760_RE_p001s | H704_2095_130_RE_cs | H704_2767_070sN | H704_0907_130sN | H704_1783_150_RE_cs | V703_0279_110_RE_pblib002_s |
| | Clade C | Clade C | Clade C | Clade C | Clade C | Clade C | Clade B | Clade B | Clade B | Clade B | Clade B | Clade B |
| Specimen ID | IC50 | IC50 | IC50 | IC50 | IC50 | IC50 | IC50 | IC50 | IC50 | IC50 | IC50 | IC50 |
| SA-C10 | 639.317 | >2500.0 | 1160.687 | 210.727 | >2500.0 | >1666.67 | 2286.262 | >2500.0 | >2500.0 | 795 | >1666.67 | 315.597 |
| SA-C48 | 277.999 | >2500.0 | 399.278 | > 2500 | >2500.0 | 1017.621 | 2371.204 | 1385.695 | >2500.0 | 1088 | >1666.67 | 444.906 |
| SA-C72 | 113.266 | 1477.2 | 291.491 | 1025.910 | >2500.0 | >1666.67 | >2500.0 | 1406.991 | >2500.0 | >1666.67 | >1666.67 | 313.508 |
| SA-C74 | 711.452 | >2500.0 | 671.339 | 1185.753 | >2500.0 | 1256.247 | >2500.0 | >2500.0 | >2500.0 | 367 | >1666.67 | 1715.938 |
| SA-C90 | 260.549 | 1609.68 | 575.145 | > 2500 | >2500.0 | >1666.67 | >2500.0 | >2500.0 | >2500.0 | 375 | 1277 | 740.313 |
| 2219 | >25.0 | >25.0 | >25.0 | >25.0 | >25.0 | >16.67 | >25.0 | 5.502 | >25.0 | >16.667 | >16.67 | >25.0 |
| 2557 | >25.0 | >25.0 | >25.0 | >25.0 | >25.0 | >16.67 | >25.0 | 16.476 | >25.0 | >16.667 | >16.67 | >25.0 |
| 3074 | 21.435 | >25.0 | >25.0 | >25.0 | >25.0 | >16.67 | >25.0 | >25.0 | >25.0 | >16.667 | >16.67 | >25.0 |
| 3869 | >25.0 | >25.0 | >25.0 | >25.0 | >25.0 | >16.67 | >25.0 | >25.0 | >25.0 | >16.667 | >16.67 | >25.0 |
| 447-52D | >25.0 | >25.0 | >25.0 | >25.0 | >25.0 | >16.67 | >25.0 | >25.0 | >25.0 | >16.667 | >16.67 | >25.0 |
| 838-12D | >25.0 | >25.0 | >25.0 | >25.0 | >25.0 | >16.67 | >25.0 | >25.0 | >25.0 | >16.667 | >16.67 | >25.0 |
| 654-30D | >25.0 | >25.0 | >25.0 | >25.0 | >25.0 | >16.67 | >25.0 | >25.0 | >25.0 | >16.667 | >16.67 | >25.0 |
| 1008-30D | >25.0 | >25.0 | >25.0 | >25.0 | >25.0 | >16.67 | >25.0 | >25.0 | >25.0 | >16.667 | >16.67 | >25.0 |
| 729-30D | >25.0 | >25.0 | >25.0 | >25.0 | >25.0 | >16.67 | >25.0 | >25.0 | >25.0 | >16.667 | >16.67 | >25.0 |
| F105 | >25.0 | >25.0 | >25.0 | >25.0 | >25.0 | >16.67 | >25.0 | >25.0 | >25.0 | >16.667 | >16.67 | >25.0 |
| VRC01 | >25.0 | >25.0 | 0.392 | 1.037 | >25.0 | 2.870 | 1.418 | 0.624 | >25.0 | 0.96 | 0.22 | 1.484 |

### Table S7, continued

**MPER Sensitive**

| | Tier 2 | Tier 2 | Tier 2 | Tier 2 | Tier 2* | Tier 2 | Tier 2 | Tier 2 | Tier 2** |
|---|---|---|---|---|---|---|---|---|---|
| | H703_1313<br>_040s | H704_1535<br>_030sN | H704_2684<br>_181_RE_p<br>001s | H704_0026<br>_231_RE_p<br>bsga001_s | H703_1060<br>_080s | H703_0279<br>_110s | V704_0372<br>_250_RE_p<br>blib001_s | H704_0855<br>_080_EsN | H704_1835<br>_150_RE_p0<br>01s_2484A |
| Specimen ID | Clade C | Clade B | Clade B | Clade B | Clade C | Clade C | Clade B | Clade B | Clade B |
| SA-C10 | 1623 | 468 | 90 | 1116 | 1182.447 | 528.4 | 388 | 789 | 874 |
| SA-C48 | 421 | 136 | 511 | 1075 | 1455.479 | 410.213 | 623 | 919 | 421 |
| SA-C72 | 461 | 168 | 301 | >1666.67 | 1429.193 | 400.006 | >1666.67 | >1666.67 | 177 |
| SA-C74 | >1666.67 | 84 | 215 | >1666.67 | 716.276 | 1646.61 | >1666.67 | >1666.67 | 265 |
| SA-C90 | 1653 | 782 | 1243 | 644 | 766.31 | 1298.502 | >1666.67 | 1150 | 499 |
| 2219 | >16.667 | >16.667 | >16.667 | >16.667 | >16.67 | >16.67 | >16.67 | >16.667 | 0.9 |
| 2557 | >16.667 | >16.667 | >16.667 | >16.667 | >16.67 | >16.67 | >16.67 | >16.667 | 1.98 |
| 3074 | >16.667 | 13 | >16.667 | >16.667 | >16.67 | >16.67 | >16.67 | >16.667 | >16.67 |
| 3869 | >16.667 | >16.667 | >16.667 | >16.667 | >16.67 | >16.67 | >16.67 | >16.667 | 14.28 |
| 447-52D | >16.667 | >16.667 | >16.667 | >16.667 | >16.67 | >16.67 | >16.67 | >16.667 | >16.67 |
| 838-12D | >16.667 | >16.667 | >16.667 | >16.667 | >16.67 | >16.67 | >16.67 | >16.667 | 5.82 |
| 654-30D | >16.667 | >16.667 | >16.667 | >16.667 | >16.67 | >16.67 | >16.67 | >16.667 | >16.67 |
| 1008-30D | >16.667 | >16.667 | >16.667 | >16.667 | >16.67 | >16.67 | >16.67 | >16.667 | >16.67 |
| 729-30D | >16.667 | >16.667 | >16.667 | >16.667 | >16.67 | >16.67 | >16.67 | >16.667 | >16.67 |
| F105 | >16.667 | >16.667 | >16.667 | >16.667 | >16.67 | >16.67 | >16.67 | >16.667 | >16.67 |
| VRC01 | 1.25 | 0.21 | 0.24 | 1.16 | 2.856 | 1.537 | 0.64 | 0.09 | >6.67 |

**V3 glycan sensitive**

| | Tier 2 | Tier 2 | Tier 2 | Tier 2 | Tier 2 | Tier 2 | Tier 2 | Tier 2 |
|---|---|---|---|---|---|---|---|---|
| | H704_0907<br>_130sN | H704_1528<br>_240_RE_p | H704_0847<br>_030_EsN | H704_0026<br>_231_RE_p | H704_2684<br>_181_RE_p | H703_0566<br>_160s | H704_2541<br>_080_RE_p | H703_1313<br>_040s |
| Specimen ID | Clade B | Clade B | Clade B | Clade B | Clade B | Clade C | Clade B | Clade C |
| SA-C10 | 795 | 505 | 226 | 1116 | 90 | 580.509 | 729 | 1623 |
| SA-C48 | 1088 | 652 | 770 | 1075 | 511 | 1527.773 | 364 | 421 |
| SA-C72 | >1666.67 | 186 | >1666.67 | >1666.67 | 301 | 380.616 | 589 | 461 |
| SA-C74 | 367 | 152 | >1666.67 | >1666.67 | 215 | 741.785 | 552 | >1666.67 |
| SA-C90 | 375 | 145 | 239 | 644 | 1243 | 51.964 | 420 | 1653 |
| 2219 | >16.667 | >16.667 | >16.667 | >16.667 | >16.667 | >16.67 | >16.667 | >16.667 |
| 2557 | >16.667 | >16.667 | >16.667 | >16.667 | >16.667 | >16.67 | >16.667 | >16.667 |
| 3074 | >16.667 | >16.667 | >16.667 | >16.667 | >16.667 | >16.67 | >16.667 | >16.667 |
| 3869 | >16.667 | >16.667 | >16.667 | >16.667 | >16.667 | >16.67 | >16.667 | >16.667 |
| 447-52D | >16.667 | >16.667 | >16.667 | >16.667 | >16.667 | >16.67 | >16.667 | >16.667 |
| 838-12D | >16.667 | >16.667 | >16.667 | >16.667 | >16.667 | >16.67 | >16.667 | >16.667 |
| 654-30D | >16.667 | >16.667 | >16.667 | >16.667 | >16.667 | >16.67 | >16.667 | >16.667 |
| 1008-30D | >16.667 | >16.667 | >16.667 | >16.667 | >16.667 | >16.67 | >16.667 | >16.667 |
| 729-30D | >16.667 | >16.667 | >16.667 | >16.667 | >16.667 | >16.67 | >16.667 | >16.667 |
| F105 | >16.667 | >16.667 | >16.667 | >16.667 | >16.667 | >16.67 | >16.667 | >16.667 |
| VRC01 | 0.96 | 3.31 | >6.667 | 1.16 | 0.24 | >6.67 | 1.02 | 1.25 |

**FP sensitive**

| | Tier 2 | Tier 2 | Tier 2 | Tier 2 | Tier 2 | Tier 2 | Tier 2 | Tier 3 |
|---|---|---|---|---|---|---|---|---|
| | H704_1528<br>_240_RE_p<br>blib_001_s | H703_0629<br>_150_RE_e<br>5A2s | H703_0566<br>_160s | H704_1180<br>_070EsN | H704_0445<br>_180_RE_co<br>n_s | H704_0575<br>_060_RE_p<br>002s | H703_2304<br>_150_RE_cs | H703_0646<br>_051sN |
| Specimen ID | Clade B | Clade C | Clade C | Clade A1/B | Clade B | Clade B | Clade C | Clade C |
| SA-C10 | 505 | >1666.67 | 580.509 | 588 | >1666.67 | 928 | >1666.67 | >1666.67 |
| SA-C48 | 652 | >1666.67 | 1527.773 | 587 | >1666.67 | 781 | >1666.67 | >1666.67 |
| SA-C72 | 186 | 1073.293 | 380.616 | 821 | 753 | 1569 | >1666.67 | >1666.67 |
| SA-C74 | 152 | >1666.67 | 741.785 | 783 | >1666.67 | 501 | 1403 | >1666.67 |
| SA-C90 | 145 | 139.3 | 51.964 | >1666.67 | 738 | 1336 | 1192 | >1666.67 |
| 2219 | >16.667 | >16.67 | >16.67 | >16.667 | >16.667 | >16.67 | >16.667 | >16.67 |
| 2557 | >16.667 | >16.67 | >16.67 | >16.667 | >16.667 | >16.67 | >16.667 | >16.67 |
| 3074 | >16.667 | >16.67 | >16.67 | >16.667 | >16.667 | >16.67 | >16.667 | >16.67 |
| 3869 | >16.667 | >16.67 | >16.67 | >16.667 | >16.667 | >16.67 | >16.667 | >16.67 |
| 447-52D | >16.667 | >16.67 | >16.67 | >16.667 | >16.667 | >16.67 | >16.667 | >16.67 |
| 838-12D | >16.667 | >16.67 | >16.67 | >16.667 | >16.667 | >16.67 | >16.667 | >16.67 |
| 654-30D | >16.667 | >16.67 | >16.67 | >16.667 | >16.667 | >16.67 | >16.667 | >16.67 |
| 1008-30D | >16.667 | >16.67 | >16.67 | >16.667 | >16.667 | >16.67 | >16.667 | >16.67 |
| 729-30D | >16.667 | >16.67 | >16.67 | >16.667 | >16.667 | >16.67 | >16.667 | >16.67 |
| F105 | >16.667 | >16.67 | >16.67 | >16.667 | >16.667 | >16.67 | >16.667 | >16.67 |
| VRC01 | 3.31 | 1.164 | >6.67 | 0.28 | >6.667 | 3.11 | 1.89 | >6.67 |

### Table S7, continued

V2 apex

| | Tier 2 | Tier 2 | Tier 2 | Tier 2 | Tier 2 | Tier 2 | Tier 2 | Tier 2 | Tier 2 | Tier 3 |
|---|---|---|---|---|---|---|---|---|---|---|
| | H703_2117_110_RE_e2A10s | H703_2149_060_RE_eB10s | H703_2018_240_RE_e6A1s | H703_2805_080Es | H703_1675_G613s | H703_1313_040s | H704_0128_220_RE_pb001_s | H704_1775_030cN_SynGtoA_567 | H703_2304_150_RE_cs | H703_0646_051sN |
| Specimen ID | Clade: C | Clade: C | Clade: C | Clade: C | Clade: C | Clade: C | Clade: F1/B | Clade: F1 | Clade C | Clade C |
| SA-C10 | >1666.67 | 1417.451 | >1666.67 | >1666.67 | 173.596 | 1623 | >1666.67 | >1666.67 | >1666.67 | >1666.67 |
| SA-C48 | >1666.67 | >1666.67 | 1017.621 | >1666.67 | >1666.67 | 421 | >1666.67 | 478 | >1666.67 | >1666.67 |
| SA-C72 | 1381 | 605.54 | >1666.67 | 629 | 1090.774 | 461 | >1666.67 | 748 | >1666.67 | >1666.67 |
| SA-C74 | 489 | 894.9 | 1256.247 | 1028 | 563.956 | >1666.67 | >1666.67 | 890 | 1403 | >1666.67 |
| SA-C90 | 1181 | 486.581 | >1666.67 | 822 | 1433.617 | 1653 | 1508 | >1666.67 | 1192 | >1666.67 |
| 2219 | >16.667 | >16.67 | >16.67 | >16.667 | >16.67 | >16.667 | >16.667 | >16.667 | >16.667 | >16.67 |
| 2557 | >16.667 | >16.67 | >16.67 | >16.667 | >16.67 | >16.667 | >16.667 | >16.667 | >16.667 | >16.67 |
| 3074 | >16.667 | >16.67 | >16.67 | >16.667 | >16.67 | >16.667 | >16.667 | >16.667 | >16.667 | >16.67 |
| 3869 | >16.667 | >16.67 | >16.67 | >16.667 | >16.67 | >16.667 | >16.667 | >16.667 | >16.667 | >16.67 |
| 447-52D | >16.667 | >16.67 | >16.67 | >16.667 | >16.67 | >16.667 | >16.667 | >16.667 | >16.667 | >16.67 |
| 838-12D | >16.667 | >16.67 | >16.67 | >16.667 | >16.67 | >16.667 | >16.667 | >16.667 | >16.667 | >16.67 |
| 654-30D | >16.667 | >16.67 | >16.67 | >16.667 | >16.67 | >16.667 | >16.667 | >16.667 | >16.667 | >16.67 |
| 1008-30D | >16.667 | >16.67 | >16.67 | >16.667 | >16.67 | >16.667 | >16.667 | >16.667 | >16.667 | >16.67 |
| 729-30D | >16.667 | >16.67 | >16.67 | >16.667 | >16.67 | >16.667 | >16.667 | >16.667 | >16.667 | >16.67 |
| F105 | >16.667 | >16.67 | >16.67 | >16.667 | >16.67 | >16.667 | >16.667 | >16.667 | >16.667 | >16.67 |
| VRC01 | 0.3 | 0.555 | 2.87 | >6.667 | 0.549 | 1.25 | 0.2 | 0.44 | 1.89 | >6.67 |

CD4bs

| | Tier 2 | Tier 2 | Tier 2 | Tier 2 | Tier 2 | Tier ?* | Tier 2 | Tier 2 | Tier 2** |
|---|---|---|---|---|---|---|---|---|---|
| | H704_0855_080_EsN | H704_0128_220_RE_pb001_s | H703_0472_030s | H704_1180_070EsN | H704_1535_030sN | H703_0537_011s_4H1* | H704_1775_030cN_SynGtoA 567 | H703_2805_080Es | H704_1783_150_RE_cs |
| Specimen ID | Clade B | Clade F1/B | Clade C | Clade A1/B | Clade B | Clade C | Clade F1 | Clade C | Clade B |
| SA-C10 | 789 | >1666.67 | >1666.67 | 588 | 468 | | >1666.67 | >1666.67 | >1666.67 |
| SA-C48 | 919 | >1666.67 | >1666.67 | 587 | 136 | | 478 | >1666.67 | >1666.67 |
| SA-C72 | >1666.67 | >1666.67 | 534 | 821 | 168 | | 748 | 629 | >1666.67 |
| SA-C74 | >1666.67 | >1666.67 | >1666.67 | 783 | 84 | | 890 | 1028 | >1666.67 |
| SA-C90 | 1150 | 1508 | >1666.67 | >1666.67 | 782 | | >1666.67 | 822 | 1277 |
| 2219 | >16.667 | >16.667 | >16.667 | >16.667 | >16.667 | | >16.667 | >16.667 | >16.67 |
| 2557 | >16.667 | >16.667 | >16.667 | >16.667 | >16.667 | | >16.667 | >16.667 | >16.67 |
| 3074 | >16.667 | >16.667 | >16.667 | >16.667 | 13 | | >16.667 | >16.667 | >16.67 |
| 3869 | >16.667 | >16.667 | >16.667 | >16.667 | >16.667 | | >16.667 | >16.667 | >16.67 |
| 447-52D | >16.667 | >16.667 | >16.667 | >16.667 | >16.667 | | >16.667 | >16.667 | >16.67 |
| 838-12D | >16.667 | >16.667 | >16.667 | >16.667 | >16.667 | | >16.667 | >16.667 | >16.67 |
| 654-30D | >16.667 | >16.667 | >16.667 | >16.667 | >16.667 | | >16.667 | >16.667 | >16.67 |
| 1008-30D | >16.667 | >16.667 | >16.667 | >16.667 | >16.667 | | >16.667 | >16.667 | >16.67 |
| 729-30D | >16.667 | >16.667 | >16.667 | >16.667 | >16.667 | | >16.667 | >16.667 | >16.67 |
| F105 | >16.667 | >16.667 | >16.667 | >16.667 | >16.667 | | >16.667 | >16.667 | >16.67 |
| VRC01 | 0.09 | 0.2 | 0.308 | 0.28 | 0.21 | | 0.44 | >6.667 | 0.22 |

| Tier | IC50 Range |
|---|---|
| Tier 1A | <64 |
| Tier 1B | 64- 310 |
| Tier 2 | 296-1603 |
| Tier 3 | >1603 |

**Table S7. Establishing the Tier status by using 5 standardized sera and a set of monoclonals that are typically only exposed on Tier 1 viruses in open conformation.** The standardized sera are called SA-C10, SA-C48, SA-C72, SA-C74, and SA-C90. Members of the SHEP-T2 panel were sometimes included in the sensitive panels for a particular class. *The two viruses with a single asterisk did not grow well, so were replaced with the viruses with the marked with ** so reagent stocks could be generated and shared.
